# Comparative analysis of Lysine and Arginine biosynthesis pathway in *Deinococcus* genomes

**DOI:** 10.1101/2021.12.28.474397

**Authors:** Sankar Mahesh, Deepa Sethi, Richa Priyadarshini, Ragothaman M. Yennamalli

## Abstract

The members of the Deinococcaceae family have the ability to survive extreme environmental conditions. Deinococcus species have a complex cell envelope composed of L-ornithine containing peptidoglycan. Anabolism of L-ornithine is intrinsically linked to L-lysine and L-arginine biosynthetic pathways. To understand these two pathways, we analyzed the L-lysine and L-arginine pathways using 23 Deinococcus genomes, including *D. indicus*. We used BLAST-P based ortholog identification using *D. radiodurans*’ genes as the query. We identified some BLAST-P hits that shared the same functional annotation. We analyzed three (class I aminotransferase, acetyl-lysine deacetylase, and acetyl glutamate/acetyl aminoadipate kinase) from L-lysine biosynthesis pathway and three (bifunctional ornithine acetyltransferase or N-acetyl glutamate synthase protein, nitric oxide synthase-like protein, and Acetyl-lysine deacetylase) from L-arginine biosynthesis pathway. Two proteins showed certain structural variations. Specifically, [LysW]-lysine hydrolase protein’s sequence and structure level changes indicated changes in oligomeric conformation, which could likely be a result of divergent evolution. And, bifunctional ornithine acetyltransferase or N-acetyl glutamate synthase had its active site pocket positions shifted at the structural level and we hypothesize that it may not perform at the optimal level. Thus, we were able to compare and contrast different Deinococcus species indicating some genes occurring because of divergent evolution.

## Introduction

As per NCBI taxonomy database, more than 90 distinct species of *Deinococcus* genus have been isolated across the world from diverse sources such as Antarctic soil, deserts, hot springs, air, radiation sites, heavy-metal contaminated soil, sewage, plant rhizosphere, and human stomach [Sayers et al, 2010]. The members of the *Deinococcaceae* family are aerobic, non-motile, chemo-organotrophs that display extreme resistance to UV/gamma radiation, as well as, desiccation. *Deinococcus radiodurans* is the most extensively studied bacterium in this genus due to its remarkable ability to resist up to 10 kGy gamma radiation [Rainey et al, 2006]. *Deinococcaceae* family members possess a complex cell envelope that comprises a thick peptidoglycan layer, an outer membrane-like lipid layer, and a paracrystalline S-layer. While most members exhibit gram-positive stain reaction, a few members are gram-negative such as *D. ficus* [Lai et al, 2006]; *D. grandis* [Oyaizu et al, 1987]; *D. indicus* [Suresh et al, 2004], *D. depolymerans* [Asker et al, 2011], *D. aquaticus and D. caeni* [Im et al, 2008]. *Deinococcus* is phylogenetically closely related to extreme thermophiles of the genus Thermus and both genera exhibit L-ornithine-glycine in their peptidoglycan [Oyaizu et al, 1987; Embley et al, 1987; Quintela et al, 1999].

Lysine biosynthesis in bacteria and higher plants proceeds via the diaminopimelate pathway (DAP), where aspartate acts as the primary substrate (Figure 1). This pathway produces meso-diaminopimelate which is an immediate precursor of lysine and is also present in the bacterial cell wall. On the other hand, the α-aminoadipate pathway (AAA) is employed by some algae, fungi, and euglenoids. The AAA pathway uses α-ketoglutarate and acetyl coenzyme A for lysine synthesis, whereα-aminoadipic acid is formed as an intermediate in the AAA pathway. However, it has been shown now that some bacterial and archeal species such *as T. thermophilus, D. radiodurans, Pyrococcus horikoshii* lack the genes for the DAP pathway and use a variant of AAA pathway for lysine biosynthesis [Lombo et al, 2004; Kosuge et al, 1998].

**Figure 1:**
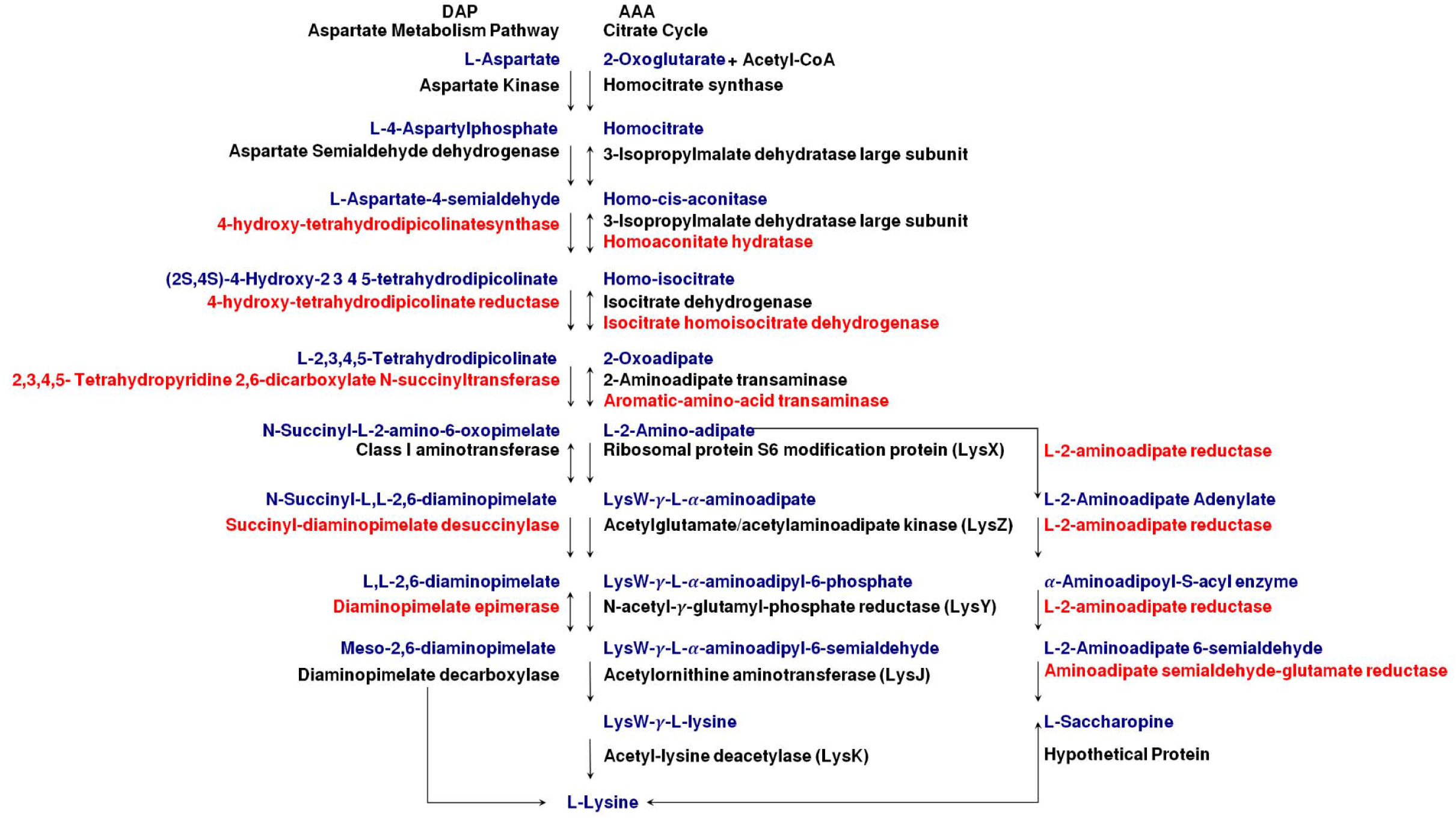
General schematic diagram of Lysine Pathway with *Deinococcus radiodurans* as the reference pathway. The genes highlighted in red are absent in *D. radiodurans* as per the KEGG pathway database. In cases where both red and black are present, it is understood that the protein in black compensates for the protein in red.

The DAP pathway begins with the phosphorylation of L-aspartate by aspartate kinase to generate L-aspartyl-4-phosphate. The enzyme aspartate semialdehyde dehydrogenase catalyzes the NADPH-dependent conversion of L-aspartyl-4-phosphate to L-aspartate-semialdehyde. This reaction is followed by a condensation step where 4-Hydroxy-tetrahydrodipicolinate synthase catalyzes the formation of (2S, 4*S*)-4-hydroxy-2,3,4,5-tetrahydro-(2*S*)-dipicolinate from pyruvate and L-aspartate semialdehyde. 4-hydroxy-tetrahydrodipicolinate reductase catalyzes the formation of L-2,3,4,5-Tetrahydrodipicolinate (THDPA). The enzyme tetrahydrodipicolinate succinylase then catalyzes the formation of N-succinyl-2-amino-6-oxopimelate from THDPA and succinyl-CoA. Further, enzyme N-succinyldiaminopimelate aminotransferase yields N-succinyl-L,L-2,6-diaminopimelate from N-succinyl-2-amino-6-ketopimelate. N-succinyl-L-diaminopimelate desuccinylase converts succinyl-diaminopimelate to L,L-2,6-Diaminopimelate through a deacylation step. Diaminopimelate epimerase (DapF) catalyzes the stereoinversion of L,L-diaminopimelate (LL-DAP) to *meso*-diaminopimelate. Lastly, a decarboxylation step forms lysine from meso-diaminopimelate. This reaction is catalyzed by Diaminopimelate decarboxylase (LysA) [Caspi et al, 2018]. The genes for lysine biosynthesis through the AAA pathway are clustered in *T. Thermophilus* and scattered on the genome in *D. radiodurans* [Nishida, 2001]. It has to be noted that in *Deinococcus*, enzymes involved in the AAA pathway for lysine biosynthesis are also involved in arginine biosynthesis [Nishida et al, 1999]. The arginine biosynthetic pathway generates ornithine as a by-product which is subsequently used in the peptidoglycan formation.

The AAA pathway can be divided into two parts. In the first part, formation of AAA from 2-oxoglutarate the enzymes involved are homocitrate synthase, (homo) aconitase, homoisocitrate dehydrogenase, and AAA aminotransferase. The second part of the lysine biosynthetic pathway, i.e., the conversion from AAA to lysine, is mediated by five enzymes: LysX, LysZ, LysY, LysJ, and LysK. Another protein called LysW functions as an amino group carrier protein [Yoshida et al, 2016]. LysX catalyzes the first reaction of the latter part by modifying the α-amino group of AAA with the γ-carboxyl group of the C terminal glutamate residue of LysW. LysW-γ-AAA is then converted to LysW-γ-lysine through a series of phosphorylation, reduction, and amination steps. Finally, LysK releases lysine from LysW-γ-lysine due to its carboxypeptidase activity. The presence of acidic amino acid residues in LysW for electrostatic interactions with each enzyme makes it an efficient carrier protein.

The similarity between reactions of the latter parts of the lysine AAA pathway and glutamate to ornithine conversion in bacterial arginine biosynthesis (Figure 2) suggests a common evolutionary origin [Ouchi et al, 2013; Yoshida et al, 2016]. Formation of ornithine by the variant L-arginine biosynthesis IV has been described in the archaebacterium *Sulfolobus acidocaldarius, Thermobaculum, Thermoproteus, Deinococcus*, etc. The enzyme encoded by the lysK gene removes the carrier protein and L-arginine is synthesized from L-ornithine via the reaction intermediates L-citrulline and L-arginino-succinate [Caspi et al, 2018].

**Figure 2:**
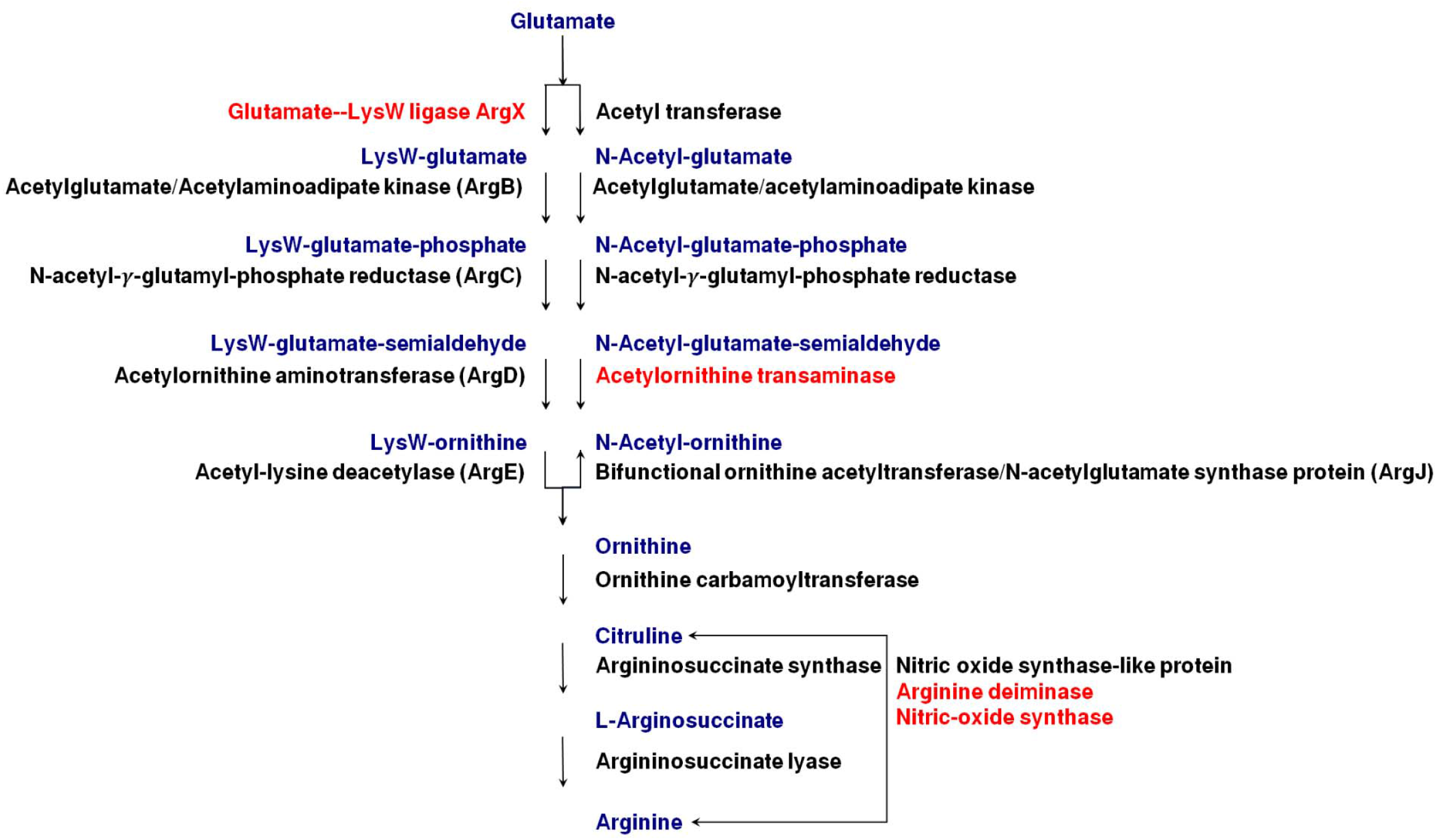
General schematic diagram of Arginine Pathway with *Deinococcus radiodurans* as the reference pathway. The genes highlighted in red are absent in *D. radiodurans* as per the KEGG pathway database. In cases where both red and black are present, it is understood that the protein in black compensates for the protein in red.

Recently our team isolated an arsenic tolerant bacterium, *Deinococcus indicus* strain DR1 from Dadri wetlands situated in North India [Chauhan et al, 2017]. Our studies also revealed the presence of L-ornithine in the cell wall of *D. indicus* [Chauhan et al, 2019], even though it is not a lysine auxotroph. Ornithine is a product in the arginine metabolic pathway and there are reports of an evolutionary link between arginine and lysine pathway. Details regarding the L-lysine and L-arginine metabolism in *Deinococcus* species are not well understood. Here, we surveyed the lysine and arginine biosynthesis pathway in 22 *Deinococcus* genomes and have made a comparative study of some of the unique differences that are present in some of these genomes.

## Materials and Methods

### Data collection and analysis

To investigate the presence or absence of all three sub-groups of Lysine biosynthesis in the *Deinococcus* genus, basic local alignment search tool-protein (BLAST-P) and basic local alignment search tool-nucleotide (BLAST-N) [Altschul et al, 1990; Camacho et al, 2009] search was performed using protein and nucleotide sequence of genes from *D. radiodurans* as the query. Genes reported to be involved in the Lysine biosynthesis pathway were extracted from the KEGG database [Kanehisa and Goto, 2000] and the sequences were downloaded from the GenBank database [Dennis et al, 2008]. In the case of arginine biosynthesis, protein and nucleotide sequences of genes involved in its synthesis as shown by the KEGG database [Kanehisa and Goto, 2000] were downloaded from GenBank database [Dennis et al, 2008] for *D*.*radiodurans* only. Genome assembly accession IDs of organisms used for BLAST-P is as follows: *D. actinosclerus* (GCF_001507665.1), *D. apachensis* (GCF_000381345.1), *D. aquatilis* (GCF_000378445.1), *D. deserti* (GCF_000020685.1), *D. ficus* (*G*CF_000430865.1), *D. frigens* (GCF_000701425.1), *D. geothermalis* (GCF_000196275.1), *D. gobiensis* (GCF_000252445.1), *D. grandis* (GCF_001485435.1), *D. hopiensis* (GCF_900176165.1), *D. indicus* (GCF_002198095.1), *D. maricopensis* (GCF_000186385.1), *D. marmoris* (GCF_000701405.1), *D. misasensis* (GCF_000745915.1), *D. murrayi* (GCF_000482805.1), *D. peraridilitoris* (GCF_000317835.1), *D. phoenicis* (GCF_000599865.1), *D. pimensis* (GCF_000519345.1), *D. proteolyticus* (GCF_000190555.1), *D. puniceus* (GCF_001644565.1), *D. radiodurans* (GCF_000008565.1), *D. reticulitermitis* (GCF_900109185.1), *D. soli* (GCF_001007995.1), *D. swuensis* (GCF_000800395.1), *D. wulumuqiensis* (GCF_000478785.1). Protein sequences from Genbank were downloaded for *D*.*radiodurans* and using these sequences as the query for BLAST-P against the genomes of other organisms (using their genome assembly ID mentioned above) we obtained the proteins involved in other organisms in this family along with BLAST statistics. (Supplementary Table 1)

### Model generation using Rosetta

Among the BLAST-P hits, those protein sequences having <60% sequence identity and <60% query coverage were selected for further analysis. Those that shared the functional annotation were analysed to identify the local and global differences. For this, we used a comparative modeling approach using Robetta [Kim et al 2004], after ascertaining that the experimentally derived structure does not exist. Comparative modeling was performed for a total of six sequences, three in the Lysine biosynthesis pathway and three in the Arginine biosynthesis pathway. Also, these sequences had homologs in PDB indicating that high-quality models can be generated using comparative modeling. For each sequence, partial threads were superimposed before a hybrid sampling approach involving 1000 models. The resultant five models are given a confidence score, which ranges between 0.0 to 1.0. Specifically, a confidence score of 0.0 is indicative of a bad model and 1.0 is a perfect model [Kimet al, 2004; Song Y et al, 2013; Chauhan et al, 2019]. The five models were structurally aligned to check their general agreement among them. If the RMSD is <0.5, then the first model was selected for further analysis. Otherwise, the THESEUS tool was implemented to identify the median structure among the five models [Douglas and Deborah, 2006; Teotia D et al, 2019].

## Results and Discussion

### Lysine biosynthesis pathway

Lysine biosynthesis pathway is divided into three sub-groups, among which one is mediated by the LysW gene (called LysW-mediated pathway) and the other two sub-groups are alpha-aminoadipate pathway (AAA) and diaminopimelate pathway (DAP) (Figure 1). We were curious to find out whether the three sub-groups of the Lysine biosynthesis pathway are present in *D. indicus* or not. Among the three, the AAA pathway is considered as being firmly a eukaryotic route and has been recently identified in deep branch bacteria (*Thermophilus* and *Deinococcus*) [Miyazaki et al, 2002; Lombo et al, 2004]. DAP pathway holds its importance as it generates diaminopimelate (DAP), a building block of peptidoglycan in many bacterial cell walls. Hence, a computational analysis was performed (BLAST-N and BLAST-P) against the whole genome and proteome of *D. indicus*. However, for further analysis, we relied on BLAST-P results as protein sequences are more conserved in nature than nucleotide sequences.

Looking at the BLAST-P results, where protein sequences from *D. radiodurans* were used as the query, the ten genes, involved in the LysW-mediated pathway of converting 2-oxoglutarate to 2-aminoadipate and further to lysine, are present in all the 22 *Deinococcus* genomes. However, *D. actinosclerus, D. deserti, D. maricopensis, D. misasensis, D. murrayi, D. peraridilitoris, D. pimensis*, and *D. wulumquensis* did not harbor high confidence orthologs for some of the genes i.e., the percentage identity of the hits was less than 60%, and the hits obtained from reverse BLAST-P show that the annotation is not the same as the query sequence (Figure 3).

**Figure 3:**
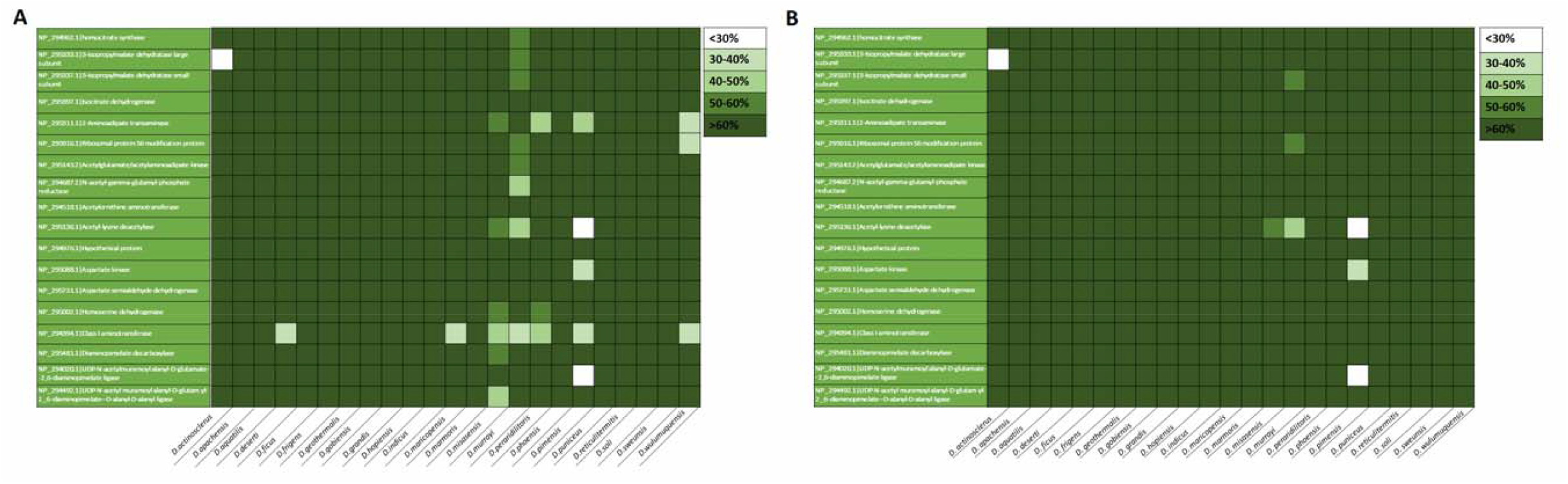
BLAST-P results of the Lysine biosynthesis pathway. The genes involved in lysine biosynthesis among *Deinococcus* species (in A) indicate that certain genes have low sequence identity with respect to *Deinococcus radiodurans* although they share the same functional annotation (in B).

Specifically, *D. actinosclerus* had one gene 3-isopropyl malate dehydratase large subunit (Acc. No: NP_295333.1) missing. Similarly, the gene Class I aminotransferase (Acc. No: NP_294394.1) was not present in *D. deserti* and *D. maricopensis*.

However, in *D. misasensis* six genes were missing, namely 2-aminoadipate transaminase (Acc. No: NP_295311.1), Acetyl-lysine deacetylase (Acc. No: NP_295136.1), Homoserine dehydrogenase (Acc. No: NP_295002.1), Class I aminotransferase (Acc. No: NP_294394.1), Diaminopimelate decarboxylase (Acc. No: NP_295481.1), and UDP-N-acetylmuramoylalanyl-D-glutamyl-2,6-diaminopimelate-D-alanyl-D-alanyl ligase (Acc. No: NP_294492.1).

On the other hand, *D. murrayi* had eight (the highest number) genes missing; namely Homocitrate synthase (Acc. No: NP_294962.1), 3-isopropylmalate dehydratase large subunit (Acc. No: NP_295333.1), 3-isopropylmalate dehydratase small subunit (Acc. No: NP_295337.1), Ribosomal protein S6 modification protein (Acc. No: NP_295916.1), Acetylglutamate/ Acetylaminoadipate kinase (Acc. No: NP_295143.2), N-acetyl-gamma-glutamyl-phosphate reductase (Acc. No: NP_294687.2), Acetyl-lysine deacetylase (Acc. No: NP_295136.1), and Class I aminotransferase (Acc. No: NP_294394.1).

Similarly, *D. peraridilitoris* had three genes missing, namely 2-aminoadipate transaminase (Acc. No: NP_295311.1), Homoserine dehydrogenase (Acc. No: NP_295002.1), and Class I aminotransferase (Acc. No: NP_294394.1).

D. *pimensis* had five genes missing, namely 2-aminoadipate transaminase (Acc. No: NP_295311.1), Acetyl-lysine deacetylase (Acc. No: NP_295136.1), Aspartate kinase (Acc. No: NP_295088.1), Class I aminotransferase (Acc. No: NP_294394.1) and UDP-N-acetyl muramoyl alanyl-D-glutamate-2,6-diaminopimelate ligase (Acc. No: NP_294020.1).

Finally, *D. wulumuquensis* had 3 genes missing, namely 2-aminoadipate transaminase (NP_295311.1), Ribosomal protein S6 modification protein (NP_295916.1), and Class I aminotransferase (NP_294394.1).

### Arginine biosynthesis pathway

Looking at the BLAST-P results, where protein sequences from *D. radiodurans* were used as the query, the ten genes involved in the LysW mediated pathway (from the conversion of 2-oxoglutarate to ornithine and further to arginine) are present in all the twenty-two *Deinococcus* genomes (Figure 4). However, *D*.*misasensis, D*.*murrayi, and D*.*pimensis* do not harbor satisfying orthologs for some of the genes i.e., the percentage identity of the hits was less than 60%, and the hits obtained from reverse BLAST-P show that the annotation is not the same as the query sequence.

**Figure 4:**
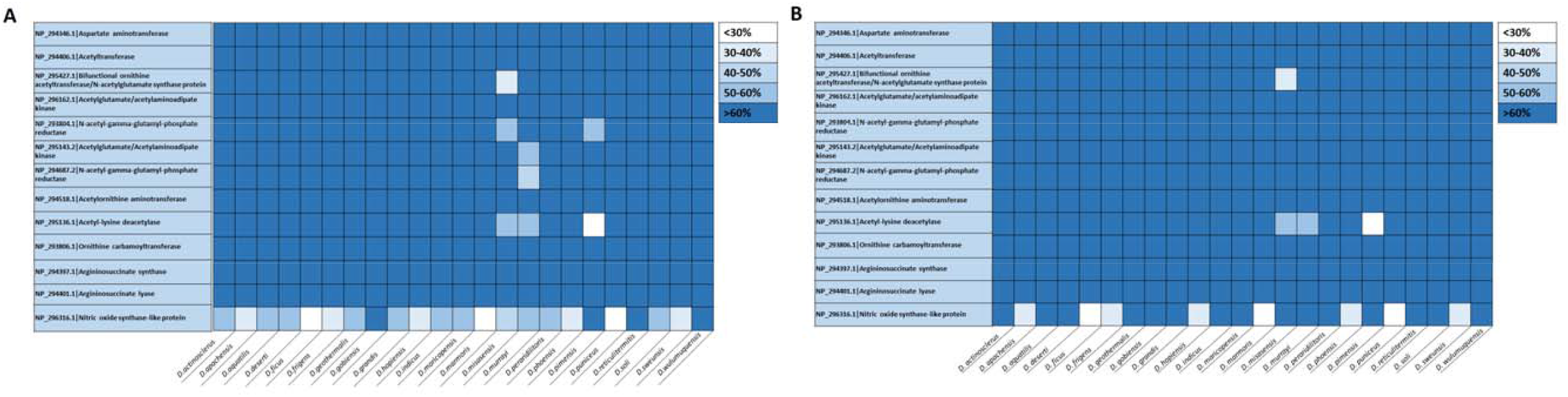
BLAST-P results of the Arginine biosynthesis pathway. The genes involved in arginine biosynthesis among *Deinococcus* species (in A) indicate that certain genes have low sequence identity with respect to *Deinococcus radiodurans* although they share the same functional annotation (in B).

Specifically, *D*.*misasensis* did not have orthologs for Bifunctional ornithine acetyltransferase/N-acetylglutamate synthase protein (Acc. No: NP_295427.1), N-acetyl-gamma-glutamyl-phosphate reductase (Acc. No: NP_293804.1), and Acetyl-lysine deacetylase (Acc. No: NP_295136.1). Similarly, *D*.*murrayi* did not have orthologs for Acetylglutamate/Acetylaminoadipate kinase (Acc. No: NP_295143.2), N-acetyl-gamma-glutamyl-phosphate reductase (Acc. No: NP_294687.2), and Acetyl-lysine deacetylase (Acc. No: NP_295136.1). Similarly, *D*.*pimensis* did not have orthologs for N-acetyl-gamma-glutamyl-phosphate reductase(Acc. No: NP_293804.1) and Acetyl-lysine deacetylase (Acc. No: NP_295136.1).

In addition to the above, 19 *Deinococcus* species did not harbor a high confidence ortholog for Nitric oxide synthase-like protein (Acc. No: NP_296316.1). The nineteen species being *D. actinosclerus, D. apachensis, D. aquatilis, D. deserti, D. ficus, D. frigens, D. geothermalis, D. grandis, D. hopiensis, D. indicus, D. maricopensis, D. marmoris, D. misasensis, D. murrayi, D. peraridilitoris, D. phoensis, D. puniceus, D. soli*, and *D. sweunsis*

Comparing *Thermus thermophilus* with *Deinococcus radiodurans* we observe that both the species produce lysine through the AAA pathway and the initial four steps are shared by the two organisms. In addition to this, the genes lys Z, Y, J, and K are homologous to arg B, C, D, and E respectively. The reactions involving arg B, C, D, and E are expected to have arisen from the duplication and further evolutionary divergence of the corresponding ancestral specific genes [Fondi et al, 2007]. Additionally, three of the six enzymes chosen for structural analysis (Acetyl-lysine deacetylase (Lys K), Acetylglutamate/acetylaminoadipate kinase (Lys Z) and Acetyl-lysine deacetylase (Arg E) have been specifically mentioned in the literature in view of the evolutionary divergence following gene duplication in primitive metabolic networks.

### Comparative modeling of low sequence identity pairs

Among the BLAST-P hits, those that had <60% sequence identity and <60% query coverage and that shared the same functional annotation were chosen for further analysis. This can be seen from the difference between figures 3A and 3B for lysine biosynthesis and figures 4A and 4B for arginine biosynthesis. High-quality 3D models were generated for the selected proteins. For proteins to preserve the same function in the presence of perturbations during the course of evolution (for example mutations, deletions, etc.) it has been observed that the active site and substrate binding-residues are relatively immutable in comparison to other parts of the protein that offer structural support for optimal function [Branden CI, and Tooze J, 2012]. Also, changes in sequence and thereby in the structure are accommodated by subtly altering the intramolecular interactions in the protein [Branden CI, and Tooze J, 2012]. We hypothesized that in the case of the various Deniococcus species, divergent evolution played a role in altering the sequence-level information among the various enzymes involved in Lysine and Arginine pathway and at the same time, the alterations were accommodated to a large extent to preserve the function of each protein. To characterize these, we identified six pairs of proteins, based upon their sequence identity and functional annotation, (three from Lysine biosynthesis pathway and three from Arginine biosynthesis pathway) and 3D models were generated using Robetta [Kim et al, 2004] Among the five models provided by Robetta, the first model (hereafter labeled as Model1) was used for further analysis. The reason being that when structurally aligned to the first model, the RMSD obtained after alignment was <0.5 indicating that there is general agreement among all the five models. For those structures that had >0.5 RMSD after structural alignment, we used THESEUS (https://theobald.brandeis.edu/theseus/)[Douglas and Deborah, 2006] to identify the median structure among the five models.

For example, the proteins Class I aminotransferase (NP_294394.1) and Acetyl glutamate/acetylaminoadipate kinase (NP_295143.2) from *D. radiodurans*, and pyridoxal phosphate-dependent aminotransferase (WP_012693528.1), [LysW]-lysine hydrolase (WP_034341940.1), [LysW]-aminoadipate kinase (WP_051363596.1) from *D. deserti, D. misasensis* and *D. murrayi* respectively in the lysine biosynthesis pathway and the proteins nitric oxide synthase-like protein (NP_296316.1) from *D. radiodurans*, and bifunctional glutamate N-acetyltransferase/amino-acid acetyltransferase ArgJ (WP_034339798.1), nitric oxide synthase oxygenase (WP_051964790.1) from *D. misasensis* in the arginine biosynthesis pathway had <0.5 RMSD value after structural alignment and thus Model1 was designated as the representative structure for analysis. Similarly, the proteins acetyl-lysine deacetylase (NP_295136.1) from *D. radiodurans* in the lysine biosynthesis pathway and the proteins bifunctional ornithine acetyltransferase/ N-acetyl glutamate synthase protein (NP_295427.1), acetyl-lysine deacetylase(NP_295136.1) from *D. radiodurans*, and [LysW]-lysine hydrolase(WP_027461251.1) from *D. murrayi* in the arginine biosynthesis pathway had >0.5 RMSD value after structural alignment and the following models were identified as representative structures. Model 2 was identified as the median structure of the protein acetyl-lysine deacetylase (NP_295136.1) and [LysW]-lysine hydrolase (WP_027461251.1). Model 1 was identified as the median structure of bifunctional ornithine acetyltransferase/N-acetyl glutamate synthase protein (NP_295427.1)

All the twelve structures thus selected have a high confidence score of 0.86-1 (Supplementary Table 1). It has been previously reported that a confidence score of 0.8 and above indicates a high- to very high-quality models generated and these are comparable to the structures solved using experimental methods [Kim et al, 2004; Chauhan et al, 2019; Song Y et al, 2013].

### Specific analysis of proteins that share low sequence similarity

#### Class I aminotransferase and pyridoxal phosphate-dependent aminotransferase of the lysine biosynthesis pathway function as dimer

Aminotransferases are enzymes that are involved in transferring an amine group. On comparing the modeled structure of class I aminotransferase (NP_294394.1) from *D. radiodurans R1* and pyridoxal phosphate-dependent aminotransferase (WP_012693528.1) from *D. deserti* we observed the RMSD value to be 1.569 (Figure 5A). This difference can be attributed to six regions of structural non-equivalence starting with Met1 to Asp35 at the N-terminal followed by Phe84 to Asp91, Val141 to Asp150, Ser264 to Lys290, Ser332 to Ile344, and Gle373 to Pro388 at the C-terminal. The active sites Lys226 in *D. radiodurans* and Lys237 in *D. deserti* are structurally conserved. A majority of the regions listed above form secondary structures of structural importance. These include Gln9 to Ser13, Thr19 to Arg31, Phe84 to Asn87, Ser272 to Lys290, Glu337 to Ile344, and Glu373 to Pro388 forming α-helices while the region Val141 to Val145 forms a β-strand. These enzymes require a coenzyme named pyridoxal phosphate to function. This binds to the lysine residue at the active site and creates the formation of a Schiff Base or aldimine. Since the active site is positioned at the dimer interface like any other enzyme in the aminotransferase family, it can be said that both the subunits of the homodimer contribute to the active site architecture [Weyand et al, 2007]. In conclusion, the putative structure of the protein pyridoxal phosphate-dependent aminotransferase (WP_012693528.1) from *D. deserti*, reveals that it has conserved its active site to retain its functionality, and functions as a dimer.

**Figure 5:**
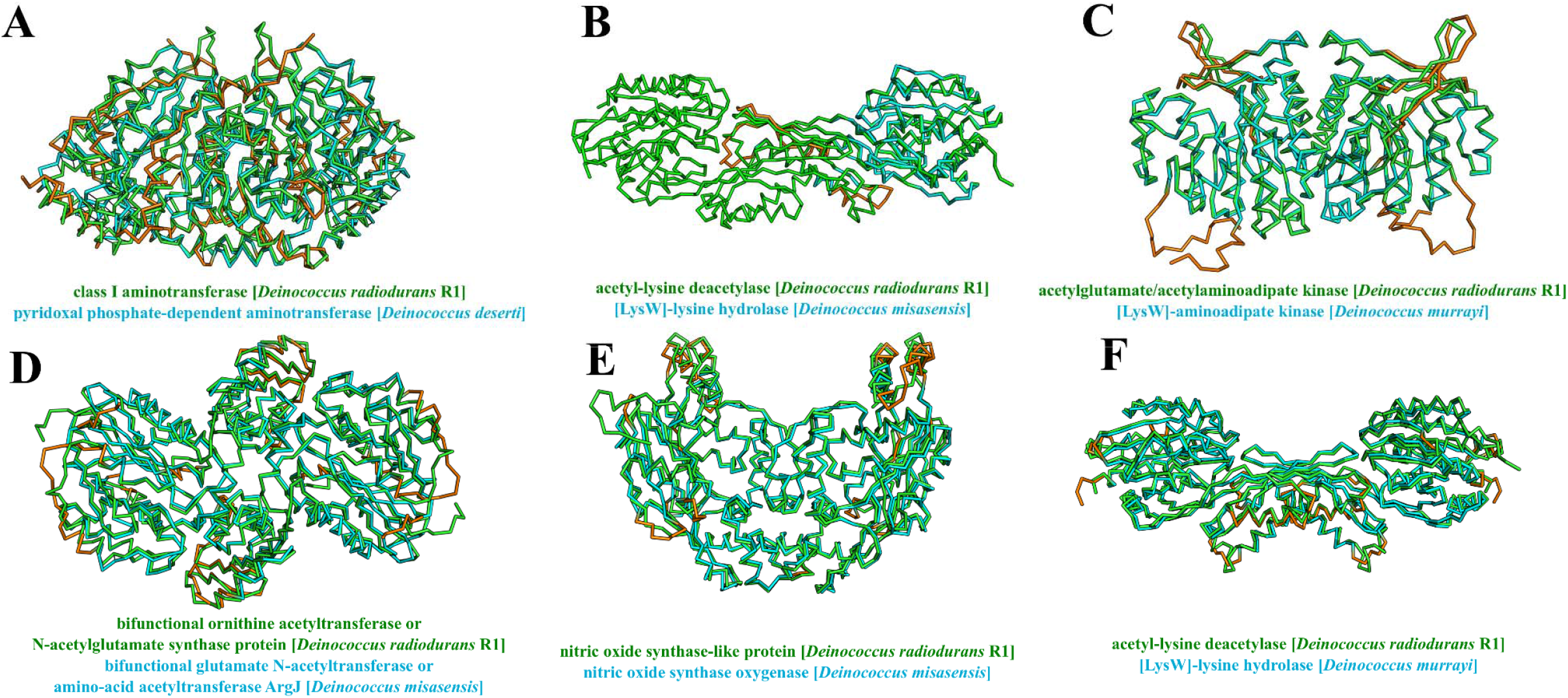
Specific case analysis. Superposed structures of *Deinococcus radiodurans* (shown in green colored C^α^ trace) with other species. The structurally non-equivalent parts are colored in orange for three enzymes involved in lysine biosynthesis pathway: A) Class I aminotransferase, B) Acetyl-lysine deacetylase, C) Acetylglutamate/acetylaminoadipate kinase; and three enzymes involved in arginine biosynthesis pathway: D) Bifunctional ornithine acetyltransferase, E) Nitric oxide synthase-like protein, and F) Acetyl-lysine deacetylase.

#### Acetyl-lysine deacetylase and LysW-lysine hydrolase of the lysine biosynthesis pathway have active site preserved

Comparing the modeled structure of the proteins Acetyl-lysine deacetylase (NP_295136.1) from *D. radiodurans* and [LysW]-lysine hydrolase (WP_034341940.1) from *D. misasensis* (Figure 5B) we observed the active sites Asp71 and Glu127 of Acetyl-lysine deacetylase to be structurally conserved at positions 64 and 119 of [LysW]-lysine hydrolase. In addition to this the metal-binding sites His69, Asp94, Glu128, Glu151, His334 of Acetyl-lysine deacetylase were conserved at positions His62, Asp86, Glu120, Glu143, His324 of [LysW]-lysine hydrolase. On aligning the protein sequences, the RMSD computed was 0.383 and this can be attributed to the misaligned turns and loops. Interestingly, we noticed that the template proteins used in comparative modeling exist in different oligomeric forms namely dimer, tetramer, and octamer. But the modeled structure of [LysW]-lysine hydrolase of *D. misasensis* indicates that it functions as a monomer only as the structurally non-equivalent positions (colored in orange in Figure 5B) are the regions in *D. radiodurans* that are involved in dimer formation. We hypothesize that this may be a result of divergent evolution. Chains in the protein sequences that possess the active sites and are involved in the functionality of the protein would be conserved through the course of evolution. In conclusion, the protein [LysW]-lysine hydrolase of *D. misasensis* has conserved active site residues to preserve its functionality in the lysine biosynthesis pathway and it is expected to function as a monomer in *D. misasensis*.

#### Acetylglutamate/acetylaminoadipate kinase and [LysW]-aminoadipate kinase of the lysine biosynthesis pathway have structural conservation

The next pair of proteins that were analyzed were Acetylaminoadipate kinase (NP_295143.2) from *D. radiodurans R1* and [LysW]-aminoadipate kinase (WP_051363596.1) from *D. murrayi* (Figure 5C) that shared a sequence identity of 55.64%. The amino acid sequences of these proteins revealed four regions of structural non-equivalence each consisting of 10 -20 residues. These are Thr2 - Pro10, Lys62 - Thr71, Arg120 - Asp139, and Gly276 - Ala289. On aligning the modeled protein structures, the RMSD computed was 0.540, attributed to the non-aligned regions consisting of either β turns or random coils. The ortholog of *D. radiodurans* i.e, [LysW]-aminoadipate kinase from *D. murrayi* had active site conserved that are involved in stabilization of the transition state of LysW-gamma-L-alpha-aminoadipate - ATP at positions Lys14 and Lys238. Interestingly, one among the two substrate-binding site residues has been replaced by residue with similar physiochemical properties. Specifically, isoleucine at 176 is replaced by leucine, as both are hydrophobic and the neighboring residue (Ala175 in *D. radiodurans*) has co-evolved by being replaced with leucine (Ala175Leu) to achieve global stability. This is thought of as a mechanism to compensate for the loss of function. Thus, the putative structure of [LysW]-aminoadipate kinase from *D. murrayi* revealed that despite a single-point mutation it can function as a kinase.

#### Bifunctional ornithine acetyltransferase or N-acetylglutamate synthase protein and bifunctional glutamate N-acetyltransferase or amino-acid acetyltransferase ArgJ from arginine biosynthesis pathway active site residue is different

Bifunctional ornithine acetyltransferase or N-acetyl glutamate synthase is a protein involved in the catalysis of two reactions in the arginine biosynthesis pathway. Superposing the modeled protein structures of bifunctional ornithine acetyltransferase or N-acetyl glutamate synthase protein (NP_295427.1) from *D. radiodurans* and its ortholog bifunctional glutamate N-acetyltransferase or amino-acid acetyltransferase (WP_034339798.1) from *D. misasensis* showed an RMSD of 0.870, indicating structural conservation (Figure 5D). The minor difference starts from the N-terminal region, Met1 to Gln6 followed by Val58 to Ile62, Arg121 to Asp138, Pro174 to Met179, and Lys309 to Asp343. Although the structurally nonequivalent regions in the N-terminal are loops, a few functionally important secondary structures (α-helix and β-sheets) in the above-mentioned regions were observed. Specifically, the sequence region Val58 to Asn60, Arg121 to Gly128, Leu136 to Asp138, and Met176 to Thr178 exhibit α-helices while Gln323 to Asp333 exhibits a β-sheet fold Since both enzymes belong to the ornithine acetyltransferase family, their catalytic mechanism involves a nucleophilic attack on the carbonyl group of amide bond using the side chain of an N-terminal threonine, serine or cysteine [Elkins et al, 2005]. Also, this reaction takes place through the formation of an oxyanion hole and Thr108 and Gly109 that are involved in the stabilization of this oxyanion were conserved [5]. This enzyme has six substrate-binding residues Thr145, Lys167, Thr178, Glu257, Asn382, and Thr387 and these were also conserved. Despite all the above observations being in favor of the enzyme’s functionality, the active site of both the enzymes (Thr184 in *D. radiodurans* and *D. misasensis)* are not structurally equivalent. On further investigation, we noticed these residues exhibit different rotamer configurations thereby altering the active site pocket. This indeed would prevent the enzyme from its expected functioning. Thus, the modeled structure of bifunctional glutamate N-acetyltransferase or amino-acid acetyltransferase from *D. misasensis* reveals that it might not possibly function as expected owing to the difference in the rotamer configuration of its active site.

#### Nitric oxide synthase from arginine biosynthesis pathway acts as a dimeric protein

The dimeric structure of *D. misasensis* nitric oxide synthase oxygenase (WP_051964790.1) when superposed with the dimeric structure of *D. radiodurans* nitric oxide synthase-like protein (NP_296316.1) showed an RMSD of 0.978 (Figure 5E). This can be attributed to four non-aligned regions in the protein sequence, each spanning 5 -20 amino acid residues. These are Ser2 - Pro7 Tyr17 - Phe36, Gly121 - Gly126, and Pro146 - Phe152. The structurally non-equivalent regions in the N-terminal of *D. misasensis* nitric oxide synthase oxygenase resulted in an α-helix and the rest of the structurally non-equivalent regions comprised of turns. The heme-binding site in both the proteins has been conserved at Cys63 and Cys58 respectively. Although most of the species classified under the genus *Deinococcus* lack the enzymes required to synthesize a cofactor called tetrahydrobiopterin (H_4_B), the H_4_B sites are conserved in this pair. It has been reported that the conversion of Arginine to Citrulline takes place in the presence of tetrahydrofolate [Pant et al, 2002], a reduced pteridine that compensates for H_4_B. The modeled structure of nitric oxide synthase oxygenase from *D. misasensis* revealed that it functions as a dimer similar to *D. radiodurans* and it catalyzes the reaction in the presence of THF [Pant et al, 2002].

#### Acetyl-lysine deacetylase and LysW-lysine hydrolase of the arginine biosynthesis pathway also acts as a dimeric protein

Superimposing the structures of LysW-L-lysine hydrolase (WP_027461251.1) of *D. murrayi* with LysW-L-ornithine hydrolase (NP_295136.1) of *D. radiodurans* the RMSD was 0.95 (Figure 5F). This is due to the absence of N-terminal residues in *D. radiodurans* Acetyl lysine deacetylase and structural differences in some secondary structure regions. Specifically, the residues that are structurally non-equivalent in *D. murrayi* are from Ala103 to Arg109 and from Gly241 to Ser258 Also, sequence-level differences in *D. murrayi* were in the non-secondary structure regions such as turns. The active sites Ala65, Glu120, and the metal-binding sites His63, Asp87, Glu121, Glu144, and His325 are conserved, thus functionally able to synthesize ornithine. Unlike the [LysW]-L-lysine hydrolase from *D. misasensis*, which has a similar sequence and structural identity to that of LysW-L-lysine hydrolase of *D. radiodurans* (Figure 5B), this protein is modeled as a dimer and its dimer interface seems to be conserved, as well. Thus, the modeled structure of LysW-L-lysine hydrolase from *D. murrayi* reveals that it possibly functions as a dimer with conserved active sites and metal-binding sites.

## Conclusions

The objective of our work was to derive insights from the genomes of various *Deinococcus* species concerning ornithine production. On analyzing the sequences and structures of various proteins in *Deinococcus* species, we were able to identify [LysW]-lysine hydrolase of *D. misasensis* functioning despite having a different global stoichiometry than its ortholog *Acetyl-lysine deacetylase* in *D. radiodurans*. Besides, we were able to predict that [LysW]-aminoadipate kinase (WP_051363596.1) from *D. murrayi* could function despite its single point mutation. Three other proteins have also shown sequence level and structure level difference, despite having the same functional annotation. Through this, we can understand how proteins can evolve to function differently further substantiating divergent evolution in proteins. These insights could lead to further classification of protein structures based on domains and have prospects in domain-based clustering of protein sequences. Our investigation of lysine and arginine biosynthetic routes also suggests that genes encoding enzymes with a broad substrate specificity are probably major players in primitive metabolic networks.

## Supporting information

Supplementary File with explanations

## Declarations

## Ethics approval and consent to participate

Not applicable

## Consent for publication

All authors have read the manuscript and have consented for publication.

## Authors’ contributions

RP and RMY conceived the study. SM and RMY performed data analysis. SM wrote the first draft of the manuscript. DS and RP wrote and edited the manuscript. All authors contributed toward the writing of the manuscript. All authors have approved the final article.

## Acknowledgments

RMY acknowledges Mr. Pulkit Anupam Srivastava the critical inputs and discussion. RMY and SM acknowledge SASTRA Deemed to be University for infrastructural support. DS is supported by a Doctoral fellowship from Shiv Nadar University. RP lab is supported by CSIR-EMR grant and start-up funds from Shiv Nadar University.

## Availability of data and materials

The data used in the manuscript are available in public repositories.

## Figure Legends

**Supplementary Table 1:**

The details of BLAST-P results of Arginine and Lysine biosynthesis pathway and details of the template used by Rosetta for performing comparative modeling, the confidence score, and the alignment of template sequences with the model sequences are given in the table.

