## Supplementary File with explanations for "Comparative analysis of Lysine and Arginine biosynthesis pathway in *Deinococcus* genomes"

**Supplementary Data.**

**Legend for BLAST-P hits.**
Rows marked in red imply that the percent identity of the hit was less than 60.
Rows marked in yellow imply that the Reverse BLAST hit didn't match the BLAST hit.
Rows marked in orange imply that more than one BLAST/Reverse BLAST hits were observed.
Rows marked in blue imply that BLAST hit matched with the Reverse BLAST hit.

**Lysine biosynthesis pathway.**

*D.actinosclerus*, *D.deserti*, and *D.maricopensis* returned one hit each that had <60% sequence identity with *D.radiodurans*.
*D.grandis*, *D.soli* returned two BLAST and Reverse BLAST hit for a single protein and the hits were inclusive.
*D.swuensis* returned two BLAST and 1 Reverse BLAST hit for a single protein whose hits were inclusive.
*D.misasensis* had six hits while D.murrayi had eight and D.pimensis had five and all these hits had less than 60% sequence identity with *D.radiodurans.*
Three hits were observed in *D.perariditoris* and *D.wulumuqiensis* and all the three had less than 60% sequence identity with *D.radiodurans.*

### LYSINE BIOSYNTHESIS BLAST-P HITS AND REVERSE BLAST-P HITS

*Deinococcus actinosclerus*

| Query ID | Subject ID | % Identity | Alignment length | Mismatches | Gap opens | Query start | Query end | Sequence start | Sequence end | E-value | Bit score | Query coverage |
|---|---|---|---|---|---|---|---|---|---|---|---|---|
| DR_1238/NP_294962.1\|homocitrate | WP_062158402.1 | 91.73 | 387 | 32 | 0 | 7 | 393 | 9 | 395 | 0 | 740 | 98 |
| DR_1610/NP_295333.1\|3-isopropylmalate | WP_082688890.1 | 26.45 | 121 | 76 | 5 | 186 | 296 | 80 | 197 | 0.2 | 29.6 | 26 |
| DR_1614/NP_295337.1\|3-isopropylmalate | WP_062158116.1 | 90.8 | 163 | 15 | 0 | 1 | 163 | 1 | 163 | 2.00E-107 | 305 | 92 |
| DR_1674/NP_295397.1\|isocitrate | WP_062157621.1 | 91.29 | 333 | 29 | 0 | 1 | 333 | 1 | 333 | 0 | 630 | 100 |
| DR_1588/NP_295311.1\|class | WP_062157117.1 | 76.54 | 405 | 89 | 2 | 16 | 419 | 6 | 405 | 0 | 622 | 96 |
| DR_2194/NP_295916.1\|ribosomal | WP_062158118.1 | 88.85 | 287 | 32 | 0 | 8 | 294 | 1 | 287 | 0 | 504 | 97 |
| DR_1420/NP_295143.2\|acetylglutamate/acetylaminoadipate | WP_062158240.1 | 88.39 | 267 | 31 | 0 | 1 | 267 | 1 | 267 | 2.00E-163 | 455 | 100 |
| DR_0963/NP_294687.2\|argC;N-acetyl-gamma-glutamyl-phosphate | WP_082689006.1 | 88.08 | 344 | 41 | 0 | 4 | 347 | 5 | 348 | 0 | 602 | 99 |
| DR_0794/NP_294518.1\|acetylornithine | WP_062158252.1 | 82.24 | 411 | 73 | 0 | 45 | 455 | 6 | 416 | 0 | 692 | 88 |
| DR_1413/NP_295136.1\|acetyl-lysine | WP_082689073.1 | 74.08 | 355 | 88 | 1 | 8 | 362 | 10 | 360 | 0 | 508 | 98 |
| DR_1252/NP_294976.1\|hypothetical | WP_062157445.1 | 90.37 | 405 | 39 | 0 | 1 | 405 | 1 | 405 | 0 | 778 | 100 |
| DR_1365/NP_295088.1\|aspartate | WP_062158132.1 | 83.84 | 458 | 74 | 0 | 10 | 467 | 5 | 462 | 0 | 728 | 98 |
| DR_2008/NP_295731.1\|aspartate-semialdehyde | WP_062158887.1 | 83.09 | 337 | 57 | 0 | 1 | 337 | 1 | 337 | 0 | 575 | 99 |
| DR_1278/NP_295002.1\|homoserine | WP_062157511.1 | 77.85 | 325 | 67 | 1 | 1 | 325 | 1 | 320 | 6.00E-178 | 496 | 100 |
| DR_0671/NP_294394.1\|class | WP_062156971.1 | 65.07 | 375 | 119 | 3 | 1 | 375 | 4 | 366 | 2.00E-145 | 418 | 98 |
| DR_1758/NP_295481.1\|diaminopimelate | WP_062158661.1 | 68.59 | 382 | 116 | 1 | 1 | 378 | 1 | 382 | 4.00E-172 | 486 | 99 |
| DR_0297/NP_294020.1\|UDP-N-acetylmuramoylalanyl-D-glutamate--2_6-diaminopimelate | WP_062159363.1 | 78.89 | 488 | 102 | 1 | 1 | 488 | 1 | 487 | 0 | 681 | 99 |
| DR_0768/NP_294492.1\|UDP-N-acetylmuramoylalanyl-D-glutamyl-2_6-diaminopimelate--D-alanyl-D-alanyl | WP_062158811.1 | 74.89 | 438 | 98 | 4 | 1 | 432 | 1 | 432 | 0 | 613 | 99 |

**REVERSE BLAST**

| | | | | | | | | | | | | |
|---|---|---|---|---|---|---|---|---|---|---|---|---|
| WP_062156971.1 | DR_0671/NP_294394.1 | 64.8 | 375 | 120 | 3 | 4 | 366 | 1 | 375 | 2.00E-155 | 444 | 97 |
| WP_062157117.1 | DR_1588/NP_295311.1 | 75.8 | 405 | 92 | 2 | 6 | 405 | 16 | 419 | 0 | 615 | 98 |
| WP_062157445.1 | DR_1252/NP_294976.1 | 90.37 | 405 | 39 | 0 | 1 | 405 | 1 | 405 | 0 | 778 | 100 |
| WP_062157511.1 | DR_1278/NP_295002.1 | 77.85 | 325 | 67 | 1 | 1 | 320 | 1 | 325 | 8.00E-178 | 496 | 100 |
| WP_062157621.1 | DR_1674/NP_295397.1 | 91.29 | 333 | 29 | 0 | 1 | 333 | 1 | 333 | 0 | 630 | 100 |
| WP_062158116.1 | DR_1614/NP_295337.1 | 90.8 | 163 | 15 | 0 | 1 | 163 | 1 | 163 | 2.00E-107 | 305 | 99 |
| WP_062158118.1 | DR_2194/NP_295916.1 | 88.85 | 287 | 32 | 0 | 1 | 287 | 8 | 294 | 0 | 504 | 100 |
| WP_062158132.1 | DR_1365/NP_295088.1 | 83.84 | 458 | 74 | 0 | 5 | 462 | 10 | 467 | 0 | 713 | 98 |
| WP_062158240.1 | DR_1420/NP_295143.2 | 88.39 | 267 | 31 | 0 | 1 | 267 | 1 | 267 | 3.00E-163 | 455 | 100 |
| WP_062158252.1 | DR_0794/NP_294518.1 | 82.24 | 411 | 73 | 0 | 6 | 416 | 45 | 455 | 0 | 713 | 99 |
| WP_062158402.1 | DR_1238/NP_294962.1 | 91.73 | 387 | 32 | 0 | 9 | 395 | 7 | 393 | 0 | 740 | 98 |
| WP_062158661.1 | DR_1758/NP_295481.1 | 68.59 | 382 | 116 | 1 | 1 | 382 | 1 | 378 | 0 | 513 | 99 |
| WP_062158811.1 | DR_0768/NP_294492.1 | 74.89 | 438 | 98 | 4 | 1 | 432 | 1 | 432 | 0 | 624 | 99 |
| WP_062158887.1 | DR_2008/NP_295731.1 | 83.09 | 337 | 57 | 0 | 1 | 337 | 1 | 337 | 0 | 575 | 99 |
| WP_062159363.1 | DR_0297/NP_294020.1 | 78.89 | 488 | 102 | 1 | 1 | 487 | 1 | 488 | 0 | 694 | 99 |
| WP_082688890.1 | NP_296139.1 | 92.13 | 216 | 17 | 0 | 1 | 216 | 1 | 216 | 1.00E-142 | 404 | 98 |
| WP_082689006.1 | DR_0963/NP_294687.2 | 88.08 | 344 | 41 | 0 | 5 | 348 | 4 | 347 | 0 | 601 | 99 |
| WP_082689073.1 | DR_1413/NP_295136.1 | 74.08 | 355 | 88 | 1 | 10 | 360 | 8 | 362 | 0 | 508 | 96 |

### *Deinococcus apachensis*

| Query ID | Subject ID | % Identity | Alignment length | Mismatches | Gap opens | Query start | Query end | Sequence start | Sequence end | E-value | Bit score | Query coverage |
|---|---|---|---|---|---|---|---|---|---|---|---|---|
| DR_1238/NP_294962.1\|homocitrate | WP_051086938.1 | 91.73 | 387 | 32 | 0 | 7 | 393 | 3 | 389 | 0 | 737 | 98 |
| DR_1610/NP_295333.1\|3-isopropylmalate | WP_026332290.1 | 89.78 | 411 | 42 | 0 | 17 | 427 | 7 | 417 | 0 | 717 | 95 |
| DR_1614/NP_295337.1\|3-isopropylmalate | WP_019585716.1 | 89.63 | 164 | 17 | 0 | 1 | 164 | 1 | 164 | 2.00E-107 | 305 | 93 |
| DR_1674/NP_295397.1\|isocitrate | WP_019588297.1 | 86.45 | 332 | 45 | 0 | 1 | 332 | 1 | 332 | 0 | 598 | 99 |
| DR_1588/NP_295311.1\|class | WP_019588759.1 | 72.4 | 413 | 105 | 4 | 12 | 420 | 1 | 408 | 0 | 565 | 97 |
| DR_2194/NP_295916.1\|ribosomal | WP_019585713.1 | 86.76 | 287 | 36 | 1 | 8 | 294 | 1 | 285 | 6.00E-180 | 499 | 97 |
| DR_1420/NP_295143.2\|acetylglutamate/acetylaminoadipate | WP_019584777.1 | 86.52 | 267 | 36 | 0 | 1 | 267 | 1 | 267 | 5.00E-159 | 444 | 100 |
| DR_0963/NP_294687.2\|argC;N-acetyl-gamma-glutamyl-phosphate | WP_026332162.1 | 85.47 | 344 | 50 | 0 | 4 | 347 | 5 | 348 | 0 | 617 | 99 |
| DR_0794/NP_294518.1\|acetylornithine | WP_019585874.1 | 83.13 | 409 | 69 | 0 | 40 | 448 | 8 | 416 | 0 | 721 | 88 |
| DR_1413/NP_295136.1\|acetyl-lysine | WP_019588678.1 | 73.58 | 352 | 87 | 2 | 10 | 361 | 11 | 356 | 4.00E-168 | 475 | 97 |
| DR_1252/NP_294976.1\|hypothetical | WP_026332415.1 | 83.21 | 405 | 68 | 0 | 1 | 405 | 1 | 405 | 0 | 725 | 100 |
| DR_1365/NP_295088.1\|aspartate | WP_019585687.1 | 81.25 | 464 | 87 | 0 | 10 | 473 | 5 | 468 | 0 | 705 | 98 |
| DR_2008/NP_295731.1\|aspartate-semialdehyde | WP_019586349.1 | 82.34 | 334 | 59 | 0 | 1 | 334 | 1 | 334 | 0 | 542 | 99 |
| DR_1278/NP_295002.1\|homoserine | WP_026332422.1 | 74.77 | 325 | 77 | 1 | 1 | 325 | 1 | 320 | 9.00E-170 | 476 | 100 |
| DR_0671/NP_294394.1\|class | WP_019585500.1 | 67.11 | 377 | 113 | 2 | 1 | 377 | 4 | 369 | 2.00E-177 | 500 | 99 |
| DR_1758/NP_295481.1\|diaminopimelate | WP_019585533.1 | 71.88 | 377 | 106 | 0 | 3 | 379 | 2 | 378 | 0 | 526 | 99 |
| DR_0297/NP_294020.1\|UDP-N-acetylmuramoylalanyl-D-glutamate--2_6-diaminopimelate | WP_019587223.1 | 78.37 | 490 | 103 | 1 | 1 | 490 | 1 | 487 | 0 | 719 | 100 |
| DR_0768/NP_294492.1\|UDP-N-acetylmuramoylalanyl-D-glutamyl-2_6-diaminopimelate--D-alanyl-D-alanyl | WP_019585961.1 | 71.89 | 434 | 110 | 4 | 1 | 432 | 1 | 424 | 0 | 605 | 99 |

**REVERSE BLAST**

| | | | | | | | | | | | | |
|---|---|---|---|---|---|---|---|---|---|---|---|---|
| WP_019584777.1 | DR_1420/NP_295143.2 | 86.52 | 267 | 36 | 0 | 1 | 267 | 1 | 267 | 9.00E-159 | 443 | 100 |
| WP_019585500.1 | DR_0671/NP_294394.1 | 66.58 | 377 | 115 | 2 | 4 | 369 | 1 | 377 | 2.00E-169 | 479 | 97 |
| WP_019585533.1 | DR_1758/NP_295481.1 | 71.88 | 377 | 106 | 0 | 2 | 378 | 3 | 379 | 0 | 543 | 99 |
| WP_019585687.1 | DR_1365/NP_295088.1 | 81.25 | 464 | 87 | 0 | 5 | 468 | 10 | 473 | 0 | 691 | 99 |
| WP_019585713.1 | DR_2194/NP_295916.1 | 86.76 | 287 | 36 | 1 | 1 | 285 | 8 | 294 | 2.00E-177 | 493 | 96 |
| WP_019585716.1 | DR_1614/NP_295337.1 | 89.63 | 164 | 17 | 0 | 1 | 164 | 1 | 164 | 2.00E-107 | 305 | 97 |
| WP_019585874.1 | DR_0794/NP_294518.1 | 83.13 | 409 | 69 | 0 | 8 | 416 | 40 | 448 | 0 | 721 | 98 |
| WP_019585961.1 | DR_0768/NP_294492.1 | 71.89 | 434 | 110 | 4 | 1 | 424 | 1 | 432 | 0 | 605 | 99 |
| WP_019586349.1 | DR_2008/NP_295731.1 | 82.34 | 334 | 59 | 0 | 1 | 334 | 1 | 334 | 0 | 558 | 99 |
| WP_019587223.1 | DR_0297/NP_294020.1 | 78.37 | 490 | 103 | 1 | 1 | 487 | 1 | 490 | 0 | 704 | 99 |
| WP_019588297.1 | DR_1674/NP_295397.1 | 86.45 | 332 | 45 | 0 | 1 | 332 | 1 | 332 | 0 | 598 | 99 |
| WP_019588678.1 | DR_1413/NP_295136.1 | 73.58 | 352 | 87 | 2 | 11 | 356 | 10 | 361 | 3.00E-178 | 500 | 96 |
| WP_019588759.1 | DR_1588/NP_295311.1 | 71.43 | 413 | 109 | 4 | 1 | 408 | 12 | 420 | 0 | 586 | 100 |
| WP_026332162.1 | DR_0963/NP_294687.2 | 85.47 | 344 | 50 | 0 | 5 | 348 | 4 | 347 | 0 | 592 | 99 |
| WP_026332290.1 | DR_1610/NP_295333.1 | 89.78 | 411 | 42 | 0 | 7 | 417 | 17 | 427 | 0 | 719 | 97 |
| WP_026332415.1 | DR_1252/NP_294976.1 | 83.21 | 405 | 68 | 0 | 1 | 405 | 1 | 405 | 0 | 725 | 100 |
| WP_026332422.1 | DR_1278/NP_295002.1 | 74.77 | 325 | 77 | 1 | 1 | 320 | 1 | 325 | 8.00E-170 | 476 | 100 |
| WP_051086938.1 | DR_1238/NP_294962.1 | 91.73 | 387 | 32 | 0 | 3 | 389 | 7 | 393 | 0 | 737 | 99 |

### *Deinococcus aquatilis*

| Query ID | Subject ID | % Identity | Alignment length | Mismatches | Gap opens | Query start | Query end | Sequence start | Sequence end | E-value | Bit score | Query coverage |
|---|---|---|---|---|---|---|---|---|---|---|---|---|
| DR_1238/NP_294962.1\|homocitrate | WP_019008047.1 | 89.11 | 395 | 34 | 1 | 8 | 393 | 4 | 398 | 0 | 728 | 98 |
| DR_1610/NP_295333.1\|3-isopropylmalate | WP_019007913.1 | 87.35 | 419 | 49 | 1 | 17 | 431 | 7 | 425 | 0 | 704 | 96 |
| DR_1614/NP_295337.1\|3-isopropylmalate | WP_019007911.1 | 83.44 | 163 | 27 | 0 | 1 | 163 | 1 | 163 | 2.00E-98 | 283 | 92 |
| DR_1674/NP_295397.1\|isocitrate | WP_019009730.1 | 88.25 | 332 | 39 | 0 | 1 | 332 | 1 | 332 | 0 | 610 | 99 |
| DR_1588/NP_295311.1\|class | WP_019009188.1 | 71.75 | 400 | 111 | 2 | 22 | 419 | 18 | 417 | 0 | 582 | 95 |
| DR_2194/NP_295916.1\|ribosomal | WP_019007907.1 | 84.38 | 288 | 40 | 1 | 8 | 295 | 1 | 283 | 9.00E-175 | 486 | 98 |
| DR_1420/NP_295143.2\|acetylglutamate/acetylaminoadipate | WP_019009650.1 | 90.64 | 267 | 25 | 0 | 1 | 267 | 1 | 267 | 1.00E-165 | 461 | 100 |
| DR_0963/NP_294687.2\|argC;N-acetyl-gamma-glutamyl-phosphate | WP_083922393.1 | 84.75 | 341 | 52 | 0 | 7 | 347 | 17 | 357 | 0 | 584 | 98 |
| DR_0794/NP_294518.1\|acetylornithine | WP_019008062.1 | 75.64 | 427 | 83 | 1 | 44 | 449 | 15 | 441 | 0 | 671 | 87 |
| DR_1413/NP_295136.1\|acetyl-lysine | WP_019009897.1 | 77.25 | 356 | 78 | 1 | 4 | 359 | 17 | 369 | 0 | 552 | 98 |
| DR_1252/NP_294976.1\|hypothetical | WP_019009907.1 | 81.98 | 405 | 73 | 0 | 1 | 405 | 1 | 405 | 0 | 714 | 100 |
| DR_1365/NP_295088.1\|aspartate | WP_040379812.1 | 81.47 | 464 | 86 | 0 | 10 | 473 | 7 | 470 | 0 | 664 | 98 |
| DR_2008/NP_295731.1\|aspartate-semialdehyde | WP_040380221.1 | 83.83 | 334 | 54 | 0 | 1 | 334 | 4 | 337 | 0 | 575 | 99 |
| DR_1278/NP_295002.1\|homoserine | WP_026298512.1 | 73.54 | 325 | 81 | 1 | 1 | 325 | 1 | 320 | 8.00E-160 | 451 | 100 |
| DR_0671/NP_294394.1\|class | WP_019011051.1 | 65.6 | 375 | 118 | 2 | 3 | 377 | 6 | 369 | 1.00E-161 | 460 | 98 |
| DR_1758/NP_295481.1\|diaminopimelate | WP_019008271.1 | 71.43 | 371 | 106 | 0 | 8 | 378 | 16 | 386 | 0 | 531 | 98 |
| DR_0297/NP_294020.1\|UDP-N-acetylmuramoylalanyl-D-glutamate--2_6-diaminopimelate | WP_019010473.1 | 78.5 | 493 | 102 | 2 | 1 | 490 | 1 | 492 | 0 | 681 | 100 |
| DR_0768/NP_294492.1\|UDP-N-acetylmuramoylalanyl-D-glutamyl-2_6-diaminopimelate--D-alanyl-D-alanyl | WP_019011137.1 | 68.71 | 441 | 118 | 5 | 1 | 431 | 1 | 431 | 0 | 577 | 99 |

**REVERSE BLAST**

| | | | | | | | | | | | | |
|---|---|---|---|---|---|---|---|---|---|---|---|---|
| WP_019007907.1 | DR_2194/NP_295916.1 | 84.38 | 288 | 40 | 1 | 1 | 283 | 8 | 295 | 3.00E-175 | 487 | 100 |
| WP_019007911.1 | DR_1614/NP_295337.1 | 83.44 | 163 | 27 | 0 | 1 | 163 | 1 | 163 | 2.00E-98 | 283 | 98 |
| WP_019007913.1 | DR_1610/NP_295333.1 | 87.35 | 419 | 49 | 1 | 7 | 425 | 17 | 431 | 0 | 704 | 98 |
| WP_019008047.1 | DR_1238/NP_294962.1 | 89.11 | 395 | 34 | 1 | 4 | 398 | 8 | 393 | 0 | 728 | 99 |
| WP_019008062.1 | DR_0794/NP_294518.1 | 75.64 | 427 | 83 | 1 | 15 | 441 | 44 | 449 | 0 | 671 | 94 |
| WP_019008271.1 | DR_1758/NP_295481.1 | 71.43 | 371 | 106 | 0 | 16 | 386 | 8 | 378 | 0 | 531 | 95 |
| WP_019009188.1 | DR_1588/NP_295311.1 | 71.75 | 400 | 111 | 2 | 18 | 417 | 22 | 419 | 0 | 567 | 96 |
| WP_019009650.1 | DR_1420/NP_295143.2 | 90.64 | 267 | 25 | 0 | 1 | 267 | 1 | 267 | 6.00E-166 | 461 | 100 |
| WP_019009730.1 | DR_1674/NP_295397.1 | 88.25 | 332 | 39 | 0 | 1 | 332 | 1 | 332 | 0 | 610 | 99 |
| WP_019009897.1 | DR_1413/NP_295136.1 | 77.25 | 356 | 78 | 1 | 17 | 369 | 4 | 359 | 0 | 552 | 93 |
| WP_019009907.1 | DR_1252/NP_294976.1 | 81.98 | 405 | 73 | 0 | 1 | 405 | 1 | 405 | 0 | 714 | 100 |
| WP_019010473.1 | DR_0297/NP_294020.1 | 78.5 | 493 | 102 | 2 | 1 | 492 | 1 | 490 | 0 | 691 | 99 |
| WP_019011051.1 | DR_0671/NP_294394.1 | 64.72 | 377 | 122 | 2 | 4 | 369 | 1 | 377 | 3.00E-160 | 456 | 95 |
| WP_019011137.1 | DR_0768/NP_294492.1 | 68.71 | 441 | 118 | 5 | 1 | 431 | 1 | 431 | 0 | 577 | 100 |
| WP_026298512.1 | DR_1278/NP_295002.1 | 73.54 | 325 | 81 | 1 | 1 | 320 | 1 | 325 | 1.00E-167 | 471 | 100 |
| WP_040379812.1 | DR_1365/NP_295088.1 | 81.47 | 464 | 86 | 0 | 7 | 470 | 10 | 473 | 0 | 695 | 99 |
| WP_040380221.1 | DR_2008/NP_295731.1 | 83.83 | 334 | 54 | 0 | 4 | 337 | 1 | 334 | 0 | 575 | 97 |
| WP_083922393.1 | DR_0963/NP_294687.2 | 84.75 | 341 | 52 | 0 | 17 | 357 | 7 | 347 | 0 | 584 | 95 |

### *Deinococcus deserti*

| Query ID | Subject ID | % Identity | Alignment length | Mismatches | Gap opens | Query start | Query end | Sequence start | Sequence end | E-value | Bit score | Query coverage |
|---|---|---|---|---|---|---|---|---|---|---|---|---|
| DR_1238/NP_294962.1\|homocitrate | WP_012693464.1 | 90.98 | 388 | 35 | 0 | 6 | 393 | 1 | 388 | 0 | 731 | 99 |
| DR_1610/NP_295333.1\|3-isopropylmalate | WP_041227498.1 | 89.47 | 418 | 44 | 0 | 15 | 432 | 5 | 422 | 0 | 730 | 96 |
| DR_1614/NP_295337.1\|3-isopropylmalate | WP_012693408.1 | 83.33 | 174 | 24 | 1 | 1 | 169 | 1 | 174 | 2.00E-104 | 298 | 95 |
| DR_1674/NP_295397.1\|isocitrate | WP_012692924.1 | 90.99 | 333 | 30 | 0 | 1 | 333 | 1 | 333 | 0 | 631 | 100 |
| DR_1588/NP_295311.1\|class | WP_012694356.1 | 79.45 | 399 | 81 | 1 | 22 | 419 | 7 | 405 | 0 | 647 | 95 |
| DR_2194/NP_295916.1\|ribosomal | WP_012693405.1 | 86.76 | 287 | 33 | 1 | 8 | 294 | 1 | 282 | 0 | 509 | 97 |
| DR_1420/NP_295143.2\|acetylglutamate/acetylaminoadipate | WP_041227135.1 | 88.76 | 267 | 30 | 0 | 1 | 267 | 1 | 267 | 8.00E-165 | 459 | 100 |
| DR_0963/NP_294687.2\|argC;N-acetyl-gamma-glutamyl-phosphate | WP_012693046.1 | 86.09 | 345 | 48 | 0 | 3 | 347 | 2 | 346 | 0 | 600 | 99 |
| DR_0794/NP_294518.1\|acetylornithine | WP_012693785.1 | 80.69 | 404 | 78 | 0 | 45 | 448 | 4 | 407 | 0 | 691 | 87 |
| DR_1413/NP_295136.1\|acetyl-lysine | WP_012693904.1 | 71.35 | 356 | 97 | 2 | 5 | 359 | 8 | 359 | 1.00E-174 | 491 | 98 |
| DR_1252/NP_294976.1\|hypothetical | WP_012692890.1 | 90.12 | 405 | 40 | 0 | 1 | 405 | 1 | 405 | 0 | 773 | 100 |
| DR_1365/NP_295088.1\|aspartate | WP_012693171.1 | 82.61 | 460 | 80 | 0 | 10 | 469 | 5 | 464 | 0 | 705 | 97 |
| DR_2008/NP_295731.1\|aspartate-semialdehyde | WP_012693661.1 | 82.63 | 334 | 58 | 0 | 1 | 334 | 1 | 334 | 0 | 566 | 99 |
| DR_1278/NP_295002.1\|homoserine | WP_012693680.1 | 79.08 | 325 | 63 | 1 | 1 | 325 | 1 | 320 | 0 | 509 | 100 |
| DR_0671/NP_294394.1\|class | WP_012693528.1 | 31.34 | 367 | 222 | 8 | 26 | 379 | 37 | 386 | 1.00E-40 | 147 | 93 |
| DR_1758/NP_295481.1\|diaminopimelate | WP_012693327.1 | 72.3 | 379 | 105 | 0 | 1 | 379 | 1 | 379 | 0 | 527 | 100 |
| DR_0297/NP_294020.1\|UDP-N-acetylmuramoylalanyl-D-glutamate--2_6-diaminopimelate | WP_012694305.1 | 80.41 | 490 | 93 | 1 | 1 | 490 | 1 | 487 | 0 | 726 | 100 |
| DR_0768/NP_294492.1\|UDP-N-acetylmuramoylalanyl-D-glutamyl-2_6-diaminopimelate--D-alanyl-D-alanyl | WP_012693854.1 | 74.94 | 439 | 97 | 4 | 1 | 433 | 1 | 432 | 0 | 636 | 100 |

**REVERSE BLAST**

| | | | | | | | | | | | | |
|---|---|---|---|---|---|---|---|---|---|---|---|---|
| WP_012692890.1 | DR_1252/NP_294976.1 | 90.12 | 405 | 40 | 0 | 1 | 405 | 1 | 405 | 0 | 773 | 100 |
| WP_012692924.1 | DR_1674/NP_295397.1 | 90.99 | 333 | 30 | 0 | 1 | 333 | 1 | 333 | 0 | 631 | 100 |
| WP_012693046.1 | DR_0963/NP_294687.2 | 86.09 | 345 | 48 | 0 | 2 | 346 | 3 | 347 | 0 | 599 | 99 |
| WP_012693171.1 | DR_1365/NP_295088.1 | 82.61 | 460 | 80 | 0 | 5 | 464 | 10 | 469 | 0 | 701 | 98 |
| WP_012693327.1 | DR_1758/NP_295481.1 | 72.3 | 379 | 105 | 0 | 1 | 379 | 1 | 379 | 0 | 550 | 100 |
| WP_012693405.1 | DR_2194/NP_295916.1 | 86.76 | 287 | 33 | 1 | 1 | 282 | 8 | 294 | 4.00E-178 | 494 | 98 |
| WP_012693408.1 | DR_1614/NP_295337.1 | 83.33 | 174 | 24 | 1 | 1 | 174 | 1 | 169 | 2.00E-104 | 298 | 99 |
| WP_012693464.1 | DR_1238/NP_294962.1 | 90.98 | 388 | 35 | 0 | 1 | 388 | 6 | 393 | 0 | 731 | 100 |
| WP_012693528.1 | NP_294346.1 | 80.62 | 387 | 75 | 0 | 1 | 387 | 1 | 387 | 0 | 634 | 99 |
| WP_012693661.1 | DR_2008/NP_295731.1 | 82.63 | 334 | 58 | 0 | 1 | 334 | 1 | 334 | 0 | 566 | 99 |
| WP_012693680.1 | DR_1278/NP_295002.1 | 79.08 | 325 | 63 | 1 | 1 | 320 | 1 | 325 | 0 | 510 | 100 |
| WP_012693785.1 | DR_0794/NP_294518.1 | 80.69 | 404 | 78 | 0 | 4 | 407 | 45 | 448 | 0 | 691 | 98 |
| WP_012693854.1 | DR_0768/NP_294492.1 | 74.94 | 439 | 97 | 4 | 1 | 432 | 1 | 433 | 0 | 636 | 99 |
| WP_012693904.1 | DR_1413/NP_295136.1 | 71.35 | 356 | 97 | 2 | 8 | 359 | 5 | 359 | 1.00E-174 | 491 | 96 |
| WP_012694305.1 | DR_0297/NP_294020.1 | 80.41 | 490 | 93 | 1 | 1 | 487 | 1 | 490 | 0 | 723 | 99 |
| WP_012694356.1 | DR_1588/NP_295311.1 | 79.45 | 399 | 81 | 1 | 7 | 405 | 22 | 419 | 0 | 639 | 98 |
| WP_041227135.1 | DR_1420/NP_295143.2 | 88.76 | 267 | 30 | 0 | 1 | 267 | 1 | 267 | 5.00E-165 | 459 | 100 |
| WP_041227498.1 | DR_1610/NP_295333.1 | 89.47 | 418 | 44 | 0 | 5 | 422 | 15 | 432 | 0 | 733 | 99 |

### *Deinococcus ficus*

| Query ID | Subject ID | % Identity | Alignment length | Mismatches | Gap opens | Query start | Query end | Sequence start | Sequence end | E-value | Bit score | Query coverage |
|---|---|---|---|---|---|---|---|---|---|---|---|---|
| DR_1238/NP_294962.1\|homocitrate | WP_043778004.1 | 85.68 | 384 | 55 | 0 | 10 | 393 | 7 | 390 | 0 | 686 | 98 |
| DR_1610/NP_295333.1\|3-isopropylmalate | WP_027461565.1 | 87.53 | 425 | 53 | 0 | 8 | 432 | 1 | 425 | 0 | 722 | 98 |
| DR_1614/NP_295337.1\|3-isopropylmalate | WP_027461563.1 | 86.83 | 167 | 22 | 0 | 1 | 167 | 1 | 167 | 1.00E-105 | 301 | 94 |
| DR_1674/NP_295397.1\|isocitrate | WP_022799649.1 | 92.19 | 333 | 26 | 0 | 1 | 333 | 1 | 333 | 0 | 637 | 100 |
| DR_1588/NP_295311.1\|class | WP_027463309.1 | 73.93 | 399 | 99 | 1 | 22 | 420 | 12 | 405 | 0 | 603 | 95 |
| DR_2194/NP_295916.1\|ribosomal | WP_027461561.1 | 85.37 | 287 | 42 | 0 | 8 | 294 | 1 | 287 | 4.00E-177 | 492 | 97 |
| DR_1420/NP_295143.2\|acetylglutamate/acetylaminoadipate | WP_027461637.1 | 88.01 | 267 | 32 | 0 | 1 | 267 | 1 | 267 | 2.00E-162 | 452 | 100 |
| DR_0963/NP_294687.2\|argC;N-acetyl-gamma-glutamyl-phosphate | WP_027461644.1 | 84.97 | 346 | 52 | 0 | 3 | 348 | 2 | 347 | 0 | 594 | 99 |
| DR_0794/NP_294518.1\|acetylornithine | WP_027461765.1 | 81.39 | 403 | 75 | 0 | 45 | 447 | 4 | 406 | 0 | 693 | 87 |
| DR_1413/NP_295136.1\|acetyl-lysine | WP_027461911.1 | 74.16 | 356 | 87 | 2 | 8 | 362 | 7 | 358 | 3.00E-167 | 473 | 98 |
| DR_1252/NP_294976.1\|hypothetical | WP_027462095.1 | 89.38 | 405 | 43 | 0 | 1 | 405 | 1 | 405 | 0 | 768 | 100 |
| DR_1365/NP_295088.1\|aspartate | WP_043776624.1 | 79.15 | 470 | 96 | 2 | 5 | 473 | 3 | 471 | 0 | 714 | 99 |
| DR_2008/NP_295731.1\|aspartate-semialdehyde | WP_027462915.1 | 81.14 | 334 | 63 | 0 | 1 | 334 | 1 | 334 | 0 | 549 | 99 |
| DR_1278/NP_295002.1\|homoserine | WP_027462024.1 | 77.85 | 325 | 67 | 1 | 1 | 325 | 1 | 320 | 0 | 517 | 100 |
| DR_0671/NP_294394.1\|class | WP_027464227.1 | 63.06 | 379 | 128 | 3 | 1 | 379 | 4 | 370 | 1.00E-154 | 442 | 99 |
| DR_1758/NP_295481.1\|diaminopimelate | WP_027462069.1 | 70.24 | 373 | 109 | 1 | 8 | 378 | 7 | 379 | 0 | 525 | 98 |
| DR_0297/NP_294020.1\|UDP-N-acetylmuramoylalanyl-D-glutamate--2_6-diaminopimelate | WP_027464116.1 | 80.6 | 469 | 89 | 1 | 24 | 490 | 20 | 488 | 0 | 671 | 95 |
| DR_0768/NP_294492.1\|UDP-N-acetylmuramoylalanyl-D-glutamyl-2_6-diaminopimelate--D-alanyl-D-alanyl | WP_027462994.1 | 73.56 | 435 | 102 | 4 | 1 | 432 | 1 | 425 | 0 | 607 | 99 |

**REVERSE BLAST**

| | | | | | | | | | | | | |
|---|---|---|---|---|---|---|---|---|---|---|---|---|
| WP_022799649.1 | DR_1674/NP_295397.1 | 92.19 | 333 | 26 | 0 | 1 | 333 | 1 | 333 | 0 | 637 | 100 |
| WP_027461561.1 | DR_2194/NP_295916.1 | 85.07 | 288 | 43 | 0 | 1 | 288 | 8 | 295 | 1.00E-178 | 495 | 100 |
| WP_027461563.1 | DR_1614/NP_295337.1 | 86.83 | 167 | 22 | 0 | 1 | 167 | 1 | 167 | 1.00E-105 | 301 | 100 |
| WP_027461565.1 | DR_1610/NP_295333.1 | 87.53 | 425 | 53 | 0 | 1 | 425 | 8 | 432 | 0 | 724 | 100 |
| WP_027461637.1 | DR_1420/NP_295143.2 | 88.01 | 267 | 32 | 0 | 1 | 267 | 1 | 267 | 2.00E-162 | 452 | 100 |
| WP_027461644.1 | DR_0963/NP_294687.2 | 85.22 | 345 | 51 | 0 | 2 | 346 | 3 | 347 | 0 | 594 | 99 |
| WP_027461765.1 | DR_0794/NP_294518.1 | 81.39 | 403 | 75 | 0 | 4 | 406 | 45 | 447 | 0 | 693 | 97 |
| WP_027461911.1 | DR_1413/NP_295136.1 | 75 | 356 | 84 | 2 | 7 | 358 | 8 | 362 | 0 | 516 | 96 |
| WP_027462024.1 | DR_1278/NP_295002.1 | 77.85 | 325 | 67 | 1 | 1 | 320 | 1 | 325 | 2.00E-179 | 500 | 100 |
| WP_027462069.1 | DR_1758/NP_295481.1 | 70.24 | 373 | 109 | 1 | 7 | 379 | 8 | 378 | 0 | 525 | 98 |
| WP_027462095.1 | DR_1252/NP_294976.1 | 89.38 | 405 | 43 | 0 | 1 | 405 | 1 | 405 | 0 | 768 | 100 |
| WP_027462915.1 | DR_2008/NP_295731.1 | 81.14 | 334 | 63 | 0 | 1 | 334 | 1 | 334 | 0 | 549 | 99 |
| WP_027462994.1 | DR_0768/NP_294492.1 | 73.56 | 435 | 102 | 4 | 1 | 425 | 1 | 432 | 0 | 607 | 98 |
| WP_027463309.1 | DR_1588/NP_295311.1 | 73.93 | 399 | 99 | 1 | 12 | 405 | 22 | 420 | 0 | 593 | 97 |
| WP_027464116.1 | DR_0297/NP_294020.1 | 79.07 | 492 | 97 | 3 | 1 | 488 | 1 | 490 | 0 | 689 | 99 |
| WP_027464227.1 | DR_0671/NP_294394.1 | 62.8 | 379 | 129 | 3 | 4 | 370 | 1 | 379 | 1.00E-146 | 421 | 97 |
| WP_043776624.1 | DR_1365/NP_295088.1 | 79.15 | 470 | 96 | 2 | 3 | 471 | 5 | 473 | 0 | 687 | 99 |
| WP_043778004.1 | DR_1238/NP_294962.1 | 85.68 | 384 | 55 | 0 | 7 | 390 | 10 | 393 | 0 | 686 | 98 |

### *Deinococcus frigens*

| Query ID | Subject ID | % Identity | Alignment length | Mismatches | Gap opens | Query start | Query end | Sequence start | Sequence end | E-value | Bit score | Query coverage |
|---|---|---|---|---|---|---|---|---|---|---|---|---|
| DR_1238/NP_294962.1\|homocitrate | WP_051668072.1 | 88.83 | 385 | 43 | 0 | 9 | 393 | 11 | 395 | 0 | 716 | 98 |
| DR_1610/NP_295333.1\|3-isopropylmalate | WP_029478225.1 | 87.59 | 419 | 50 | 1 | 9 | 427 | 4 | 420 | 0 | 708 | 97 |
| DR_1614/NP_295337.1\|3-isopropylmalate | WP_029476821.1 | 83.63 | 171 | 28 | 0 | 1 | 171 | 1 | 171 | 2.00E-103 | 296 | 97 |
| DR_1674/NP_295397.1\|isocitrate | WP_029478273.1 | 88.55 | 332 | 38 | 0 | 1 | 332 | 1 | 332 | 0 | 605 | 99 |
| DR_1588/NP_295311.1\|class | WP_029477561.1 | 75.56 | 405 | 93 | 2 | 16 | 419 | 3 | 402 | 0 | 611 | 96 |
| DR_2194/NP_295916.1\|ribosomal | WP_029476826.1 | 81.94 | 288 | 47 | 1 | 8 | 295 | 1 | 283 | 2.00E-164 | 459 | 98 |
| DR_1420/NP_295143.2\|acetylglutamate/acetylaminoadipate | WP_029477634.1 | 90.26 | 267 | 26 | 0 | 1 | 267 | 1 | 267 | 3.00E-166 | 462 | 100 |
| DR_0963/NP_294687.2\|argC;N-acetyl-gamma-glutamyl-phosphate | WP_029475842.1 | 85.84 | 346 | 49 | 0 | 2 | 347 | 5 | 350 | 0 | 626 | 99 |
| DR_0794/NP_294518.1\|acetylornithine | WP_029477833.1 | 83.99 | 406 | 65 | 0 | 45 | 450 | 11 | 416 | 0 | 726 | 87 |
| DR_1413/NP_295136.1\|acetyl-lysine | WP_034417539.1 | 72.7 | 359 | 92 | 3 | 3 | 359 | 10 | 364 | 6.00E-179 | 503 | 99 |
| DR_1252/NP_294976.1\|hypothetical | WP_029478751.1 | 85.89 | 404 | 57 | 0 | 1 | 404 | 1 | 404 | 0 | 723 | 99 |
| DR_1365/NP_295088.1\|aspartate | WP_029478216.1 | 83.62 | 458 | 75 | 0 | 10 | 467 | 5 | 462 | 0 | 719 | 97 |
| DR_2008/NP_295731.1\|aspartate-semialdehyde | WP_029478118.1 | 83.53 | 334 | 55 | 0 | 1 | 334 | 1 | 334 | 0 | 576 | 99 |
| DR_1278/NP_295002.1\|homoserine | WP_029477925.1 | 80.31 | 325 | 59 | 1 | 1 | 325 | 1 | 320 | 0 | 527 | 100 |
| DR_0671/NP_294394.1\|class | WP_029477190.1 | 66.23 | 382 | 119 | 3 | 3 | 380 | 6 | 381 | 3.00E-163 | 464 | 99 |
| DR_1758/NP_295481.1\|diaminopimelate | WP_029476970.1 | 71.61 | 384 | 104 | 1 | 1 | 379 | 1 | 384 | 0 | 510 | 100 |
| DR_0297/NP_294020.1\|UDP-N-acetylmuramoylalanyl-D-glutamate--2_6-diaminopimelate | WP_029479609.1 | 76.34 | 486 | 108 | 2 | 1 | 479 | 1 | 486 | 0 | 664 | 98 |
| DR_0768/NP_294492.1\|UDP-N-acetylmuramoylalanyl-D-glutamyl-2_6-diaminopimelate--D-alanyl-D-alanyl | WP_029475883.1 | 74.13 | 433 | 106 | 2 | 1 | 432 | 1 | 428 | 0 | 602 | 99 |

**REVERSE BLAST**

| | | | | | | | | | | | | |
|---|---|---|---|---|---|---|---|---|---|---|---|---|
| WP_029475842.1 | DR_0963/NP_294687.2 | 85.84 | 346 | 49 | 0 | 5 | 350 | 2 | 347 | 0 | 601 | 99 |
| WP_029475883.1 | DR_0768/NP_294492.1 | 74.13 | 433 | 106 | 2 | 1 | 428 | 1 | 432 | 0 | 624 | 98 |
| WP_029476821.1 | DR_1614/NP_295337.1 | 83.63 | 171 | 28 | 0 | 1 | 171 | 1 | 171 | 2.00E-103 | 296 | 98 |
| WP_029476826.1 | DR_2194/NP_295916.1 | 81.94 | 288 | 47 | 1 | 1 | 283 | 8 | 295 | 1.00E-166 | 465 | 100 |
| WP_029476970.1 | DR_1758/NP_295481.1 | 71.61 | 384 | 104 | 1 | 1 | 384 | 1 | 379 | 0 | 545 | 100 |
| WP_029477190.1 | DR_0671/NP_294394.1 | 65.62 | 384 | 122 | 3 | 4 | 381 | 1 | 380 | 3.00E-165 | 469 | 98 |
| WP_029477561.1 | DR_1588/NP_295311.1 | 75.56 | 405 | 93 | 2 | 3 | 402 | 16 | 419 | 0 | 603 | 99 |
| WP_029477634.1 | DR_1420/NP_295143.2 | 90.26 | 267 | 26 | 0 | 1 | 267 | 1 | 267 | 2.00E-166 | 462 | 100 |
| WP_029477833.1 | DR_0794/NP_294518.1 | 83.99 | 406 | 65 | 0 | 11 | 416 | 45 | 450 | 0 | 726 | 97 |
| WP_029477925.1 | DR_1278/NP_295002.1 | 80.31 | 325 | 59 | 1 | 1 | 320 | 1 | 325 | 0 | 509 | 100 |
| WP_029478118.1 | DR_2008/NP_295731.1 | 83.53 | 334 | 55 | 0 | 1 | 334 | 1 | 334 | 0 | 576 | 99 |
| WP_029478216.1 | DR_1365/NP_295088.1 | 83.62 | 458 | 75 | 0 | 5 | 462 | 10 | 467 | 0 | 704 | 97 |
| WP_029478225.1 | DR_1610/NP_295333.1 | 87.59 | 419 | 50 | 1 | 4 | 420 | 9 | 427 | 0 | 709 | 98 |
| WP_029478273.1 | DR_1674/NP_295397.1 | 88.55 | 332 | 38 | 0 | 1 | 332 | 1 | 332 | 0 | 605 | 100 |
| WP_029478751.1 | DR_1252/NP_294976.1 | 85.89 | 404 | 57 | 0 | 1 | 404 | 1 | 404 | 0 | 738 | 100 |
| WP_029479609.1 | DR_0297/NP_294020.1 | 76.16 | 495 | 111 | 2 | 1 | 495 | 1 | 488 | 0 | 677 | 98 |
| WP_034417539.1 | DR_1413/NP_295136.1 | 72.7 | 359 | 92 | 3 | 10 | 364 | 3 | 359 | 5.00E-179 | 503 | 94 |
| WP_051668072.1 | DR_1238/NP_294962.1 | 88.83 | 385 | 43 | 0 | 11 | 395 | 9 | 393 | 0 | 716 | 97 |

### *Deinococcus geothermalis*

| Query ID | Subject ID | % Identity | Alignment length | Mismatches | Gap opens | Query start | Query end | Sequence start | Sequence end | E-value | Bit score | Query coverage |
|---|---|---|---|---|---|---|---|---|---|---|---|---|
| DR_1238/NP_294962.1\|homocitrate | WP_011530390.1 | 90.41 | 386 | 37 | 0 | 8 | 393 | 4 | 389 | 0 | 722 | 98 |
| DR_1610/NP_295333.1\|3-isopropylmalate | WP_011530290.1 | 88.1 | 420 | 47 | 1 | 8 | 427 | 1 | 417 | 0 | 715 | 97 |
| DR_1614/NP_295337.1\|3-isopropylmalate | WP_011530288.1 | 88.41 | 164 | 19 | 0 | 1 | 164 | 1 | 164 | 8.00E-108 | 306 | 93 |
| DR_1674/NP_295397.1\|isocitrate | WP_011530587.1 | 86.75 | 332 | 44 | 0 | 1 | 332 | 1 | 332 | 0 | 600 | 99 |
| DR_1588/NP_295311.1\|class | WP_011531204.1 | 75.36 | 418 | 88 | 5 | 4 | 419 | 1 | 405 | 0 | 579 | 99 |
| DR_2194/NP_295916.1\|ribosomal | WP_011530285.1 | 86.41 | 287 | 39 | 0 | 8 | 294 | 1 | 287 | 0 | 503 | 97 |
| DR_1420/NP_295143.2\|acetylglutamate/acetylaminoadipate | WP_011529821.1 | 87.64 | 267 | 33 | 0 | 1 | 267 | 1 | 267 | 5.00E-163 | 454 | 100 |
| DR_0963/NP_294687.2\|argC;N-acetyl-gamma-glutamyl-phosphate | WP_011529828.1 | 83.48 | 345 | 57 | 0 | 3 | 347 | 4 | 348 | 0 | 604 | 99 |
| DR_0794/NP_294518.1\|acetylornithine | WP_011530545.1 | 83.62 | 409 | 67 | 0 | 37 | 445 | 1 | 409 | 0 | 721 | 88 |
| DR_1413/NP_295136.1\|acetyl-lysine | WP_011530522.1 | 70.64 | 361 | 100 | 2 | 1 | 361 | 1 | 355 | 0 | 507 | 99 |
| DR_1252/NP_294976.1\|hypothetical | WP_011530136.1 | 80.99 | 405 | 77 | 0 | 1 | 405 | 1 | 405 | 0 | 714 | 100 |
| DR_1365/NP_295088.1\|aspartate | WP_011530261.1 | 82.97 | 464 | 79 | 0 | 10 | 473 | 5 | 468 | 0 | 714 | 98 |
| DR_2008/NP_295731.1\|aspartate-semialdehyde | WP_011530908.1 | 80.24 | 334 | 66 | 0 | 1 | 334 | 1 | 334 | 0 | 539 | 99 |
| DR_1278/NP_295002.1\|homoserine | WP_011529754.1 | 76 | 325 | 73 | 1 | 1 | 325 | 1 | 320 | 7.00E-170 | 476 | 100 |
| DR_0671/NP_294394.1\|class | WP_011531189.1 | 65.52 | 377 | 119 | 2 | 1 | 377 | 4 | 369 | 6.00E-164 | 465 | 99 |
| DR_1758/NP_295481.1\|diaminopimelate | WP_011529933.1 | 70.94 | 382 | 107 | 1 | 1 | 378 | 1 | 382 | 2.00E-178 | 502 | 99 |
| DR_0297/NP_294020.1\|UDP-N-acetylmuramoylalanyl-D-glutamate--2_6-diaminopimelate | WP_011529249.1 | 81.17 | 478 | 87 | 1 | 1 | 478 | 1 | 475 | 0 | 719 | 98 |
| DR_0768/NP_294492.1\|UDP-N-acetylmuramoylalanyl-D-glutamyl-2_6-diaminopimelate--D-alanyl-D-alanyl | WP_011530849.1 | 73.74 | 438 | 98 | 5 | 1 | 431 | 1 | 428 | 0 | 590 | 99 |

**REVERSE BLAST**

| | | | | | | | | | | | | |
|---|---|---|---|---|---|---|---|---|---|---|---|---|
| WP_011529249.1 | DR_0297/NP_294020.1 | 80.82 | 490 | 91 | 1 | 1 | 487 | 1 | 490 | 0 | 726 | 99 |
| WP_011529754.1 | DR_1278/NP_295002.1 | 76 | 325 | 73 | 1 | 1 | 320 | 1 | 325 | 6.00E-170 | 476 | 100 |
| WP_011529821.1 | DR_1420/NP_295143.2 | 87.64 | 267 | 33 | 0 | 1 | 267 | 1 | 267 | 1.00E-162 | 453 | 100 |
| WP_011529828.1 | DR_0963/NP_294687.2 | 83.48 | 345 | 57 | 0 | 4 | 348 | 3 | 347 | 0 | 582 | 99 |
| WP_011529933.1 | DR_1758/NP_295481.1 | 70.94 | 382 | 107 | 1 | 1 | 382 | 1 | 378 | 0 | 528 | 99 |
| WP_011530136.1 | DR_1252/NP_294976.1 | 80.99 | 405 | 77 | 0 | 1 | 405 | 1 | 405 | 0 | 714 | 100 |
| WP_011530261.1 | DR_1365/NP_295088.1 | 82.97 | 464 | 79 | 0 | 5 | 468 | 10 | 473 | 0 | 698 | 99 |
| WP_011530285.1 | DR_2194/NP_295916.1 | 86.41 | 287 | 39 | 0 | 1 | 287 | 8 | 294 | 0 | 503 | 98 |
| WP_011530288.1 | DR_1614/NP_295337.1 | 88.41 | 164 | 19 | 0 | 1 | 164 | 1 | 164 | 8.00E-108 | 306 | 97 |
| WP_011530290.1 | DR_1610/NP_295333.1 | 88.1 | 420 | 47 | 1 | 1 | 417 | 8 | 427 | 0 | 717 | 99 |
| WP_011530390.1 | DR_1238/NP_294962.1 | 90.41 | 386 | 37 | 0 | 4 | 389 | 8 | 393 | 0 | 722 | 99 |
| WP_011530522.1 | DR_1413/NP_295136.1 | 70.64 | 361 | 100 | 2 | 1 | 355 | 1 | 361 | 0 | 507 | 99 |
| WP_011530545.1 | DR_0794/NP_294518.1 | 83.62 | 409 | 67 | 0 | 1 | 409 | 37 | 445 | 0 | 721 | 99 |
| WP_011530587.1 | DR_1674/NP_295397.1 | 86.75 | 332 | 44 | 0 | 1 | 332 | 1 | 332 | 0 | 600 | 99 |
| WP_011530849.1 | DR_0768/NP_294492.1 | 73.74 | 438 | 98 | 5 | 1 | 428 | 1 | 431 | 0 | 618 | 99 |
| WP_011530908.1 | DR_2008/NP_295731.1 | 80.24 | 334 | 66 | 0 | 1 | 334 | 1 | 334 | 0 | 539 | 99 |
| WP_011531189.1 | DR_0671/NP_294394.1 | 65.25 | 377 | 120 | 2 | 4 | 369 | 1 | 377 | 1.00E-160 | 457 | 98 |
| WP_011531204.1 | DR_1588/NP_295311.1 | 74.4 | 418 | 92 | 5 | 1 | 405 | 4 | 419 | 0 | 606 | 99 |

### *Deinococcus gobiensis*

| Query ID | Subject ID | % Identity | Alignment length | Mismatches | Gap opens | Query start | Query end | Sequence start | Sequence end | E-value | Bit score | Query coverage |
|---|---|---|---|---|---|---|---|---|---|---|---|---|
| DR_1238/NP_294962.1\|homocitrate | WP_014684826.1 | 89.69 | 388 | 40 | 0 | 6 | 393 | 1 | 388 | 0 | 728 | 99 |
| DR_1610/NP_295333.1\|3-isopropylmalate | WP_014684716.1 | 88.81 | 420 | 47 | 0 | 8 | 427 | 1 | 420 | 0 | 728 | 97 |
| DR_1614/NP_295337.1\|3-isopropylmalate | WP_014684718.1 | 88.1 | 168 | 15 | 1 | 1 | 163 | 1 | 168 | 5.00E-105 | 300 | 92 |
| DR_1674/NP_295397.1\|isocitrate | WP_014684534.1 | 90.39 | 333 | 32 | 0 | 1 | 333 | 1 | 333 | 0 | 623 | 100 |
| DR_1588/NP_295311.1\|class | AFD24357.1 | 76.42 | 386 | 91 | 0 | 34 | 419 | 1 | 386 | 0 | 599 | 92 |
| DR_2194/NP_295916.1\|ribosomal | WP_085961051.1 | 85.08 | 295 | 39 | 1 | 1 | 295 | 1 | 290 | 0 | 509 | 100 |
| DR_1420/NP_295143.2\|acetylglutamate/acetylaminoadipate | WP_043803425.1 | 92.88 | 267 | 19 | 0 | 1 | 267 | 1 | 267 | 4.00E-171 | 475 | 100 |
| DR_0963/NP_294687.2\|argC;N-acetyl-gamma-glutamyl-phosphate | WP_083847250.1 | 86.09 | 345 | 48 | 0 | 3 | 347 | 4 | 348 | 0 | 598 | 99 |
| DR_0794/NP_294518.1\|acetylornithine | WP_043801752.1 | 81.15 | 419 | 78 | 1 | 37 | 455 | 1 | 418 | 0 | 715 | 90 |
| DR_1413/NP_295136.1\|acetyl-lysine | WP_050920684.1 | 77.34 | 353 | 77 | 1 | 9 | 361 | 6 | 355 | 0 | 509 | 98 |
| DR_1252/NP_294976.1\|hypothetical | WP_014685142.1 | 89.38 | 405 | 43 | 0 | 1 | 405 | 1 | 405 | 0 | 768 | 100 |
| DR_1365/NP_295088.1\|aspartate | WP_014684982.1 | 85.13 | 464 | 69 | 0 | 10 | 473 | 5 | 468 | 0 | 744 | 98 |
| DR_2008/NP_295731.1\|aspartate-semialdehyde | WP_014684233.1 | 83.83 | 334 | 54 | 0 | 1 | 334 | 1 | 334 | 0 | 567 | 99 |
| DR_1278/NP_295002.1\|homoserine | WP_043801217.1 | 81.23 | 325 | 56 | 1 | 1 | 325 | 1 | 320 | 0 | 521 | 100 |
| DR_0671/NP_294394.1\|class | WP_014683795.1 | 67.82 | 376 | 109 | 5 | 3 | 376 | 6 | 371 | 2.00E-159 | 454 | 98 |
| DR_1758/NP_295481.1\|diaminopimelate | WP_043801459.1 | 71.51 | 372 | 106 | 0 | 8 | 379 | 7 | 378 | 6.00E-173 | 489 | 98 |
| DR_0297/NP_294020.1\|UDP-N-acetylmuramoylalanyl-D-glutamate--2_6-diaminopimelate | WP_014683589.1 | 80 | 490 | 95 | 1 | 1 | 490 | 1 | 487 | 0 | 731 | 100 |
| DR_0768/NP_294492.1\|UDP-N-acetylmuramoylalanyl-D-glutamyl-2_6-diaminopimelate--D-alanyl-D-alanyl | WP_014685666.1 | 74.31 | 436 | 103 | 2 | 1 | 432 | 1 | 431 | 0 | 620 | 99 |

**REVERSE BLAST**

| | | | | | | | | | | | | |
|---|---|---|---|---|---|---|---|---|---|---|---|---|
| WP_014683589.1 | DR_0297/NP_294020.1 | 80 | 490 | 95 | 1 | 1 | 487 | 1 | 490 | 0 | 717 | 100 |
| WP_014683795.1 | DR_0671/NP_294394.1 | 67.2 | 378 | 112 | 5 | 4 | 371 | 1 | 376 | 1.00E-159 | 454 | 98 |
| WP_014684233.1 | DR_2008/NP_295731.1 | 83.83 | 334 | 54 | 0 | 1 | 334 | 1 | 334 | 0 | 567 | 99 |
| WP_014684534.1 | DR_1674/NP_295397.1 | 90.39 | 333 | 32 | 0 | 1 | 333 | 1 | 333 | 0 | 623 | 100 |
| WP_014684716.1 | DR_1610/NP_295333.1 | 88.81 | 420 | 47 | 0 | 1 | 420 | 8 | 427 | 0 | 727 | 99 |
| WP_014684718.1 | DR_1614/NP_295337.1 | 88.1 | 168 | 15 | 1 | 1 | 168 | 1 | 163 | 4.00E-105 | 300 | 98 |
| WP_014684826.1 | DR_1238/NP_294962.1 | 89.69 | 388 | 40 | 0 | 1 | 388 | 6 | 393 | 0 | 728 | 100 |
| WP_014684982.1 | DR_1365/NP_295088.1 | 85.13 | 464 | 69 | 0 | 5 | 468 | 10 | 473 | 0 | 733 | 99 |
| WP_014685142.1 | DR_1252/NP_294976.1 | 89.38 | 405 | 43 | 0 | 1 | 405 | 1 | 405 | 0 | 768 | 100 |
| WP_014685666.1 | DR_0768/NP_294492.1 | 74.31 | 436 | 103 | 2 | 1 | 431 | 1 | 432 | 0 | 620 | 99 |
| WP_043801217.1 | DR_1278/NP_295002.1 | 81.23 | 325 | 56 | 1 | 1 | 320 | 1 | 325 | 0 | 505 | 100 |
| WP_043801459.1 | DR_1758/NP_295481.1 | 71.51 | 372 | 106 | 0 | 7 | 378 | 8 | 379 | 0 | 523 | 96 |
| WP_043801752.1 | DR_0794/NP_294518.1 | 81.15 | 419 | 78 | 1 | 1 | 418 | 37 | 455 | 0 | 715 | 100 |
| WP_043803425.1 | DR_1420/NP_295143.2 | 92.88 | 267 | 19 | 0 | 1 | 267 | 1 | 267 | 4.00E-171 | 474 | 100 |
| WP_050920684.1 | DR_1413/NP_295136.1 | 77.34 | 353 | 77 | 1 | 6 | 355 | 9 | 361 | 0 | 530 | 94 |
| WP_083847250.1 | DR_0963/NP_294687.2 | 86.09 | 345 | 48 | 0 | 4 | 348 | 3 | 347 | 0 | 598 | 99 |
| WP_085961051.1 | DR_2194/NP_295916.1 | 85.08 | 295 | 39 | 1 | 1 | 290 | 1 | 295 | 3.00E-177 | 492 | 100 |
| AFD24357.1 | WP_010888227.1 | 76.425 | 386 | 91 | 0 | 1 | 386 | 34 | 419 | 0 | 591 | 99 |

### *Deinococcus grandis*

| Query ID | Subject ID | % Identity | Alignment length | Mismatches | Gap opens | Query start | Query end | Sequence start | Sequence end | E-value | Bit score | Query coverage |
|---|---|---|---|---|---|---|---|---|---|---|---|---|
| DR_1238/NP_294962.1\|homocitrate | WP_058976346.1 | 91.99 | 387 | 31 | 0 | 7 | 393 | 9 | 395 | 0 | 742 | 98 |
| DR_1610/NP_295333.1\|3-isopropylmalate | WP_058976666.1 | 88.49 | 417 | 44 | 1 | 15 | 427 | 5 | 421 | 0 | 708 | 95 |
| DR_1614/NP_295337.1\|3-isopropylmalate | WP_058976665.1 | 89.51 | 162 | 17 | 0 | 1 | 162 | 1 | 162 | 4.00E-106 | 302 | 92 |
| DR_1674/NP_295397.1\|isocitrate | WP_058975975.1 | 92.19 | 333 | 26 | 0 | 1 | 333 | 1 | 333 | 0 | 635 | 100 |
| DR_1588/NP_295311.1\|class | WP_058978231.1 | 75.62 | 406 | 93 | 2 | 16 | 420 | 6 | 406 | 0 | 617 | 96 |
| DR_2194/NP_295916.1\|ribosomal | WP_058976662.1 | 89.55 | 287 | 30 | 0 | 8 | 294 | 1 | 287 | 0 | 510 | 97 |
| DR_1420/NP_295143.2\|acetylglutamate/acetylaminoadipate | WP_058976731.1 | 87.27 | 267 | 34 | 0 | 1 | 267 | 1 | 267 | 9.00E-161 | 449 | 100 |
| DR_0963/NP_294687.2\|argC;N-acetyl-gamma-glutamyl-phosphate | WP_083523981.1 | 87.21 | 344 | 44 | 0 | 4 | 347 | 5 | 348 | 0 | 599 | 99 |
| DR_0794/NP_294518.1\|acetylornithine | WP_058976578.1 | 82 | 411 | 74 | 0 | 45 | 455 | 6 | 416 | 0 | 692 | 88 |
| DR_1413/NP_295136.1\|acetyl-lysine | WP_058975089.1 | 73.31 | 356 | 90 | 2 | 8 | 362 | 7 | 358 | 1.00E-179 | 504 | 98 |
| DR_1252/NP_294976.1\|hypothetical | WP_058977807.1 | 89.88 | 405 | 41 | 0 | 1 | 405 | 1 | 405 | 0 | 773 | 100 |
| DR_1365/NP_295088.1\|aspartate | WP_058976535.1 | 83.41 | 458 | 76 | 0 | 10 | 467 | 5 | 462 | 0 | 718 | 98 |
| DR_1365/NP_295088.1\|aspartate | WP_058976535.1 | 34.29 | 70 | 46 | 0 | 402 | 471 | 322 | 391 | 4.3 | 26.6 | 98 |
| DR_2008/NP_295731.1\|aspartate-semialdehyde | WP_058977719.1 | 82.49 | 337 | 59 | 0 | 1 | 337 | 1 | 337 | 0 | 573 | 99 |
| DR_1278/NP_295002.1\|homoserine | WP_058977494.1 | 78.15 | 325 | 66 | 1 | 1 | 325 | 1 | 320 | 4.00E-178 | 497 | 100 |
| DR_0671/NP_294394.1\|class | WP_058978387.1 | 65.16 | 376 | 118 | 4 | 1 | 376 | 4 | 366 | 2.00E-144 | 416 | 99 |
| DR_1758/NP_295481.1\|diaminopimelate | WP_083524034.1 | 68.62 | 376 | 118 | 0 | 3 | 378 | 45 | 420 | 9.00E-172 | 487 | 99 |
| DR_0297/NP_294020.1\|UDP-N-acetylmuramoylalanyl-D-glutamate--2_6-diaminopimelate | WP_058974787.1 | 79.3 | 483 | 99 | 1 | 1 | 483 | 1 | 482 | 0 | 694 | 99 |
| DR_0768/NP_294492.1\|UDP-N-acetylmuramoylalanyl-D-glutamyl-2_6-diaminopimelate--D-alanyl-D-alanyl | WP_058975618.1 | 74.77 | 440 | 98 | 5 | 1 | 432 | 1 | 435 | 0 | 629 | 99 |

**REVERSE BLAST**

| | | | | | | | | | | | | |
|---|---|---|---|---|---|---|---|---|---|---|---|---|
| WP_058974787.1 | DR_0297/NP_294020.1 | 79.3 | 483 | 99 | 1 | 1 | 482 | 1 | 483 | 0 | 692 | 99 |
| WP_058975089.1 | DR_1413/NP_295136.1 | 73.31 | 356 | 90 | 2 | 7 | 358 | 8 | 362 | 9.00E-180 | 504 | 97 |
| WP_058975618.1 | DR_0768/NP_294492.1 | 74.77 | 440 | 98 | 5 | 1 | 435 | 1 | 432 | 0 | 629 | 99 |
| WP_058975975.1 | DR_1674/NP_295397.1 | 92.19 | 333 | 26 | 0 | 1 | 333 | 1 | 333 | 0 | 635 | 100 |
| WP_058976346.1 | DR_1238/NP_294962.1 | 91.99 | 387 | 31 | 0 | 9 | 395 | 7 | 393 | 0 | 742 | 98 |
| WP_058976535.1 | DR_1365/NP_295088.1 | 83.41 | 458 | 76 | 0 | 5 | 462 | 10 | 467 | 0 | 706 | 98 |
| WP_058976535.1 | DR_1365/NP_295088.1 | 32.89 | 76 | 51 | 0 | 316 | 391 | 396 | 471 | 1.00E-05 | 44.3 | 98 |
| WP_058976578.1 | DR_0794/NP_294518.1 | 82 | 411 | 74 | 0 | 6 | 416 | 45 | 455 | 0 | 713 | 99 |
| WP_058976662.1 | DR_2194/NP_295916.1 | 89.55 | 287 | 30 | 0 | 1 | 287 | 8 | 294 | 0 | 509 | 100 |
| WP_058976665.1 | DR_1614/NP_295337.1 | 89.51 | 162 | 17 | 0 | 1 | 162 | 1 | 162 | 3.00E-106 | 302 | 99 |
| WP_058976666.1 | DR_1610/NP_295333.1 | 88.49 | 417 | 44 | 1 | 5 | 421 | 15 | 427 | 0 | 708 | 97 |
| WP_058976731.1 | DR_1420/NP_295143.2 | 87.27 | 267 | 34 | 0 | 1 | 267 | 1 | 267 | 2.00E-160 | 447 | 100 |
| WP_058977494.1 | DR_1278/NP_295002.1 | 78.15 | 325 | 66 | 1 | 1 | 320 | 1 | 325 | 3.00E-178 | 497 | 100 |
| WP_058977719.1 | DR_2008/NP_295731.1 | 82.49 | 337 | 59 | 0 | 1 | 337 | 1 | 337 | 0 | 573 | 99 |
| WP_058977807.1 | DR_1252/NP_294976.1 | 89.88 | 405 | 41 | 0 | 1 | 405 | 1 | 405 | 0 | 773 | 100 |
| WP_058978231.1 | DR_1588/NP_295311.1 | 74.88 | 406 | 96 | 2 | 6 | 406 | 16 | 420 | 0 | 610 | 98 |
| WP_058978387.1 | DR_0671/NP_294394.1 | 64.89 | 376 | 119 | 4 | 4 | 366 | 1 | 376 | 2.00E-153 | 438 | 98 |
| WP_083523981.1 | DR_0963/NP_294687.2 | 87.21 | 344 | 44 | 0 | 5 | 348 | 4 | 347 | 0 | 599 | 99 |
| WP_083524034.1 | DR_1758/NP_295481.1 | 68.62 | 376 | 118 | 0 | 45 | 420 | 3 | 378 | 0 | 513 | 89 |

### *Deinococcus hopiensis*

| Query ID | Subject ID | % Identity | Alignment length | Mismatches | Gap opens | Query start | Query end | Sequence start | Sequence end | E-value | Bit score | Query coverage |
|---|---|---|---|---|---|---|---|---|---|---|---|---|
| DR_1238/NP_294962.1\|homocitrate | WP_084049092.1 | 90.46 | 388 | 37 | 0 | 6 | 393 | 1 | 388 | 0 | 724 | 99 |
| DR_1610/NP_295333.1\|3-isopropylmalate | WP_084048988.1 | 89.42 | 416 | 44 | 0 | 17 | 432 | 7 | 422 | 0 | 724 | 96 |
| DR_1614/NP_295337.1\|3-isopropylmalate | WP_084048990.1 | 87.04 | 162 | 21 | 0 | 1 | 162 | 1 | 162 | 6.00E-104 | 297 | 92 |
| DR_1674/NP_295397.1\|isocitrate | WP_084048692.1 | 85.24 | 332 | 49 | 0 | 1 | 332 | 1 | 332 | 0 | 589 | 99 |
| DR_1588/NP_295311.1\|class | WP_084050426.1 | 73.25 | 400 | 100 | 3 | 22 | 419 | 15 | 409 | 0 | 553 | 95 |
| DR_2194/NP_295916.1\|ribosomal | WP_084048993.1 | 87.85 | 288 | 30 | 1 | 8 | 295 | 1 | 283 | 4.00E-178 | 495 | 98 |
| DR_1420/NP_295143.2\|acetylglutamate/acetylaminoadipate | WP_084050826.1 | 90.23 | 266 | 26 | 0 | 1 | 266 | 1 | 266 | 7.00E-160 | 447 | 99 |
| DR_0963/NP_294687.2\|argC;N-acetyl-gamma-glutamyl-phosphate | WP_084050898.1 | 86.98 | 338 | 44 | 0 | 11 | 348 | 1 | 338 | 0 | 591 | 97 |
| DR_0794/NP_294518.1\|acetylornithine | WP_084048503.1 | 83.29 | 407 | 67 | 1 | 43 | 448 | 5 | 411 | 0 | 715 | 87 |
| DR_1413/NP_295136.1\|acetyl-lysine | WP_084048755.1 | 73.5 | 351 | 92 | 1 | 9 | 359 | 3 | 352 | 7.00E-179 | 503 | 97 |
| DR_1252/NP_294976.1\|hypothetical | WP_084049365.1 | 83.7 | 405 | 66 | 0 | 1 | 405 | 1 | 405 | 0 | 726 | 100 |
| DR_1365/NP_295088.1\|aspartate | WP_084049015.1 | 81.47 | 464 | 86 | 0 | 10 | 473 | 5 | 468 | 0 | 716 | 98 |
| DR_2008/NP_295731.1\|aspartate-semialdehyde | WP_084049619.1 | 83.68 | 337 | 55 | 0 | 1 | 337 | 1 | 337 | 0 | 566 | 99 |
| DR_1278/NP_295002.1\|homoserine | WP_084049592.1 | 76 | 325 | 73 | 1 | 1 | 325 | 1 | 320 | 1.00E-170 | 478 | 100 |
| DR_0671/NP_294394.1\|class | WP_084050371.1 | 63.49 | 378 | 127 | 2 | 3 | 380 | 6 | 372 | 1.00E-166 | 473 | 99 |
| DR_1758/NP_295481.1\|diaminopimelate | WP_084049094.1 | 72.03 | 379 | 106 | 0 | 1 | 379 | 1 | 379 | 0 | 526 | 100 |
| DR_0297/NP_294020.1\|UDP-N-acetylmuramoylalanyl-D-glutamate--2_6-diaminopimelate | WP_084050599.1 | 77.5 | 480 | 105 | 1 | 1 | 480 | 1 | 477 | 0 | 687 | 98 |
| DR_0768/NP_294492.1\|UDP-N-acetylmuramoylalanyl-D-glutamyl-2_6-diaminopimelate--D-alanyl-D-alanyl | WP_084049830.1 | 72.44 | 439 | 104 | 5 | 1 | 432 | 1 | 429 | 0 | 577 | 99 |

**REVERSE BLAST**

| | | | | | | | | | | | | |
|---|---|---|---|---|---|---|---|---|---|---|---|---|
| WP_084048503.1 | DR_0794/NP_294518.1 | 83.29 | 407 | 67 | 1 | 5 | 411 | 43 | 448 | 0 | 715 | 98 |
| WP_084048692.1 | DR_1674/NP_295397.1 | 85.24 | 332 | 49 | 0 | 1 | 332 | 1 | 332 | 0 | 589 | 99 |
| WP_084048755.1 | DR_1413/NP_295136.1 | 73.5 | 351 | 92 | 1 | 3 | 352 | 9 | 359 | 4.00E-179 | 503 | 98 |
| WP_084048988.1 | DR_1610/NP_295333.1 | 89.42 | 416 | 44 | 0 | 7 | 422 | 17 | 432 | 0 | 727 | 99 |
| WP_084048990.1 | DR_1614/NP_295337.1 | 87.04 | 162 | 21 | 0 | 1 | 162 | 1 | 162 | 4.00E-104 | 297 | 93 |
| WP_084048993.1 | DR_2194/NP_295916.1 | 87.85 | 288 | 30 | 1 | 1 | 283 | 8 | 295 | 8.00E-179 | 496 | 100 |
| WP_084049015.1 | DR_1365/NP_295088.1 | 81.47 | 464 | 86 | 0 | 5 | 468 | 10 | 473 | 0 | 691 | 99 |
| WP_084049092.1 | DR_1238/NP_294962.1 | 90.46 | 388 | 37 | 0 | 1 | 388 | 6 | 393 | 0 | 724 | 100 |
| WP_084049094.1 | DR_1758/NP_295481.1 | 72.03 | 379 | 106 | 0 | 1 | 379 | 1 | 379 | 0 | 540 | 98 |
| WP_084049365.1 | DR_1252/NP_294976.1 | 83.7 | 405 | 66 | 0 | 1 | 405 | 1 | 405 | 0 | 726 | 100 |
| WP_084049592.1 | DR_1278/NP_295002.1 | 76 | 325 | 73 | 1 | 1 | 320 | 1 | 325 | 9.00E-171 | 478 | 100 |
| WP_084049619.1 | DR_2008/NP_295731.1 | 83.68 | 337 | 55 | 0 | 1 | 337 | 1 | 337 | 0 | 566 | 99 |
| WP_084049830.1 | DR_0768/NP_294492.1 | 72.44 | 439 | 104 | 5 | 1 | 429 | 1 | 432 | 0 | 597 | 99 |
| WP_084050371.1 | DR_0671/NP_294394.1 | 63.23 | 378 | 128 | 2 | 6 | 372 | 3 | 380 | 5.00E-161 | 458 | 96 |
| WP_084050426.1 | DR_1588/NP_295311.1 | 72.5 | 400 | 103 | 3 | 15 | 409 | 22 | 419 | 0 | 578 | 96 |
| WP_084050599.1 | DR_0297/NP_294020.1 | 77.35 | 490 | 108 | 1 | 1 | 487 | 1 | 490 | 0 | 696 | 99 |
| WP_084050826.1 | DR_1420/NP_295143.2 | 90.23 | 266 | 26 | 0 | 1 | 266 | 1 | 266 | 3.00E-162 | 452 | 99 |
| WP_084050898.1 | WP_010887608.1 | 87.24 | 337 | 43 | 0 | 1 | 337 | 11 | 347 | 0 | 591 | 87 |

### *Deinococcus indicus*

| Query ID | Subject ID | % Identity | Alignment length | Mismatches | Gap opens | Query start | Query end | Sequence start | Sequence end | E-value | Bit score | Query coverage |
|---|---|---|---|---|---|---|---|---|---|---|---|---|
| DR_1238/NP_294962.1\|homocitrate | WP_088249070.1 | 90.885 | 384 | 35 | 0 | 10 | 393 | 12 | 395 | 0 | 729 | 98 |
| DR_1610/NP_295333.1\|3-isopropylmalate | WP_088247729.1 | 88.489 | 417 | 44 | 1 | 15 | 427 | 9 | 425 | 0 | 708 | 95 |
| DR_1614/NP_295337.1\|3-isopropylmalate | WP_088247727.1 | 90.123 | 162 | 16 | 0 | 1 | 162 | 1 | 162 | 2.92E-108 | 303 | 92 |
| DR_1674/NP_295397.1\|isocitrate | WP_088250045.1 | 90.39 | 333 | 32 | 0 | 1 | 333 | 1 | 333 | 0 | 621 | 100 |
| DR_1588/NP_295311.1\|class | WP_088250196.1 | 76.039 | 409 | 92 | 2 | 12 | 419 | 1 | 404 | 0 | 623 | 97 |
| DR_2194/NP_295916.1\|ribosomal | WP_088247722.1 | 88.85 | 287 | 32 | 0 | 8 | 294 | 1 | 287 | 0 | 507 | 97 |
| DR_1420/NP_295143.2\|acetylglutamate/acetylaminoadipate | WP_088247923.1 | 89.139 | 267 | 29 | 0 | 1 | 267 | 1 | 267 | 2.78E-165 | 455 | 100 |
| DR_0963/NP_294687.2\|argC;N-acetyl-gamma-glutamyl-phosphate | WP_088247541.1 | 86.628 | 344 | 46 | 0 | 4 | 347 | 5 | 348 | 0 | 595 | 99 |
| DR_0794/NP_294518.1\|acetylornithine | WP_088247362.1 | 79.767 | 430 | 83 | 2 | 37 | 465 | 1 | 427 | 0 | 680 | 92 |
| DR_1413/NP_295136.1\|acetyl-lysine | WP_088248220.1 | 70.083 | 361 | 102 | 3 | 3 | 362 | 8 | 363 | 4.06E-172 | 481 | 99 |
| DR_1252/NP_294976.1\|hypothetical | WP_088249699.1 | 89.877 | 405 | 41 | 0 | 1 | 405 | 1 | 405 | 0 | 771 | 100 |
| DR_1365/NP_295088.1\|aspartate | WP_088247627.1 | 82.759 | 464 | 80 | 0 | 10 | 473 | 5 | 468 | 0 | 735 | 98 |
| DR_2008/NP_295731.1\|aspartate-semialdehyde | WP_088247127.1 | 82.789 | 337 | 58 | 0 | 1 | 337 | 1 | 337 | 0 | 569 | 99 |
| DR_1278/NP_295002.1\|homoserine | WP_088249652.1 | 78.154 | 325 | 66 | 1 | 1 | 325 | 1 | 320 | 4.37E-178 | 492 | 100 |
| DR_0671/NP_294394.1\|class | WP_088246803.1 | 65.782 | 377 | 118 | 2 | 1 | 377 | 4 | 369 | 2.53E-149 | 424 | 99 |
| DR_1758/NP_295481.1\|diaminopimelate | WP_088249925.1 | 70.106 | 378 | 113 | 0 | 1 | 378 | 1 | 378 | 2.57E-178 | 498 | 99 |
| DR_0297/NP_294020.1\|UDP-N-acetylmuramoylalanyl-D-glutamate--2_6-diaminopimelate | WP_088248928.1 | 81.573 | 483 | 84 | 2 | 1 | 483 | 1 | 478 | 0 | 694 | 99 |
| DR_0768/NP_294492.1\|UDP-N-acetylmuramoylalanyl-D-glutamyl-2_6-diaminopimelate--D-alanyl-D-alanyl | WP_088249986.1 | 74.201 | 438 | 97 | 4 | 1 | 432 | 1 | 428 | 0 | 623 | 99 |

**REVERSE BLAST**

| | | | | | | | | | | | | |
|---|---|---|---|---|---|---|---|---|---|---|---|---|
| WP_088246803.1 | DR_0671/NP_294394.1 | 65.517 | 377 | 119 | 2 | 4 | 369 | 1 | 377 | 1.51E-160 | 452 | 98 |
| WP_088247127.1 | DR_2008/NP_295731.1 | 82.789 | 337 | 58 | 0 | 1 | 337 | 1 | 337 | 0 | 569 | 99 |
| WP_088247362.1 | DR_0794/NP_294518.1 | 79.767 | 430 | 83 | 2 | 1 | 427 | 37 | 465 | 0 | 714 | 100 |
| WP_088247541.1 | DR_0963/NP_294687.2 | 86.628 | 344 | 46 | 0 | 5 | 348 | 4 | 347 | 0 | 595 | 99 |
| WP_088247627.1 | DR_1365/NP_295088.1 | 82.759 | 464 | 80 | 0 | 5 | 468 | 10 | 473 | 0 | 719 | 99 |
| WP_088247722.1 | DR_2194/NP_295916.1 | 88.85 | 287 | 32 | 0 | 1 | 287 | 8 | 294 | 0 | 506 | 100 |
| WP_088247727.1 | DR_1614/NP_295337.1 | 90.123 | 162 | 16 | 0 | 1 | 162 | 1 | 162 | 2.09E-108 | 303 | 99 |
| WP_088247729.1 | DR_1610/NP_295333.1 | 88.489 | 417 | 44 | 1 | 9 | 425 | 15 | 427 | 0 | 708 | 97 |
| WP_088247923.1 | DR_1420/NP_295143.2 | 89.139 | 267 | 29 | 0 | 1 | 267 | 1 | 267 | 2.90E-165 | 455 | 100 |
| WP_088248220.1 | DR_1413/NP_295136.1 | 70.083 | 361 | 102 | 3 | 8 | 363 | 3 | 362 | 3.19E-172 | 481 | 97 |
| WP_088248928.1 | DR_0297/NP_294020.1 | 81.573 | 483 | 84 | 2 | 1 | 478 | 1 | 483 | 0 | 711 | 99 |
| WP_088249070.1 | DR_1238/NP_294962.1 | 90.885 | 384 | 35 | 0 | 12 | 395 | 10 | 393 | 0 | 729 | 97 |
| WP_088249652.1 | DR_1278/NP_295002.1 | 78.154 | 325 | 66 | 1 | 1 | 320 | 1 | 325 | 3.08E-178 | 492 | 100 |
| WP_088249699.1 | DR_1252/NP_294976.1 | 89.877 | 405 | 41 | 0 | 1 | 405 | 1 | 405 | 0 | 771 | 100 |
| WP_088249925.1 | DR_1758/NP_295481.1 | 70.106 | 378 | 113 | 0 | 1 | 378 | 1 | 378 | 0 | 522 | 99 |
| WP_088249986.1 | DR_0768/NP_294492.1 | 74.201 | 438 | 97 | 4 | 1 | 428 | 1 | 432 | 0 | 623 | 99 |
| WP_088250045.1 | DR_1674/NP_295397.1 | 90.39 | 333 | 32 | 0 | 1 | 333 | 1 | 333 | 0 | 621 | 100 |
| WP_088250196.1 | DR_1588/NP_295311.1 | 75.55 | 409 | 94 | 2 | 1 | 404 | 12 | 419 | 0 | 610 | 99 |

### *Deinococcus maricopensis*

| Query ID | Subject ID | % Identity | Alignment length | Mismatches | Gap opens | Query start | Query end | Sequence start | Sequence end | E-value | Bit score | Query coverage |
|---|---|---|---|---|---|---|---|---|---|---|---|---|
| DR_1238/NP_294962.1\|homocitrate | WP_013556513.1 | 82.73 | 388 | 62 | 3 | 6 | 393 | 1 | 383 | 0 | 654 | 99 |
| DR_1610/NP_295333.1\|3-isopropylmalate | WP_013556510.1 | 88.05 | 410 | 47 | 1 | 18 | 427 | 3 | 410 | 0 | 686 | 94 |
| DR_1614/NP_295337.1\|3-isopropylmalate | WP_013556508.1 | 83.95 | 162 | 26 | 0 | 1 | 162 | 1 | 162 | 3.00E-99 | 285 | 92 |
| DR_1674/NP_295397.1\|isocitrate | WP_013556699.1 | 80.18 | 333 | 64 | 2 | 1 | 332 | 1 | 332 | 0 | 555 | 99 |
| DR_1588/NP_295311.1\|class | WP_013555215.1 | 68.83 | 401 | 117 | 4 | 22 | 419 | 8 | 403 | 0 | 548 | 95 |
| DR_2194/NP_295916.1\|ribosomal | WP_013556505.1 | 83.28 | 287 | 39 | 2 | 8 | 294 | 1 | 278 | 1.00E-175 | 488 | 97 |
| DR_1420/NP_295143.2\|acetylglutamate/acetylaminoadipate | WP_013556501.1 | 88.72 | 266 | 30 | 0 | 1 | 266 | 1 | 266 | 3.00E-161 | 449 | 99 |
| DR_0963/NP_294687.2\|argC;N-acetyl-gamma-glutamyl-phosphate | WP_013556504.1 | 81.16 | 345 | 65 | 0 | 3 | 347 | 2 | 346 | 0 | 596 | 99 |
| DR_0794/NP_294518.1\|acetylornithine | WP_013557594.1 | 78.55 | 415 | 85 | 2 | 37 | 450 | 1 | 412 | 0 | 671 | 89 |
| DR_1413/NP_295136.1\|acetyl-lysine | WP_013557730.1 | 70.03 | 347 | 99 | 1 | 14 | 360 | 18 | 359 | 5.00E-166 | 469 | 96 |
| DR_1252/NP_294976.1\|hypothetical | WP_013556067.1 | 83.21 | 405 | 68 | 0 | 1 | 405 | 1 | 405 | 0 | 716 | 100 |
| DR_1365/NP_295088.1\|aspartate | WP_013556975.1 | 66.52 | 463 | 152 | 1 | 10 | 469 | 1 | 463 | 0 | 542 | 97 |
| DR_2008/NP_295731.1\|aspartate-semialdehyde | WP_013557815.1 | 80.48 | 333 | 65 | 0 | 1 | 333 | 1 | 333 | 0 | 538 | 99 |
| DR_1278/NP_295002.1\|homoserine | WP_013556614.1 | 73.23 | 325 | 82 | 1 | 1 | 325 | 1 | 320 | 6.00E-165 | 464 | 100 |
| DR_0671/NP_294394.1\|class | WP_013558211.1 | 32.05 | 337 | 200 | 7 | 54 | 378 | 65 | 384 | 2.00E-40 | 146 | 85 |
| DR_1758/NP_295481.1\|diaminopimelate | WP_013557797.1 | 66.13 | 372 | 125 | 1 | 8 | 379 | 40 | 410 | 1.00E-166 | 474 | 98 |
| DR_0297/NP_294020.1\|UDP-N-acetylmuramoylalanyl-D-glutamate--2_6-diaminopimelate | WP_013557958.1 | 74.08 | 490 | 122 | 2 | 1 | 490 | 1 | 485 | 0 | 667 | 100 |
| DR_0768/NP_294492.1\|UDP-N-acetylmuramoylalanyl-D-glutamyl-2_6-diaminopimelate--D-alanyl-D-alanyl | WP_013556808.1 | 72.16 | 431 | 110 | 2 | 1 | 431 | 1 | 421 | 0 | 563 | 99 |

**REVERSE BLAST**

| | | | | | | | | | | | | |
|---|---|---|---|---|---|---|---|---|---|---|---|---|
| WP_013555215.1 | DR_1588/NP_295311.1 | 68.83 | 401 | 117 | 4 | 8 | 403 | 22 | 419 | 0 | 541 | 98 |
| WP_013556067.1 | DR_1252/NP_294976.1 | 83.21 | 405 | 68 | 0 | 1 | 405 | 1 | 405 | 0 | 716 | 100 |
| WP_013556501.1 | DR_1420/NP_295143.2 | 88.72 | 266 | 30 | 0 | 1 | 266 | 1 | 266 | 2.00E-161 | 450 | 99 |
| WP_013556504.1 | DR_0963/NP_294687.2 | 81.16 | 345 | 65 | 0 | 2 | 346 | 3 | 347 | 0 | 572 | 99 |
| WP_013556505.1 | DR_2194/NP_295916.1 | 83.28 | 287 | 39 | 2 | 1 | 278 | 8 | 294 | 3.00E-169 | 471 | 99 |
| WP_013556508.1 | DR_1614/NP_295337.1 | 83.95 | 162 | 26 | 0 | 1 | 162 | 1 | 162 | 3.00E-99 | 285 | 93 |
| WP_013556510.1 | DR_1610/NP_295333.1 | 88.05 | 410 | 47 | 1 | 3 | 410 | 18 | 427 | 0 | 686 | 98 |
| WP_013556513.1 | DR_1238/NP_294962.1 | 82.73 | 388 | 62 | 3 | 1 | 383 | 6 | 393 | 0 | 654 | 100 |
| WP_013556614.1 | DR_1278/NP_295002.1 | 73.23 | 325 | 82 | 1 | 1 | 320 | 1 | 325 | 4.00E-166 | 466 | 100 |
| WP_013556699.1 | DR_1674/NP_295397.1 | 80.18 | 333 | 64 | 2 | 1 | 332 | 1 | 332 | 0 | 555 | 99 |
| WP_013556808.1 | DR_0768/NP_294492.1 | 72.16 | 431 | 110 | 2 | 1 | 421 | 1 | 431 | 0 | 600 | 99 |
| WP_013557594.1 | DR_0794/NP_294518.1 | 78.55 | 415 | 85 | 2 | 1 | 412 | 37 | 450 | 0 | 671 | 100 |
| WP_013557730.1 | DR_1413/NP_295136.1 | 70.09 | 351 | 100 | 1 | 14 | 359 | 10 | 360 | 1.00E-167 | 474 | 96 |
| WP_013557797.1 | DR_1758/NP_295481.1 | 66.13 | 372 | 125 | 1 | 40 | 410 | 8 | 379 | 1.00E-170 | 484 | 90 |
| WP_013557815.1 | DR_2008/NP_295731.1 | 80.48 | 333 | 65 | 0 | 1 | 333 | 1 | 333 | 0 | 538 | 99 |
| WP_013557958.1 | DR_0297/NP_294020.1 | 74.08 | 490 | 122 | 2 | 1 | 485 | 1 | 490 | 0 | 652 | 99 |
| WP_013558211.1 | NP_294346.1 | 72.7 | 381 | 104 | 0 | 6 | 386 | 7 | 387 | 0 | 565 | 98 |
| WP_013556975.1 | WP_010888006.1 | 66.523 | 463 | 152 | 1 | 1 | 463 | 10 | 469 | 0 | 555 | 67 |

### *Deinococcus marmoris*

| Query ID | Subject ID | % Identity | Alignment length | Mismatches | Gap opens | Query start | Query end | Sequence start | Sequence end | E-value | Bit score | Query coverage |
|---|---|---|---|---|---|---|---|---|---|---|---|---|
| DR_1238/NP_294962.1\|homocitrate | WP_029482933.1 | 89.74 | 380 | 39 | 0 | 14 | 393 | 12 | 391 | 0 | 714 | 97 |
| DR_1610/NP_295333.1\|3-isopropylmalate | WP_081846195.1 | 87.62 | 412 | 49 | 1 | 16 | 427 | 4 | 413 | 0 | 699 | 95 |
| DR_1614/NP_295337.1\|3-isopropylmalate | WP_029480722.1 | 85.8 | 169 | 24 | 0 | 1 | 169 | 1 | 169 | 7.00E-106 | 301 | 95 |
| DR_1674/NP_295397.1\|isocitrate | WP_029481733.1 | 88.25 | 332 | 39 | 0 | 1 | 332 | 1 | 332 | 0 | 608 | 99 |
| DR_1588/NP_295311.1\|class | WP_081846087.1 | 76.23 | 387 | 86 | 2 | 34 | 419 | 1 | 382 | 0 | 596 | 92 |
| DR_2194/NP_295916.1\|ribosomal | WP_029480719.1 | 84.03 | 288 | 46 | 0 | 8 | 295 | 1 | 288 | 0 | 506 | 98 |
| DR_1420/NP_295143.2\|acetylglutamate/acetylaminoadipate | WP_029482428.1 | 91.76 | 267 | 22 | 0 | 1 | 267 | 1 | 267 | 4.00E-169 | 470 | 100 |
| DR_0963/NP_294687.2\|argC;N-acetyl-gamma-glutamyl-phosphate | WP_029480901.1 | 86.99 | 346 | 45 | 0 | 2 | 347 | 5 | 350 | 0 | 605 | 99 |
| DR_0794/NP_294518.1\|acetylornithine | WP_029479832.1 | 76.03 | 438 | 84 | 1 | 34 | 450 | 1 | 438 | 0 | 693 | 90 |
| DR_1413/NP_295136.1\|acetyl-lysine | WP_034414216.1 | 72.42 | 359 | 93 | 3 | 3 | 359 | 10 | 364 | 7.00E-179 | 503 | 99 |
| DR_1252/NP_294976.1\|hypothetical | WP_029483567.1 | 85.15 | 404 | 60 | 0 | 1 | 404 | 1 | 404 | 0 | 734 | 99 |
| DR_1365/NP_295088.1\|aspartate | WP_029481736.1 | 83.62 | 458 | 75 | 0 | 10 | 467 | 5 | 462 | 0 | 708 | 97 |
| DR_2008/NP_295731.1\|aspartate-semialdehyde | WP_029483346.1 | 84.73 | 334 | 51 | 0 | 1 | 334 | 1 | 334 | 0 | 585 | 99 |
| DR_1278/NP_295002.1\|homoserine | WP_029481113.1 | 78.46 | 325 | 65 | 1 | 1 | 325 | 1 | 320 | 0 | 518 | 100 |
| DR_0671/NP_294394.1\|class | WP_029481230.1 | 67.37 | 377 | 116 | 3 | 3 | 379 | 6 | 375 | 8.00E-173 | 489 | 99 |
| DR_1758/NP_295481.1\|diaminopimelate | WP_029483307.1 | 70.31 | 384 | 109 | 1 | 1 | 379 | 1 | 384 | 2.00E-180 | 508 | 100 |
| DR_0297/NP_294020.1\|UDP-N-acetylmuramoylalanyl-D-glutamate--2_6-diaminopimelate | WP_029482980.1 | 78.18 | 472 | 98 | 1 | 22 | 488 | 19 | 490 | 0 | 674 | 95 |
| DR_0768/NP_294492.1\|UDP-N-acetylmuramoylalanyl-D-glutamyl-2_6-diaminopimelate--D-alanyl-D-alanyl | WP_029479737.1 | 73.04 | 434 | 106 | 3 | 1 | 433 | 1 | 424 | 0 | 582 | 100 |

**REVERSE BLAST**

| | | | | | | | | | | | | |
|---|---|---|---|---|---|---|---|---|---|---|---|---|
| WP_029479737.1 | DR_0768/NP_294492.1 | 73.04 | 434 | 106 | 3 | 1 | 424 | 1 | 433 | 0 | 602 | 97 |
| WP_029479832.1 | DR_0794/NP_294518.1 | 76.03 | 438 | 84 | 1 | 1 | 438 | 34 | 450 | 0 | 693 | 99 |
| WP_029480719.1 | DR_2194/NP_295916.1 | 84.03 | 288 | 46 | 0 | 1 | 288 | 8 | 295 | 5.00E-177 | 491 | 100 |
| WP_029480722.1 | DR_1614/NP_295337.1 | 85.8 | 169 | 24 | 0 | 1 | 169 | 1 | 169 | 5.00E-106 | 301 | 99 |
| WP_029480901.1 | DR_0963/NP_294687.2 | 86.99 | 346 | 45 | 0 | 5 | 350 | 2 | 347 | 0 | 606 | 99 |
| WP_029481113.1 | DR_1278/NP_295002.1 | 78.46 | 325 | 65 | 1 | 1 | 320 | 1 | 325 | 4.00E-180 | 502 | 100 |
| WP_029481230.1 | DR_0671/NP_294394.1 | 66.75 | 379 | 119 | 3 | 4 | 375 | 1 | 379 | 3.00E-165 | 469 | 96 |
| WP_029481733.1 | DR_1674/NP_295397.1 | 88.25 | 332 | 39 | 0 | 1 | 332 | 1 | 332 | 0 | 608 | 100 |
| WP_029481736.1 | DR_1365/NP_295088.1 | 83.62 | 458 | 75 | 0 | 5 | 462 | 10 | 467 | 0 | 708 | 97 |
| WP_029482428.1 | DR_1420/NP_295143.2 | 91.76 | 267 | 22 | 0 | 1 | 267 | 1 | 267 | 2.00E-169 | 470 | 100 |
| WP_029482933.1 | DR_1238/NP_294962.1 | 88.89 | 387 | 43 | 0 | 5 | 391 | 7 | 393 | 0 | 719 | 99 |
| WP_029482980.1 | DR_0297/NP_294020.1 | 76.88 | 493 | 106 | 2 | 1 | 490 | 1 | 488 | 0 | 680 | 98 |
| WP_029483307.1 | DR_1758/NP_295481.1 | 70.31 | 384 | 109 | 1 | 1 | 384 | 1 | 379 | 0 | 541 | 100 |
| WP_029483346.1 | DR_2008/NP_295731.1 | 84.73 | 334 | 51 | 0 | 1 | 334 | 1 | 334 | 0 | 585 | 99 |
| WP_029483567.1 | DR_1252/NP_294976.1 | 85.15 | 404 | 60 | 0 | 1 | 404 | 1 | 404 | 0 | 734 | 100 |
| WP_034414216.1 | DR_1413/NP_295136.1 | 72.42 | 359 | 93 | 3 | 10 | 364 | 3 | 359 | 6.00E-179 | 503 | 95 |
| WP_081846087.1 | DR_1588/NP_295311.1 | 76.23 | 387 | 86 | 2 | 1 | 382 | 34 | 419 | 0 | 588 | 99 |
| WP_081846195.1 | DR_1610/NP_295333.1 | 87.62 | 412 | 49 | 1 | 4 | 413 | 16 | 427 | 0 | 699 | 99 |

### *Deinococcus misasensis*

| Query ID | Subject ID | % Identity | Alignment length | Mismatches | Gap opens | Query start | Query end | Sequence start | Sequence end | E-value | Bit score | Query coverage |
|---|---|---|---|---|---|---|---|---|---|---|---|---|
| DR_1238/NP_294962.1\|homocitrate | WP_051964282.1 | 76.84 | 380 | 84 | 2 | 14 | 393 | 14 | 389 | 0 | 614 | 97 |
| DR_1610/NP_295333.1\|3-isopropylmalate | WP_051964321.1 | 80.59 | 407 | 79 | 0 | 21 | 427 | 1 | 407 | 0 | 660 | 94 |
| DR_1614/NP_295337.1\|3-isopropylmalate | WP_034341311.1 | 72.39 | 163 | 44 | 1 | 1 | 162 | 1 | 163 | 5.00E-85 | 249 | 92 |
| DR_1674/NP_295397.1\|isocitrate | WP_034334830.1 | 73.19 | 332 | 88 | 1 | 1 | 332 | 1 | 331 | 1.00E-179 | 502 | 99 |
| DR_1588/NP_295311.1\|class | WP_034341490.1 | 58.94 | 397 | 155 | 4 | 26 | 419 | 11 | 402 | 1.00E-165 | 473 | 94 |
| DR_2194/NP_295916.1\|ribosomal | WP_034341313.1 | 78.6 | 285 | 53 | 2 | 8 | 292 | 1 | 277 | 1.00E-168 | 471 | 97 |
| DR_1420/NP_295143.2\|acetylglutamate/acetylaminoadipate | WP_034341330.1 | 83.15 | 267 | 45 | 0 | 1 | 267 | 1 | 267 | 2.00E-157 | 440 | 100 |
| DR_0963/NP_294687.2\|argC;N-acetyl-gamma-glutamyl-phosphate | WP_034341317.1 | 73.18 | 343 | 92 | 0 | 5 | 347 | 3 | 345 | 0 | 511 | 99 |
| DR_0794/NP_294518.1\|acetylornithine | WP_051964295.1 | 74.36 | 390 | 98 | 2 | 46 | 434 | 7 | 395 | 0 | 606 | 84 |
| DR_1413/NP_295136.1\|acetyl-lysine | WP_034341940.1 | 56.98 | 351 | 147 | 4 | 12 | 361 | 4 | 351 | 4.00E-139 | 401 | 97 |
| DR_1252/NP_294976.1\|hypothetical | WP_034345194.1 | 77.23 | 404 | 91 | 1 | 1 | 404 | 1 | 403 | 0 | 671 | 99 |
| DR_1365/NP_295088.1\|aspartate | WP_034335210.1 | 63.07 | 463 | 165 | 3 | 10 | 469 | 1 | 460 | 0 | 582 | 97 |
| DR_2008/NP_295731.1\|aspartate-semialdehyde | WP_034346361.1 | 75.98 | 333 | 80 | 0 | 1 | 333 | 1 | 333 | 0 | 530 | 99 |
| DR_1278/NP_295002.1\|homoserine | WP_051964544.1 | 54.29 | 326 | 141 | 3 | 1 | 324 | 1 | 320 | 2.00E-109 | 323 | 99 |
| DR_0671/NP_294394.1\|class | WP_034339242.1 | 41.54 | 390 | 200 | 7 | 1 | 375 | 4 | 380 | 6.00E-83 | 259 | 98 |
| DR_1758/NP_295481.1\|diaminopimelate | WP_034338895.1 | 58.78 | 376 | 154 | 1 | 3 | 378 | 2 | 376 | 3.00E-150 | 431 | 99 |
| DR_0297/NP_294020.1\|UDP-N-acetylmuramoylalanyl-D-glutamate--2_6-diaminopimelate | WP_034346290.1 | 62.86 | 490 | 171 | 5 | 1 | 490 | 1 | 479 | 0 | 612 | 100 |
| DR_0768/NP_294492.1\|UDP-N-acetylmuramoylalanyl-D-glutamyl-2_6-diaminopimelate--D-alanyl-D-alanyl | WP_051963587.1 | 49.28 | 418 | 200 | 5 | 1 | 415 | 1 | 409 | 1.00E-129 | 382 | 96 |

**REVERSE BLAST**

| | | | | | | | | | | | | |
|---|---|---|---|---|---|---|---|---|---|---|---|---|
| WP_034334830.1 | DR_1674/NP_295397.1 | 73.19 | 332 | 88 | 1 | 1 | 331 | 1 | 332 | 6.00E-180 | 502 | 99 |
| WP_034335210.1 | DR_1365/NP_295088.1 | 63.07 | 463 | 165 | 3 | 1 | 460 | 10 | 469 | 0 | 537 | 99 |
| WP_034338895.1 | DR_1758/NP_295481.1 | 58.78 | 376 | 154 | 1 | 2 | 376 | 3 | 378 | 2.00E-150 | 431 | 99 |
| WP_034339242.1 | DR_0671/NP_294394.1 | 41.98 | 393 | 194 | 8 | 4 | 380 | 1 | 375 | 2.00E-79 | 249 | 97 |
| WP_034341311.1 | DR_1614/NP_295337.1 | 72.39 | 163 | 44 | 1 | 1 | 163 | 1 | 162 | 3.00E-85 | 249 | 100 |
| WP_034341313.1 | DR_2194/NP_295916.1 | 78.6 | 285 | 53 | 2 | 1 | 277 | 8 | 292 | 4.00E-163 | 456 | 99 |
| WP_034341317.1 | DR_0963/NP_294687.2 | 73.18 | 343 | 92 | 0 | 3 | 345 | 5 | 347 | 0 | 511 | 99 |
| WP_034341330.1 | DR_1420/NP_295143.2 | 83.15 | 267 | 45 | 0 | 1 | 267 | 1 | 267 | 2.00E-154 | 432 | 97 |
| WP_034341490.1 | DR_1588/NP_295311.1 | 57.93 | 397 | 159 | 4 | 11 | 402 | 26 | 419 | 1.00E-162 | 464 | 98 |
| WP_034341940.1 | DR_1413/NP_295136.1 | 56.98 | 351 | 147 | 4 | 4 | 351 | 12 | 361 | 3.00E-139 | 401 | 99 |
| WP_034345194.1 | DR_1252/NP_294976.1 | 77.23 | 404 | 91 | 1 | 1 | 403 | 1 | 404 | 0 | 671 | 100 |
| WP_034346290.1 | DR_0297/NP_294020.1 | 62.86 | 490 | 171 | 5 | 1 | 479 | 1 | 490 | 0 | 565 | 99 |
| WP_034346361.1 | DR_2008/NP_295731.1 | 75.98 | 333 | 80 | 0 | 1 | 333 | 1 | 333 | 0 | 530 | 98 |
| WP_051963587.1 | DR_0768/NP_294492.1 | 49.28 | 418 | 200 | 5 | 1 | 409 | 1 | 415 | 7.00E-130 | 382 | 99 |
| WP_051964282.1 | DR_1238/NP_294962.1 | 76.84 | 380 | 84 | 2 | 14 | 389 | 14 | 393 | 0 | 614 | 97 |
| WP_051964295.1 | DR_0794/NP_294518.1 | 74.36 | 390 | 98 | 2 | 7 | 395 | 46 | 434 | 0 | 606 | 98 |
| WP_051964321.1 | DR_1610/NP_295333.1 | 80.59 | 407 | 79 | 0 | 1 | 407 | 21 | 427 | 0 | 639 | 100 |
| WP_051964544.1 | DR_1278/NP_295002.1 | 54.29 | 326 | 141 | 3 | 1 | 320 | 1 | 324 | 3.00E-111 | 327 | 94 |

### *Deinococcus murrayi*

| Query ID | Subject ID | % Identity | Alignment length | Mismatches | Gap opens | Query start | Query end | Sequence start | Sequence end | E-value | Bit score | Query coverage |
|---|---|---|---|---|---|---|---|---|---|---|---|---|
| DR_1238/NP_294962.1\|homocitrate | WP_084542982.1 | 51.35 | 370 | 175 | 3 | 15 | 384 | 5 | 369 | 1.00E-122 | 361 | 94 |
| DR_1610/NP_295333.1\|3-isopropylmalate | WP_034403848.1 | 56.03 | 423 | 179 | 4 | 15 | 432 | 2 | 422 | 3.00E-149 | 432 | 96 |
| DR_1614/NP_295337.1\|3-isopropylmalate | WP_051363597.1 | 55.97 | 159 | 69 | 1 | 3 | 161 | 17 | 174 | 2.00E-61 | 189 | 90 |
| DR_1674/NP_295397.1\|isocitrate | WP_027460381.1 | 87.05 | 332 | 43 | 0 | 1 | 332 | 1 | 332 | 0 | 601 | 99 |
| DR_1588/NP_295311.1\|class | WP_027459826.1 | 72.48 | 407 | 104 | 4 | 16 | 419 | 6 | 407 | 0 | 584 | 96 |
| DR_2194/NP_295916.1\|ribosomal | WP_034409500.1 | 51.61 | 279 | 127 | 2 | 11 | 289 | 9 | 279 | 6.00E-91 | 273 | 95 |
| DR_1420/NP_295143.2\|acetylglutamate/acetylaminoadipate | WP_051363596.1 | 55.64 | 266 | 116 | 2 | 2 | 266 | 11 | 275 | 2.00E-76 | 234 | 99 |
| DR_0963/NP_294687.2\|argC;N-acetyl-gamma-glutamyl-phosphate | WP_027461253.1 | 49.71 | 342 | 169 | 2 | 8 | 348 | 4 | 343 | 5.00E-102 | 305 | 98 |
| DR_0794/NP_294518.1\|acetylornithine | WP_027461022.1 | 80.93 | 409 | 77 | 1 | 37 | 445 | 1 | 408 | 0 | 690 | 88 |
| DR_1413/NP_295136.1\|acetyl-lysine | WP_027461251.1 | 45.51 | 356 | 181 | 4 | 9 | 359 | 3 | 350 | 3.00E-77 | 242 | 97 |
| DR_1252/NP_294976.1\|hypothetical | WP_027461322.1 | 83.5 | 406 | 66 | 1 | 1 | 405 | 1 | 406 | 0 | 728 | 100 |
| DR_1365/NP_295088.1\|aspartate | WP_027460751.1 | 81.96 | 460 | 83 | 0 | 10 | 469 | 5 | 464 | 0 | 643 | 97 |
| DR_2008/NP_295731.1\|aspartate-semialdehyde | WP_027460069.1 | 81.74 | 334 | 61 | 0 | 1 | 334 | 1 | 334 | 0 | 548 | 99 |
| DR_1278/NP_295002.1\|homoserine | WP_084542893.1 | 75 | 324 | 76 | 1 | 2 | 325 | 43 | 361 | 1.00E-171 | 482 | 99 |
| DR_0671/NP_294394.1\|class | WP_027459459.1 | 34.43 | 334 | 190 | 7 | 54 | 375 | 66 | 382 | 4.00E-39 | 142 | 85 |
| DR_1758/NP_295481.1\|diaminopimelate | WP_027461283.1 | 70.56 | 377 | 110 | 1 | 3 | 378 | 2 | 378 | 0 | 524 | 99 |
| DR_0297/NP_294020.1\|UDP-N-acetylmuramoylalanyl-D-glutamate--2,6-diaminopimelate | WP_027460570.1 | 76.46 | 480 | 110 | 1 | 1 | 480 | 1 | 477 | 0 | 679 | 98 |
| DR_0768/NP_294492.1\|UDP-N-acetylmuramoylalanyl-D-glutamyl-2_6-diaminopimelate--D-alanyl-D-alanyl | WP_027459520.1 | 71.75 | 439 | 107 | 5 | 1 | 432 | 1 | 429 | 0 | 560 | 99 |

**REVERSE BLAST**

| | | | | | | | | | | | | |
|---|---|---|---|---|---|---|---|---|---|---|---|---|
| WP_027459459.1 | NP_294346.1 | 73.39 | 387 | 103 | 0 | 1 | 387 | 1 | 387 | 0 | 578 | 99 |
| WP_027459520.1 | DR_0768/NP_294492.1 | 71.75 | 439 | 107 | 5 | 1 | 429 | 1 | 432 | 0 | 598 | 99 |
| WP_027459826.1 | DR_1588/NP_295311.1 | 72.24 | 407 | 105 | 4 | 6 | 407 | 16 | 419 | 0 | 577 | 98 |
| WP_027460069.1 | DR_2008/NP_295731.1 | 81.74 | 334 | 61 | 0 | 1 | 334 | 1 | 334 | 0 | 567 | 99 |
| WP_027460381.1 | DR_1674/NP_295397.1 | 87.05 | 332 | 43 | 0 | 1 | 332 | 1 | 332 | 0 | 601 | 100 |
| WP_027460570.1 | DR_0297/NP_294020.1 | 76.33 | 490 | 113 | 1 | 1 | 487 | 1 | 490 | 0 | 681 | 99 |
| WP_027460751.1 | DR_1365/NP_295088.1 | 81.96 | 460 | 83 | 0 | 5 | 464 | 10 | 469 | 0 | 691 | 98 |
| WP_027461022.1 | DR_0794/NP_294518.1 | 80.93 | 409 | 77 | 1 | 1 | 408 | 37 | 445 | 0 | 690 | 99 |
| WP_027461251.1 | DR_1413/NP_295136.1 | 45.51 | 356 | 181 | 4 | 3 | 350 | 9 | 359 | 9.00E-87 | 266 | 98 |
| WP_027461253.1 | DR_0963/NP_294687.2 | 49.71 | 342 | 169 | 2 | 4 | 343 | 8 | 348 | 3.00E-102 | 305 | 99 |
| WP_027461283.1 | DR_1758/NP_295481.1 | 70.56 | 377 | 110 | 1 | 2 | 378 | 3 | 378 | 0 | 524 | 99 |
| WP_027461322.1 | DR_1252/NP_294976.1 | 83.5 | 406 | 66 | 1 | 1 | 406 | 1 | 405 | 0 | 728 | 100 |
| WP_034403848.1 | DR_1610/NP_295333.1 | 56.03 | 423 | 179 | 4 | 2 | 422 | 15 | 432 | 1.00E-144 | 420 | 99 |
| WP_034409500.1 | DR_2194/NP_295916.1 | 51.42 | 282 | 129 | 2 | 9 | 282 | 11 | 292 | 7.00E-89 | 268 | 92 |
| WP_051363596.1 | DR_1420/NP_295143.2 | 55.64 | 266 | 116 | 2 | 11 | 275 | 2 | 266 | 1.00E-83 | 253 | 92 |
| WP_051363597.1 | DR_1614/NP_295337.1 | 55.97 | 159 | 69 | 1 | 17 | 174 | 3 | 161 | 2.00E-61 | 189 | 89 |
| WP_084542893.1 | DR_1278/NP_295002.1 | 75 | 324 | 76 | 1 | 43 | 361 | 2 | 325 | 1.00E-171 | 482 | 88 |
| WP_084542982.1 | DR_1238/NP_294962.1 | 51.35 | 370 | 175 | 3 | 5 | 369 | 15 | 384 | 5.00E-132 | 385 | 91 |

### *Deinococcus peraridilitoris*

| Query ID | Subject ID | % Identity | Alignment length | Mismatches | Gap opens | Query start | Query end | Sequence start | Sequence end | E-value | Bit score | Query coverage |
|---|---|---|---|---|---|---|---|---|---|---|---|---|
| DR_1238/NP_294962.1\|homocitrate | WP_083865818.1 | 78.53 | 382 | 78 | 2 | 12 | 393 | 33 | 410 | 0 | 629 | 97 |
| DR_1610/NP_295333.1\|3-isopropylmalate | WP_015236530.1 | 84.78 | 414 | 62 | 1 | 16 | 429 | 6 | 418 | 0 | 706 | 95 |
| DR_1614/NP_295337.1\|3-isopropylmalate | WP_015236528.1 | 74.55 | 165 | 41 | 1 | 1 | 164 | 1 | 165 | 6.00E-86 | 251 | 93 |
| DR_1674/NP_295397.1\|isocitrate | WP_015237531.1 | 66.67 | 330 | 108 | 2 | 1 | 329 | 1 | 329 | 4.00E-160 | 452 | 99 |
| DR_1588/NP_295311.1\|class | WP_015236637.1 | 43.22 | 398 | 214 | 5 | 26 | 414 | 4 | 398 | 2.00E-101 | 308 | 93 |
| DR_2194/NP_295916.1\|ribosomal | WP_015236525.1 | 75.26 | 287 | 62 | 2 | 8 | 294 | 1 | 278 | 3.00E-157 | 441 | 97 |
| DR_1420/NP_295143.2\|acetylglutamate/acetylaminoadipate | WP_041231552.1 | 87.64 | 267 | 33 | 0 | 1 | 267 | 1 | 267 | 3.00E-172 | 478 | 100 |
| DR_0963/NP_294687.2\|argC;N-acetyl-gamma-glutamyl-phosphate | WP_015236524.1 | 76.18 | 340 | 81 | 0 | 8 | 347 | 6 | 345 | 0 | 528 | 98 |
| DR_0794/NP_294518.1\|acetylornithine | WP_015236501.1 | 76.62 | 402 | 91 | 1 | 37 | 438 | 1 | 399 | 0 | 643 | 86 |
| DR_1413/NP_295136.1\|acetyl-lysine | WP_015234424.1 | 64.46 | 377 | 112 | 4 | 1 | 361 | 1 | 371 | 2.00E-160 | 456 | 99 |
| DR_1252/NP_294976.1\|hypothetical | WP_015237324.1 | 80.74 | 405 | 78 | 0 | 1 | 405 | 1 | 405 | 0 | 707 | 100 |
| DR_1365/NP_295088.1\|aspartate | WP_015237416.1 | 60.91 | 463 | 174 | 2 | 10 | 467 | 1 | 461 | 2.00E-172 | 495 | 97 |
| DR_2008/NP_295731.1\|aspartate-semialdehyde | WP_015234775.1 | 73.57 | 333 | 88 | 0 | 1 | 333 | 1 | 333 | 2.00E-180 | 504 | 99 |
| DR_1278/NP_295002.1\|homoserine | WP_015234086.1 | 54.01 | 324 | 144 | 1 | 1 | 324 | 1 | 319 | 3.00E-117 | 342 | 99 |
| DR_0671/NP_294394.1\|class | WP_041230848.1 | 43 | 393 | 185 | 11 | 11 | 378 | 14 | 392 | 4.00E-78 | 246 | 97 |
| DR_1758/NP_295481.1\|diaminopimelate | WP_015236600.1 | 62.92 | 383 | 135 | 3 | 3 | 379 | 23 | 404 | 3.00E-157 | 450 | 99 |
| DR_0297/NP_294020.1\|UDP-N-acetylmuramoylalanyl-D-glutamate--2_6-diaminopimelate | WP_015236459.1 | 70.29 | 488 | 134 | 5 | 1 | 488 | 1 | 477 | 0 | 628 | 99 |
| DR_0768/NP_294492.1\|UDP-N-acetylmuramoylalanyl-D-glutamyl-2_6-diaminopimelate--D-alanyl-D-alanyl | WP_015234106.1 | 61.66 | 433 | 152 | 5 | 1 | 429 | 3 | 425 | 8.00E-172 | 490 | 99 |

**REVERSE BLAST**

| | | | | | | | | | | | | |
|---|---|---|---|---|---|---|---|---|---|---|---|---|
| WP_015234086.1 | DR_1278/NP_295002.1 | 54.01 | 324 | 144 | 1 | 1 | 319 | 1 | 324 | 4.00E-113 | 332 | 99 |
| WP_015234106.1 | DR_0768/NP_294492.1 | 61.66 | 433 | 152 | 5 | 3 | 425 | 1 | 429 | 6.00E-172 | 490 | 97 |
| WP_015234424.1 | DR_1413/NP_295136.1 | 64.46 | 377 | 112 | 4 | 1 | 371 | 1 | 361 | 2.00E-160 | 456 | 99 |
| WP_015234775.1 | DR_2008/NP_295731.1 | 73.57 | 333 | 88 | 0 | 1 | 333 | 1 | 333 | 2.00E-180 | 504 | 99 |
| WP_015236459.1 | DR_0297/NP_294020.1 | 70.29 | 488 | 134 | 5 | 1 | 477 | 1 | 488 | 0 | 599 | 99 |
| WP_015236501.1 | DR_0794/NP_294518.1 | 76.62 | 402 | 91 | 1 | 1 | 399 | 37 | 438 | 0 | 643 | 97 |
| WP_015236524.1 | DR_0963/NP_294687.2 | 76.18 | 340 | 81 | 0 | 6 | 345 | 8 | 347 | 0 | 528 | 98 |
| WP_015236525.1 | DR_2194/NP_295916.1 | 75.26 | 287 | 62 | 2 | 1 | 278 | 8 | 294 | 1.00E-151 | 427 | 99 |
| WP_015236528.1 | DR_1614/NP_295337.1 | 74.55 | 165 | 41 | 1 | 1 | 165 | 1 | 164 | 4.00E-86 | 251 | 99 |
| WP_015236530.1 | DR_1610/NP_295333.1 | 85.19 | 412 | 60 | 1 | 6 | 416 | 16 | 427 | 0 | 679 | 97 |
| WP_015236600.1 | DR_1758/NP_295481.1 | 62.92 | 383 | 135 | 3 | 23 | 404 | 3 | 379 | 2.00E-160 | 458 | 92 |
| WP_015236637.1 | DR_1588/NP_295311.1 | 43.22 | 398 | 214 | 5 | 4 | 398 | 26 | 414 | 2.00E-99 | 303 | 99 |
| WP_015237324.1 | DR_1252/NP_294976.1 | 80.74 | 405 | 78 | 0 | 1 | 405 | 1 | 405 | 0 | 707 | 100 |
| WP_015237416.1 | DR_1365/NP_295088.1 | 60.91 | 463 | 174 | 2 | 1 | 461 | 10 | 467 | 7.00E-180 | 513 | 99 |
| WP_015237531.1 | DR_1674/NP_295397.1 | 66.67 | 330 | 108 | 2 | 1 | 329 | 1 | 329 | 3.00E-160 | 452 | 99 |
| WP_041230848.1 | DR_0671/NP_294394.1 | 43 | 393 | 185 | 11 | 14 | 392 | 11 | 378 | 7.00E-76 | 240 | 96 |
| WP_041231552.1 | DR_1420/NP_295143.2 | 87.64 | 267 | 33 | 0 | 1 | 267 | 1 | 267 | 2.00E-162 | 452 | 100 |
| WP_083865818.1 | DR_1238/NP_294962.1 | 78.53 | 382 | 78 | 2 | 33 | 410 | 12 | 393 | 0 | 629 | 92 |

*Deinococcus phoenicis*

| Query ID | Subject ID | % Identity | Alignment length | Mismatches | Gap opens | Query start | Query end | Sequence start | Sequence end | E-value | Bit score | Query coverage |
|---|---|---|---|---|---|---|---|---|---|---|---|---|
| DR_1238/NP_294962.1\|homocitrate | WP_034356272.1 | 89.38 | 386 | 41 | 0 | 8 | 393 | 4 | 389 | 0 | 719 | 98 |
| DR_1610/NP_295333.1\|3-isopropylmalate | WP_034357910.1 | 90.02 | 421 | 41 | 1 | 8 | 427 | 4 | 424 | 0 | 725 | 97 |
| DR_1614/NP_295337.1\|3-isopropylmalate | WP_034357912.1 | 86.42 | 162 | 22 | 0 | 1 | 162 | 1 | 162 | 2.00E-101 | 290 | 92 |
| DR_1674/NP_295397.1\|isocitrate | WP_034355835.1 | 87.95 | 332 | 40 | 0 | 1 | 332 | 1 | 332 | 0 | 605 | 99 |
| DR_1588/NP_295311.1\|class | WP_034352500.1 | 75.25 | 400 | 92 | 3 | 22 | 419 | 12 | 406 | 0 | 566 | 95 |
| DR_2194/NP_295916.1\|ribosomal | WP_034357917.1 | 84.67 | 287 | 44 | 0 | 8 | 294 | 1 | 287 | 0 | 509 | 97 |
| DR_1420/NP_295143.2\|acetylglutamate/acetylaminoadipate | WP_034360177.1 | 88.01 | 267 | 32 | 0 | 1 | 267 | 1 | 267 | 4.00E-162 | 452 | 100 |
| DR_0963/NP_294687.2\|argC;N-acetyl-gamma-glutamyl-phosphate | WP_081790728.1 | 83.77 | 345 | 56 | 0 | 3 | 347 | 4 | 348 | 0 | 607 | 99 |
| DR_0794/NP_294518.1\|acetylornithine | WP_034359779.1 | 83.57 | 414 | 68 | 0 | 37 | 450 | 1 | 414 | 0 | 735 | 89 |
| DR_1413/NP_295136.1\|acetyl-lysine | WP_034356795.1 | 72.51 | 342 | 88 | 2 | 10 | 351 | 11 | 346 | 1.00E-167 | 474 | 94 |
| DR_1252/NP_294976.1\|hypothetical | WP_034356468.1 | 83.21 | 405 | 68 | 0 | 1 | 405 | 1 | 405 | 0 | 722 | 100 |
| DR_1365/NP_295088.1\|aspartate | WP_034357955.1 | 81.25 | 464 | 87 | 0 | 10 | 473 | 5 | 468 | 0 | 710 | 98 |
| DR_2008/NP_295731.1\|aspartate-semialdehyde | WP_034356169.1 | 80.54 | 334 | 65 | 0 | 1 | 334 | 1 | 334 | 0 | 544 | 99 |
| DR_1278/NP_295002.1\|homoserine | WP_034358473.1 | 76 | 325 | 73 | 1 | 1 | 325 | 1 | 320 | 7.00E-180 | 501 | 100 |
| DR_0671/NP_294394.1\|class | WP_034352558.1 | 65.61 | 378 | 119 | 2 | 3 | 380 | 6 | 372 | 3.00E-158 | 451 | 99 |
| DR_1758/NP_295481.1\|diaminopimelate | WP_034357559.1 | 70.71 | 379 | 111 | 0 | 1 | 379 | 1 | 379 | 8.00E-180 | 506 | 100 |
| DR_0297/NP_294020.1\|UDP-N-acetylmuramoylalanyl-D-glutamate--2_6-diaminopimelate | WP_034359073.1 | 81.04 | 480 | 88 | 2 | 1 | 480 | 1 | 477 | 0 | 706 | 98 |
| DR_0768/NP_294492.1\|UDP-N-acetylmuramoylalanyl-D-glutamyl-2_6-diaminopimelate--D-alanyl-D-alanyl | WP_034358741.1 | 75.35 | 434 | 95 | 4 | 1 | 432 | 1 | 424 | 0 | 604 | 99 |

**REVERSE BLAST**

| | | | | | | | | | | | | |
|---|---|---|---|---|---|---|---|---|---|---|---|---|
| WP_034352500.1 | DR_1588/NP_295311.1 | 73.35 | 409 | 102 | 3 | 3 | 406 | 13 | 419 | 0 | 591 | 99 |
| WP_034352558.1 | DR_0671/NP_294394.1 | 64.74 | 380 | 123 | 2 | 4 | 372 | 1 | 380 | 2.00E-160 | 456 | 98 |
| WP_034355835.1 | DR_1674/NP_295397.1 | 87.95 | 332 | 40 | 0 | 1 | 332 | 1 | 332 | 0 | 605 | 99 |
| WP_034356169.1 | DR_2008/NP_295731.1 | 80.54 | 334 | 65 | 0 | 1 | 334 | 1 | 334 | 0 | 544 | 99 |
| WP_034356272.1 | DR_1238/NP_294962.1 | 89.38 | 386 | 41 | 0 | 4 | 389 | 8 | 393 | 0 | 719 | 99 |
| WP_034356468.1 | DR_1252/NP_294976.1 | 83.21 | 405 | 68 | 0 | 1 | 405 | 1 | 405 | 0 | 722 | 100 |
| WP_034356795.1 | DR_1413/NP_295136.1 | 73.43 | 350 | 87 | 2 | 11 | 354 | 10 | 359 | 5.00E-179 | 503 | 94 |
| WP_034357559.1 | DR_1758/NP_295481.1 | 70.71 | 379 | 111 | 0 | 1 | 379 | 1 | 379 | 0 | 536 | 99 |
| WP_034357910.1 | DR_1610/NP_295333.1 | 90.02 | 421 | 41 | 1 | 4 | 424 | 8 | 427 | 0 | 727 | 98 |
| WP_034357912.1 | DR_1614/NP_295337.1 | 86.42 | 162 | 22 | 0 | 1 | 162 | 1 | 162 | 2.00E-101 | 290 | 93 |
| WP_034357917.1 | DR_2194/NP_295916.1 | 84.67 | 287 | 44 | 0 | 1 | 287 | 8 | 294 | 7.00E-177 | 491 | 99 |
| WP_034357955.1 | DR_1365/NP_295088.1 | 81.25 | 464 | 87 | 0 | 5 | 468 | 10 | 473 | 0 | 695 | 99 |
| WP_034358473.1 | DR_1278/NP_295002.1 | 76 | 325 | 73 | 1 | 1 | 320 | 1 | 325 | 6.00E-174 | 486 | 100 |
| WP_034358741.1 | DR_0768/NP_294492.1 | 75.35 | 434 | 95 | 4 | 1 | 424 | 1 | 432 | 0 | 617 | 99 |
| WP_034359073.1 | DR_0297/NP_294020.1 | 81.22 | 490 | 89 | 2 | 1 | 487 | 1 | 490 | 0 | 715 | 99 |
| WP_034359779.1 | DR_0794/NP_294518.1 | 83.57 | 414 | 68 | 0 | 1 | 414 | 37 | 450 | 0 | 735 | 99 |
| WP_034360177.1 | DR_1420/NP_295143.2 | 88.01 | 267 | 32 | 0 | 1 | 267 | 1 | 267 | 4.00E-162 | 452 | 100 |
| WP_081790728.1 | DR_0963/NP_294687.2 | 83.77 | 345 | 56 | 0 | 4 | 348 | 3 | 347 | 0 | 582 | 99 |

### *Deinococcus pimensis*

| Query ID | Subject ID | % Identity | Alignment length | Mismatches | Gap opens | Query start | Query end | Sequence start | Sequence end | E-value | Bit score | Query coverage |
|---|---|---|---|---|---|---|---|---|---|---|---|---|
| DR_1238/NP_294962.1\|homocitrate | WP_045233577.1 | 76.44 | 382 | 86 | 2 | 12 | 393 | 8 | 385 | 0 | 604 | 97 |
| DR_1610/NP_295333.1\|3-isopropylmalate | WP_045233590.1 | 84.35 | 409 | 63 | 1 | 21 | 429 | 1 | 408 | 0 | 676 | 94 |
| DR_1614/NP_295337.1\|3-isopropylmalate | WP_027480895.1 | 71.95 | 164 | 45 | 1 | 1 | 163 | 1 | 164 | 2.00E-85 | 249 | 92 |
| DR_1674/NP_295397.1\|isocitrate | WP_027481901.1 | 63.94 | 330 | 118 | 1 | 1 | 330 | 1 | 329 | 1.00E-152 | 433 | 99 |
| DR_1588/NP_295311.1\|class | WP_045233972.1 | 41.92 | 396 | 220 | 4 | 26 | 413 | 4 | 397 | 8.00E-98 | 299 | 92 |
| DR_2194/NP_295916.1\|ribosomal | WP_027480898.1 | 76.04 | 288 | 60 | 3 | 8 | 295 | 1 | 279 | 6.00E-158 | 443 | 98 |
| DR_1420/NP_295143.2\|acetylglutamate/acetylaminoadipate | WP_045233588.1 | 85.02 | 267 | 40 | 0 | 1 | 267 | 1 | 267 | 1.00E-154 | 434 | 100 |
| DR_0963/NP_294687.2\|argC;N-acetyl-gamma-glutamyl-phosphate | WP_084473930.1 | 79.29 | 338 | 70 | 0 | 10 | 347 | 1 | 338 | 0 | 554 | 97 |
| DR_0794/NP_294518.1\|acetylornithine | WP_027480884.1 | 75.3 | 413 | 100 | 2 | 37 | 448 | 1 | 412 | 0 | 629 | 89 |
| DR_1413/NP_295136.1\|acetyl-lysine | WP_027483089.1 | 25.49 | 357 | 226 | 14 | 27 | 356 | 29 | 372 | 4.00E-05 | 42 | 91 |
| DR_1252/NP_294976.1\|hypothetical | WP_027482233.1 | 79.95 | 404 | 81 | 0 | 1 | 404 | 1 | 404 | 0 | 698 | 99 |
| DR_1365/NP_295088.1\|aspartate | WP_027480760.1 | 31.3 | 115 | 69 | 3 | 202 | 309 | 117 | 228 | 1.00E-04 | 40.4 | 23 |
| DR_2008/NP_295731.1\|aspartate-semialdehyde | WP_027481960.1 | 74.54 | 271 | 69 | 0 | 1 | 271 | 1 | 271 | 9.00E-145 | 411 | 80 |
| DR_1278/NP_295002.1\|homoserine | WP_027481820.1 | 72.92 | 325 | 83 | 1 | 1 | 325 | 1 | 320 | 4.00E-166 | 467 | 100 |
| DR_0671/NP_294394.1\|class | WP_027481504.1 | 31.07 | 338 | 204 | 6 | 54 | 379 | 65 | 385 | 7.00E-42 | 150 | 86 |
| DR_1758/NP_295481.1\|diaminopimelate | WP_045233367.1 | 63.49 | 378 | 136 | 2 | 3 | 379 | 2 | 378 | 4.00E-156 | 446 | 99 |
| DR_0297/NP_294020.1\|UDP-N-acetylmuramoylalanyl-D-glutamate--2_6-diaminopimelate | WP_084475045.1 | 28.57 | 406 | 221 | 21 | 100 | 459 | 347 | 729 | 7.00E-14 | 70.9 | 73 |
| DR_0768/NP_294492.1\|UDP-N-acetylmuramoylalanyl-D-glutamyl-2_6-diaminopimelate--D-alanyl-D-alanyl | WP_084474532.1 | 63.19 | 144 | 48 | 1 | 286 | 429 | 1 | 139 | 1.00E-50 | 167 | 33 |

**REVERSE BLAST**

| | | | | | | | | | | | | |
|---|---|---|---|---|---|---|---|---|---|---|---|---|
| WP_027480760.1 | NP_295234.1 | 77.12 | 236 | 53 | 1 | 1 | 236 | 9 | 243 | 4.00E-131 | 371 | 99 |
| WP_027480884.1 | DR_0794/NP_294518.1 | 75.3 | 413 | 100 | 2 | 1 | 412 | 37 | 448 | 0 | 642 | 99 |
| WP_027480895.1 | DR_1614/NP_295337.1 | 71.95 | 164 | 45 | 1 | 1 | 164 | 1 | 163 | 1.00E-85 | 249 | 100 |
| WP_027480898.1 | DR_2194/NP_295916.1 | 76.04 | 288 | 60 | 3 | 1 | 279 | 8 | 295 | 8.00E-152 | 427 | 99 |
| WP_027481504.1 | NP_294346.1 | 71.01 | 376 | 109 | 0 | 6 | 381 | 7 | 382 | 0 | 560 | 97 |
| WP_027481820.1 | DR_1278/NP_295002.1 | 72.92 | 325 | 83 | 1 | 1 | 320 | 1 | 325 | 2.00E-166 | 467 | 100 |
| WP_027481901.1 | DR_1674/NP_295397.1 | 63.94 | 330 | 118 | 1 | 1 | 329 | 1 | 330 | 1.00E-152 | 433 | 99 |
| WP_027481960.1 | DR_2008/NP_295731.1 | 74.54 | 271 | 69 | 0 | 1 | 271 | 1 | 271 | 6.00E-145 | 411 | 100 |
| WP_027482233.1 | DR_1252/NP_294976.1 | 79.95 | 404 | 81 | 0 | 1 | 404 | 1 | 404 | 0 | 698 | 100 |
| WP_027483089.1 | NP_296213.1 | 31.78 | 321 | 187 | 14 | 65 | 372 | 66 | 367 | 6.00E-13 | 66.2 | 82 |
| WP_045233367.1 | DR_1758/NP_295481.1 | 63.49 | 378 | 136 | 2 | 2 | 378 | 3 | 379 | 5.00E-163 | 463 | 98 |
| WP_045233577.1 | DR_1238/NP_294962.1 | 76.44 | 382 | 86 | 2 | 8 | 385 | 12 | 393 | 0 | 604 | 98 |
| WP_045233588.1 | DR_1420/NP_295143.2 | 85.02 | 267 | 40 | 0 | 1 | 267 | 1 | 267 | 4.00E-154 | 432 | 94 |
| WP_045233590.1 | DR_1610/NP_295333.1 | 84.35 | 409 | 63 | 1 | 1 | 408 | 21 | 429 | 0 | 651 | 99 |
| WP_045233972.1 | DR_1588/NP_295311.1 | 42.35 | 392 | 216 | 4 | 4 | 393 | 26 | 409 | 2.00E-97 | 298 | 97 |
| WP_084473930.1 | DR_0963/NP_294687.2 | 79.29 | 338 | 70 | 0 | 1 | 338 | 10 | 347 | 0 | 531 | 100 |
| WP_084474532.1 | DR_0768/NP_294492.1 | 63.19 | 144 | 48 | 1 | 1 | 139 | 286 | 429 | 4.00E-51 | 167 | 96 |
| WP_084475045.1 | DR_0297/NP_294020.1 | 28.71 | 404 | 223 | 18 | 347 | 729 | 100 | 459 | 7.00E-13 | 68.2 | 50 |

### *Deinococcus puniceus*

| Query ID | Subject ID | % Identity | Alignment length | Mismatches | Gap opens | Query start | Query end | Sequence start | Sequence end | E-value | Bit score | Query coverage |
|---|---|---|---|---|---|---|---|---|---|---|---|---|
| DR_1238/NP_294962.1\|homocitrate | WP_064015010.1 | 89.66 | 387 | 39 | 1 | 8 | 393 | 11 | 397 | 0 | 720 | 98 |
| DR_1610/NP_295333.1\|3-isopropylmalate | WP_064014912.1 | 86.96 | 414 | 50 | 1 | 21 | 430 | 1 | 414 | 0 | 692 | 94 |
| DR_1614/NP_295337.1\|3-isopropylmalate | WP_064014911.1 | 82.84 | 169 | 29 | 0 | 1 | 169 | 1 | 169 | 3.00E-99 | 285 | 95 |
| DR_1674/NP_295397.1\|isocitrate | WP_064014584.1 | 88.25 | 332 | 39 | 0 | 1 | 332 | 1 | 332 | 0 | 607 | 99 |
| DR_1588/NP_295311.1\|class | WP_064013749.1 | 72 | 400 | 110 | 2 | 22 | 419 | 15 | 414 | 0 | 581 | 95 |
| DR_2194/NP_295916.1\|ribosomal | WP_064014908.1 | 84.38 | 288 | 40 | 1 | 8 | 295 | 1 | 283 | 4.00E-180 | 499 | 98 |
| DR_1420/NP_295143.2\|acetylglutamate/acetylaminoadipate | WP_064014707.1 | 90.64 | 267 | 25 | 0 | 1 | 267 | 1 | 267 | 1.00E-165 | 461 | 100 |
| DR_0963/NP_294687.2\|argC;N-acetyl-gamma-glutamyl-phosphate | WP_064015918.1 | 85.46 | 337 | 49 | 0 | 11 | 347 | 1 | 337 | 0 | 582 | 97 |
| DR_0794/NP_294518.1\|acetylornithine | WP_064015024.1 | 75.93 | 428 | 82 | 1 | 44 | 450 | 15 | 442 | 0 | 674 | 88 |
| DR_1413/NP_295136.1\|acetyl-lysine | WP_064014844.1 | 77.53 | 356 | 77 | 1 | 4 | 359 | 13 | 365 | 0 | 533 | 98 |
| DR_1252/NP_294976.1\|hypothetical | WP_064014853.1 | 81.98 | 405 | 73 | 0 | 1 | 405 | 1 | 405 | 0 | 716 | 100 |
| DR_1365/NP_295088.1\|aspartate | WP_064014671.1 | 81.52 | 460 | 85 | 0 | 10 | 469 | 7 | 466 | 0 | 685 | 97 |
| DR_2008/NP_295731.1\|aspartate-semialdehyde | WP_064016003.1 | 83.43 | 338 | 56 | 0 | 1 | 338 | 1 | 338 | 0 | 575 | 100 |
| DR_1278/NP_295002.1\|homoserine | WP_064014269.1 | 74.77 | 325 | 77 | 1 | 1 | 325 | 1 | 320 | 5.00E-163 | 458 | 100 |
| DR_0671/NP_294394.1\|class | WP_064013767.1 | 64.27 | 375 | 123 | 2 | 3 | 377 | 6 | 369 | 1.00E-158 | 452 | 98 |
| DR_1758/NP_295481.1\|diaminopimelate | WP_064014172.1 | 70.47 | 386 | 106 | 2 | 1 | 378 | 1 | 386 | 0 | 530 | 99 |
| DR_0297/NP_294020.1\|UDP-N-acetylmuramoylalanyl-D-glutamate--2_6-diaminopimelate | WP_064013559.1 | 78.16 | 490 | 106 | 1 | 1 | 490 | 1 | 489 | 0 | 681 | 100 |
| DR_0768/NP_294492.1\|UDP-N-acetylmuramoylalanyl-D-glutamyl-2_6-diaminopimelate--D-alanyl-D-alanyl | WP_064013989.1 | 69.79 | 437 | 116 | 4 | 1 | 431 | 1 | 427 | 0 | 576 | 99 |

**REVERSE BLAST**

| | | | | | | | | | | | | |
|---|---|---|---|---|---|---|---|---|---|---|---|---|
| WP_064013559.1 | DR_0297/NP_294020.1 | 78.16 | 490 | 106 | 1 | 1 | 489 | 1 | 490 | 0 | 696 | 99 |
| WP_064013749.1 | DR_1588/NP_295311.1 | 72 | 400 | 110 | 2 | 15 | 414 | 22 | 419 | 0 | 566 | 96 |
| WP_064013767.1 | DR_0671/NP_294394.1 | 63.4 | 377 | 127 | 2 | 4 | 369 | 1 | 377 | 3.00E-157 | 448 | 97 |
| WP_064013989.1 | DR_0768/NP_294492.1 | 69.79 | 437 | 116 | 4 | 1 | 427 | 1 | 431 | 0 | 576 | 100 |
| WP_064014172.1 | DR_1758/NP_295481.1 | 70.47 | 386 | 106 | 2 | 1 | 386 | 1 | 378 | 0 | 530 | 99 |
| WP_064014269.1 | DR_1278/NP_295002.1 | 74.77 | 325 | 77 | 1 | 1 | 320 | 1 | 325 | 9.00E-171 | 478 | 100 |
| WP_064014584.1 | DR_1674/NP_295397.1 | 88.25 | 332 | 39 | 0 | 1 | 332 | 1 | 332 | 0 | 607 | 99 |
| WP_064014671.1 | DR_1365/NP_295088.1 | 81.52 | 460 | 85 | 0 | 7 | 466 | 10 | 469 | 0 | 695 | 98 |
| WP_064014707.1 | DR_1420/NP_295143.2 | 90.64 | 267 | 25 | 0 | 1 | 267 | 1 | 267 | 6.00E-166 | 461 | 100 |
| WP_064014844.1 | DR_1413/NP_295136.1 | 77.16 | 359 | 73 | 2 | 13 | 365 | 4 | 359 | 0 | 553 | 94 |
| WP_064014853.1 | DR_1252/NP_294976.1 | 81.98 | 405 | 73 | 0 | 1 | 405 | 1 | 405 | 0 | 716 | 100 |
| WP_064014908.1 | DR_2194/NP_295916.1 | 84.38 | 288 | 40 | 1 | 1 | 283 | 8 | 295 | 9.00E-176 | 488 | 100 |
| WP_064014911.1 | DR_1614/NP_295337.1 | 82.84 | 169 | 29 | 0 | 1 | 169 | 1 | 169 | 4.00E-99 | 285 | 92 |
| WP_064014912.1 | DR_1610/NP_295333.1 | 86.96 | 414 | 50 | 1 | 1 | 414 | 21 | 430 | 0 | 692 | 98 |
| WP_064015010.1 | DR_1238/NP_294962.1 | 89.66 | 387 | 39 | 1 | 11 | 397 | 8 | 393 | 0 | 720 | 97 |
| WP_064015024.1 | DR_0794/NP_294518.1 | 75.93 | 428 | 82 | 1 | 15 | 442 | 44 | 450 | 0 | 674 | 96 |
| WP_064015918.1 | DR_0963/NP_294687.2 | 85.21 | 338 | 50 | 0 | 1 | 338 | 11 | 348 | 0 | 582 | 100 |
| WP_064016003.1 | DR_2008/NP_295731.1 | 83.43 | 338 | 56 | 0 | 1 | 338 | 1 | 338 | 0 | 575 | 99 |

### *Deinococcus reticulitermitis*

| Query ID | Subject ID | % Identity | Alignment length | Mismatches | Gap opens | Query start | Query end | Sequence start | Sequence end | E-value | Bit score | Query coverage |
|---|---|---|---|---|---|---|---|---|---|---|---|---|
| DR_1238/NP_294962.1\|homocitrate | WP_092262641.1 | 90.72 | 388 | 36 | 0 | 6 | 393 | 1 | 388 | 0 | 728 | 99 |
| DR_1610/NP_295333.1\|3-isopropylmalate | WP_092264977.1 | 88.02 | 434 | 47 | 2 | 1 | 434 | 1 | 429 | 0 | 728 | 100 |
| DR_1614/NP_295337.1\|3-isopropylmalate | WP_092262486.1 | 90.36 | 166 | 16 | 0 | 1 | 166 | 1 | 166 | 9.00E-109 | 308 | 94 |
| DR_1674/NP_295397.1\|isocitrate | WP_092263588.1 | 91.89 | 333 | 27 | 0 | 1 | 333 | 1 | 333 | 0 | 637 | 100 |
| DR_1588/NP_295311.1\|class | WP_092263643.1 | 80.63 | 413 | 74 | 2 | 8 | 420 | 6 | 412 | 0 | 662 | 98 |
| DR_2194/NP_295916.1\|ribosomal | WP_092264022.1 | 86.81 | 288 | 38 | 0 | 8 | 295 | 1 | 288 | 0 | 522 | 98 |
| DR_1420/NP_295143.2\|acetylglutamate/acetylaminoadipate | WP_092262618.1 | 92.13 | 267 | 21 | 0 | 1 | 267 | 1 | 267 | 4.00E-168 | 467 | 100 |
| DR_0963/NP_294687.2\|argC;N-acetyl-gamma-glutamyl-phosphate | WP_092265096.1 | 89.63 | 347 | 36 | 0 | 1 | 347 | 1 | 347 | 0 | 626 | 99 |
| DR_0794/NP_294518.1\|acetylornithine | WP_092264079.1 | 82.48 | 451 | 67 | 1 | 27 | 465 | 21 | 471 | 0 | 777 | 94 |
| DR_1413/NP_295136.1\|acetyl-lysine | WP_092262416.1 | 81.06 | 359 | 61 | 2 | 8 | 362 | 2 | 357 | 0 | 576 | 98 |
| DR_1252/NP_294976.1\|hypothetical | WP_092262579.1 | 90.86 | 405 | 37 | 0 | 1 | 405 | 1 | 405 | 0 | 777 | 100 |
| DR_1365/NP_295088.1\|aspartate | WP_092264943.1 | 87.23 | 470 | 59 | 1 | 5 | 473 | 3 | 472 | 0 | 766 | 99 |
| DR_2008/NP_295731.1\|aspartate-semialdehyde | WP_092265411.1 | 84.27 | 337 | 53 | 0 | 1 | 337 | 1 | 337 | 0 | 581 | 99 |
| DR_1278/NP_295002.1\|homoserine | WP_092263569.1 | 83.38 | 325 | 49 | 1 | 1 | 325 | 1 | 320 | 0 | 525 | 100 |
| DR_0671/NP_294394.1\|class | WP_092263446.1 | 69.31 | 378 | 109 | 2 | 3 | 380 | 6 | 376 | 9.00E-173 | 488 | 99 |
| DR_1758/NP_295481.1\|diaminopimelate | WP_092263782.1 | 76.78 | 379 | 88 | 0 | 1 | 379 | 1 | 379 | 0 | 546 | 100 |
| DR_0297/NP_294020.1\|UDP-N-acetylmuramoylalanyl-D-glutamate--2_6-diaminopimelate | WP_092265801.1 | 80.04 | 461 | 92 | 0 | 22 | 482 | 19 | 479 | 0 | 693 | 94 |
| DR_0768/NP_294492.1\|UDP-N-acetylmuramoylalanyl-D-glutamyl-2_6-diaminopimelate--D-alanyl-D-alanyl | WP_092264042.1 | 79.21 | 433 | 80 | 2 | 1 | 433 | 1 | 423 | 0 | 650 | 100 |

**REVERSE BLAST**

| | | | | | | | | | | | | |
|---|---|---|---|---|---|---|---|---|---|---|---|---|
| WP_092262416.1 | DR_1413/NP_295136.1 | 81.06 | 359 | 61 | 2 | 2 | 357 | 8 | 362 | 0 | 576 | 99 |
| WP_092262486.1 | DR_1614/NP_295337.1 | 90.36 | 166 | 16 | 0 | 1 | 166 | 1 | 166 | 8.00E-109 | 308 | 100 |
| WP_092262579.1 | DR_1252/NP_294976.1 | 90.86 | 405 | 37 | 0 | 1 | 405 | 1 | 405 | 0 | 777 | 100 |
| WP_092262618.1 | DR_1420/NP_295143.2 | 92.13 | 267 | 21 | 0 | 1 | 267 | 1 | 267 | 3.00E-168 | 467 | 100 |
| WP_092262641.1 | DR_1238/NP_294962.1 | 90.72 | 388 | 36 | 0 | 1 | 388 | 6 | 393 | 0 | 728 | 100 |
| WP_092263446.1 | DR_0671/NP_294394.1 | 68.68 | 380 | 112 | 2 | 4 | 376 | 1 | 380 | 1.00E-173 | 490 | 98 |
| WP_092263569.1 | DR_1278/NP_295002.1 | 83.38 | 325 | 49 | 1 | 1 | 320 | 1 | 325 | 0 | 529 | 100 |
| WP_092263588.1 | DR_1674/NP_295397.1 | 91.89 | 333 | 27 | 0 | 1 | 333 | 1 | 333 | 0 | 637 | 100 |
| WP_092263643.1 | DR_1588/NP_295311.1 | 80.63 | 413 | 74 | 2 | 6 | 412 | 8 | 420 | 0 | 655 | 99 |
| WP_092263782.1 | DR_1758/NP_295481.1 | 76.78 | 379 | 88 | 0 | 1 | 379 | 1 | 379 | 0 | 583 | 98 |
| WP_092264022.1 | DR_2194/NP_295916.1 | 86.81 | 288 | 38 | 0 | 1 | 288 | 8 | 295 | 0 | 507 | 100 |
| WP_092264042.1 | DR_0768/NP_294492.1 | 79.21 | 433 | 80 | 2 | 1 | 423 | 1 | 433 | 0 | 672 | 99 |
| WP_092264079.1 | DR_0794/NP_294518.1 | 82.48 | 451 | 67 | 1 | 21 | 471 | 27 | 465 | 0 | 777 | 96 |
| WP_092264943.1 | DR_1365/NP_295088.1 | 87.23 | 470 | 59 | 1 | 3 | 472 | 5 | 473 | 0 | 750 | 99 |
| WP_092264977.1 | DR_1610/NP_295333.1 | 88.02 | 434 | 47 | 2 | 1 | 429 | 1 | 434 | 0 | 721 | 100 |
| WP_092265096.1 | DR_0963/NP_294687.2 | 89.63 | 347 | 36 | 0 | 1 | 347 | 1 | 347 | 0 | 627 | 99 |
| WP_092265411.1 | DR_2008/NP_295731.1 | 84.27 | 337 | 53 | 0 | 1 | 337 | 1 | 337 | 0 | 581 | 99 |
| WP_092265801.1 | DR_0297/NP_294020.1 | 79.1 | 488 | 99 | 1 | 1 | 485 | 1 | 488 | 0 | 701 | 98 |

### *Deinococcus soli*

| Query ID | Subject ID | % Identity | Alignment length | Mismatches | Gap opens | Query start | Query end | Sequence start | Sequence end | E-value | Bit score | Query coverage |
|---|---|---|---|---|---|---|---|---|---|---|---|---|
| DR_1238/NP_294962.1\|homocitrate | WP_046842651.1 | 91.47 | 387 | 33 | 0 | 7 | 393 | 9 | 395 | 0 | 739 | 98 |
| DR_1610/NP_295333.1\|3-isopropylmalate | WP_046842606.1 | 87.74 | 424 | 48 | 1 | 15 | 434 | 5 | 428 | 0 | 713 | 97 |
| DR_1614/NP_295337.1\|3-isopropylmalate | WP_046842605.1 | 93.21 | 162 | 11 | 0 | 1 | 162 | 1 | 162 | 3.00E-110 | 312 | 92 |
| DR_1674/NP_295397.1\|isocitrate | WP_046842937.1 | 92.49 | 333 | 25 | 0 | 1 | 333 | 1 | 333 | 0 | 637 | 100 |
| DR_1588/NP_295311.1\|class | WP_046844009.1 | 76.11 | 406 | 91 | 2 | 16 | 420 | 6 | 406 | 0 | 622 | 96 |
| DR_2194/NP_295916.1\|ribosomal | WP_046842602.1 | 89.2 | 287 | 31 | 0 | 8 | 294 | 1 | 287 | 0 | 506 | 97 |
| DR_1420/NP_295143.2\|acetylglutamate/acetylaminoadipate | WP_046844754.1 | 88.76 | 267 | 30 | 0 | 1 | 267 | 1 | 267 | 2.00E-163 | 455 | 100 |
| DR_0963/NP_294687.2\|argC;N-acetyl-gamma-glutamyl-phosphate | WP_081424482.1 | 86.63 | 344 | 46 | 0 | 4 | 347 | 5 | 348 | 0 | 594 | 99 |
| DR_0794/NP_294518.1\|acetylornithine | WP_046842479.1 | 81.51 | 411 | 76 | 0 | 45 | 455 | 6 | 416 | 0 | 692 | 88 |
| DR_1413/NP_295136.1\|acetyl-lysine | WP_046843345.1 | 74.16 | 356 | 87 | 2 | 8 | 362 | 9 | 360 | 7.00E-178 | 499 | 98 |
| DR_1252/NP_294976.1\|hypothetical | WP_046844235.1 | 89.88 | 405 | 41 | 0 | 1 | 405 | 1 | 405 | 0 | 771 | 100 |
| DR_1365/NP_295088.1\|aspartate | WP_046842439.1 | 84.06 | 458 | 73 | 0 | 10 | 467 | 5 | 462 | 0 | 702 | 98 |
| DR_1365/NP_295088.1\|aspartate | WP_046842439.1 | 35.71 | 70 | 45 | 0 | 402 | 471 | 322 | 391 | 1.1 | 28.1 | 98 |
| DR_2008/NP_295731.1\|aspartate-semialdehyde | WP_046844284.1 | 83.09 | 337 | 57 | 0 | 1 | 337 | 1 | 337 | 0 | 576 | 99 |
| DR_1278/NP_295002.1\|homoserine | WP_046844389.1 | 77.85 | 325 | 67 | 1 | 1 | 325 | 1 | 320 | 8.00E-177 | 493 | 100 |
| DR_0671/NP_294394.1\|class | WP_046843851.1 | 64.63 | 376 | 120 | 4 | 1 | 376 | 4 | 366 | 1.00E-141 | 408 | 99 |
| DR_1758/NP_295481.1\|diaminopimelate | WP_046844438.1 | 69.05 | 378 | 117 | 0 | 1 | 378 | 1 | 378 | 2.00E-170 | 482 | 99 |
| DR_0297/NP_294020.1\|UDP-N-acetylmuramoylalanyl-D-glutamate--2_6-diaminopimelate | WP_046843507.1 | 78.47 | 483 | 103 | 1 | 1 | 483 | 1 | 482 | 0 | 691 | 99 |
| DR_0768/NP_294492.1\|UDP-N-acetylmuramoylalanyl-D-glutamyl-2_6-diaminopimelate--D-alanyl-D-alanyl | WP_046843104.1 | 74.89 | 438 | 94 | 4 | 1 | 432 | 1 | 428 | 0 | 629 | 99 |

**REVERSE BLAST**

| | | | | | | | | | | | | |
|---|---|---|---|---|---|---|---|---|---|---|---|---|
| WP_046842439.1 | DR_1365/NP_295088.1 | 84.06 | 458 | 73 | 0 | 5 | 462 | 10 | 467 | 0 | 711 | 98 |
| WP_046842439.1 | DR_1365/NP_295088.1 | 34.21 | 76 | 50 | 0 | 316 | 391 | 396 | 471 | 3.00E-06 | 45.8 | 98 |
| WP_046842479.1 | DR_0794/NP_294518.1 | 81.51 | 411 | 76 | 0 | 6 | 416 | 45 | 455 | 0 | 712 | 99 |
| WP_046842602.1 | DR_2194/NP_295916.1 | 89.2 | 287 | 31 | 0 | 1 | 287 | 8 | 294 | 0 | 506 | 100 |
| WP_046842605.1 | DR_1614/NP_295337.1 | 93.21 | 162 | 11 | 0 | 1 | 162 | 1 | 162 | 3.00E-110 | 312 | 99 |
| WP_046842606.1 | DR_1610/NP_295333.1 | 87.74 | 424 | 48 | 1 | 5 | 428 | 15 | 434 | 0 | 712 | 99 |
| WP_046842651.1 | DR_1238/NP_294962.1 | 91.47 | 387 | 33 | 0 | 9 | 395 | 7 | 393 | 0 | 739 | 98 |
| WP_046842937.1 | DR_1674/NP_295397.1 | 92.49 | 333 | 25 | 0 | 1 | 333 | 1 | 333 | 0 | 637 | 100 |
| WP_046843104.1 | DR_0768/NP_294492.1 | 74.89 | 438 | 94 | 4 | 1 | 428 | 1 | 432 | 0 | 629 | 99 |
| WP_046843345.1 | DR_1413/NP_295136.1 | 74.16 | 356 | 87 | 2 | 9 | 360 | 8 | 362 | 0 | 512 | 96 |
| WP_046843507.1 | DR_0297/NP_294020.1 | 78.47 | 483 | 103 | 1 | 1 | 482 | 1 | 483 | 0 | 689 | 99 |
| WP_046843851.1 | DR_0671/NP_294394.1 | 64.36 | 376 | 121 | 4 | 4 | 366 | 1 | 376 | 1.00E-150 | 431 | 97 |
| WP_046844009.1 | DR_1588/NP_295311.1 | 75.37 | 406 | 94 | 2 | 6 | 406 | 16 | 420 | 0 | 615 | 99 |
| WP_046844235.1 | DR_1252/NP_294976.1 | 89.88 | 405 | 41 | 0 | 1 | 405 | 1 | 405 | 0 | 771 | 100 |
| WP_046844284.1 | DR_2008/NP_295731.1 | 83.09 | 337 | 57 | 0 | 1 | 337 | 1 | 337 | 0 | 576 | 99 |
| WP_046844389.1 | DR_1278/NP_295002.1 | 77.85 | 325 | 67 | 1 | 1 | 320 | 1 | 325 | 1.00E-176 | 493 | 100 |
| WP_046844438.1 | DR_1758/NP_295481.1 | 69.05 | 378 | 117 | 0 | 1 | 378 | 1 | 378 | 0 | 519 | 99 |
| WP_046844754.1 | DR_1420/NP_295143.2 | 88.76 | 267 | 30 | 0 | 1 | 267 | 1 | 267 | 3.00E-163 | 454 | 100 |
| WP_081424482.1 | DR_0963/NP_294687.2 | 86.63 | 344 | 46 | 0 | 5 | 348 | 4 | 347 | 0 | 594 | 99 |

### *Deinococcus swuensis*

| Query ID | Subject ID | % Identity | Alignment length | Mismatches | Gap opens | Query start | Query end | Sequence start | Sequence end | E-value | Bit score | Query coverage |
|---|---|---|---|---|---|---|---|---|---|---|---|---|
| DR_1238/NP_294962.1\|homocitrate | WP_052195446.1 | 90.31 | 382 | 37 | 0 | 12 | 393 | 14 | 395 | 0 | 719 | 97 |
| DR_1610/NP_295333.1\|3-isopropylmalate | WP_081995052.1 | 88.45 | 407 | 45 | 1 | 21 | 427 | 1 | 405 | 0 | 694 | 94 |
| DR_1614/NP_295337.1\|3-isopropylmalate | WP_039683632.1 | 85.38 | 171 | 25 | 0 | 1 | 171 | 1 | 171 | 4.00E-105 | 300 | 97 |
| DR_1674/NP_295397.1\|isocitrate | WP_039683063.1 | 89.76 | 332 | 34 | 0 | 1 | 332 | 1 | 332 | 0 | 614 | 99 |
| DR_1588/NP_295311.1\|class | WP_039687402.1 | 75.5 | 400 | 92 | 2 | 21 | 419 | 8 | 402 | 0 | 610 | 95 |
| DR_2194/NP_295916.1\|ribosomal | WP_039683638.1 | 83.97 | 287 | 46 | 0 | 8 | 294 | 1 | 287 | 0 | 504 | 97 |
| DR_1420/NP_295143.2\|acetylglutamate/acetylaminoadipate | WP_039682409.1 | 90.64 | 267 | 25 | 0 | 1 | 267 | 1 | 267 | 3.00E-166 | 462 | 100 |
| DR_0963/NP_294687.2\|argC;N-acetyl-gamma-glutamyl-phosphate | WP_081995135.1 | 86.99 | 346 | 45 | 0 | 2 | 347 | 5 | 350 | 0 | 629 | 99 |
| DR_0794/NP_294518.1\|acetylornithine | WP_039684598.1 | 83.78 | 413 | 67 | 0 | 38 | 450 | 4 | 416 | 0 | 731 | 89 |
| DR_1413/NP_295136.1\|acetyl-lysine | WP_039685416.1 | 72.02 | 361 | 94 | 3 | 6 | 359 | 8 | 368 | 2.00E-177 | 499 | 98 |
| DR_1252/NP_294976.1\|hypothetical | WP_039682321.1 | 86.63 | 404 | 54 | 0 | 1 | 404 | 1 | 404 | 0 | 743 | 99 |
| DR_1365/NP_295088.1\|aspartate | WP_039683644.1 | 83.33 | 462 | 77 | 0 | 10 | 471 | 5 | 466 | 0 | 716 | 98 |
| DR_1365/NP_295088.1\|aspartate | WP_039683644.1 | 36.99 | 73 | 46 | 0 | 326 | 398 | 396 | 468 | 2.00E-06 | 46.6 | 98 |
| DR_2008/NP_295731.1\|aspartate-semialdehyde | WP_039681543.1 | 84.73 | 334 | 51 | 0 | 1 | 334 | 1 | 334 | 0 | 582 | 99 |
| DR_1278/NP_295002.1\|homoserine | WP_039686059.1 | 76.92 | 325 | 70 | 1 | 1 | 325 | 1 | 320 | 0 | 506 | 100 |
| DR_0671/NP_294394.1\|class | WP_039686140.1 | 69.52 | 374 | 112 | 2 | 3 | 376 | 6 | 377 | 5.00E-169 | 479 | 98 |
| DR_1758/NP_295481.1\|diaminopimelate | WP_039684073.1 | 70.05 | 384 | 110 | 1 | 1 | 379 | 1 | 384 | 2.00E-179 | 505 | 100 |
| DR_0297/NP_294020.1\|UDP-N-acetylmuramoylalanyl-D-glutamate--2_6-diaminopimelate | WP_039686130.1 | 78.97 | 466 | 97 | 1 | 15 | 480 | 13 | 477 | 0 | 673 | 95 |
| DR_0768/NP_294492.1\|UDP-N-acetylmuramoylalanyl-D-glutamyl-2_6-diaminopimelate--D-alanyl-D-alanyl | WP_039682824.1 | 75.06 | 433 | 97 | 3 | 1 | 432 | 1 | 423 | 0 | 592 | 99 |

**REVERSE BLAST**

| | | | | | | | | | | | | |
|---|---|---|---|---|---|---|---|---|---|---|---|---|
| WP_039681543.1 | DR_2008/NP_295731.1 | 84.73 | 334 | 51 | 0 | 1 | 334 | 1 | 334 | 0 | 582 | 99 |
| WP_039682321.1 | DR_1252/NP_294976.1 | 86.63 | 404 | 54 | 0 | 1 | 404 | 1 | 404 | 0 | 743 | 100 |
| WP_039682409.1 | DR_1420/NP_295143.2 | 90.64 | 267 | 25 | 0 | 1 | 267 | 1 | 267 | 1.00E-166 | 463 | 100 |
| WP_039682824.1 | DR_0768/NP_294492.1 | 75.06 | 433 | 97 | 3 | 1 | 423 | 1 | 432 | 0 | 613 | 99 |
| WP_039683063.1 | DR_1674/NP_295397.1 | 89.76 | 332 | 34 | 0 | 1 | 332 | 1 | 332 | 0 | 614 | 100 |
| WP_039683632.1 | DR_1614/NP_295337.1 | 85.38 | 171 | 25 | 0 | 1 | 171 | 1 | 171 | 3.00E-105 | 300 | 98 |
| WP_039683638.1 | DR_2194/NP_295916.1 | 83.97 | 287 | 46 | 0 | 1 | 287 | 8 | 294 | 4.00E-175 | 487 | 97 |
| WP_039683644.1 | DR_1365/NP_295088.1 | 83.33 | 462 | 77 | 0 | 5 | 466 | 10 | 471 | 0 | 702 | 98 |
| WP_039684073.1 | DR_1758/NP_295481.1 | 70.05 | 384 | 110 | 1 | 1 | 384 | 1 | 379 | 0 | 529 | 99 |
| WP_039684598.1 | DR_0794/NP_294518.1 | 83.78 | 413 | 67 | 0 | 4 | 416 | 38 | 450 | 0 | 731 | 98 |
| WP_039685416.1 | DR_1413/NP_295136.1 | 72.02 | 361 | 94 | 3 | 8 | 368 | 6 | 359 | 2.00E-177 | 499 | 96 |
| WP_039686059.1 | DR_1278/NP_295002.1 | 76.92 | 325 | 70 | 1 | 1 | 320 | 1 | 325 | 5.00E-176 | 491 | 100 |
| WP_039686130.1 | DR_0297/NP_294020.1 | 78.48 | 488 | 102 | 2 | 1 | 485 | 1 | 488 | 0 | 685 | 98 |
| WP_039686140.1 | DR_0671/NP_294394.1 | 68.88 | 376 | 115 | 2 | 4 | 377 | 1 | 376 | 5.00E-172 | 486 | 97 |
| WP_039687402.1 | DR_1588/NP_295311.1 | 73.58 | 405 | 101 | 2 | 3 | 402 | 16 | 419 | 0 | 600 | 99 |
| WP_052195446.1 | DR_1238/NP_294962.1 | 90.31 | 382 | 37 | 0 | 14 | 395 | 12 | 393 | 0 | 719 | 97 |
| WP_081995052.1 | DR_1610/NP_295333.1 | 88.45 | 407 | 45 | 1 | 1 | 405 | 21 | 427 | 0 | 694 | 99 |
| WP_081995135.1 | DR_0963/NP_294687.2 | 86.99 | 346 | 45 | 0 | 5 | 350 | 2 | 347 | 0 | 604 | 99 |

### *Deinococcus wulumuqiensis*

| Query ID | Subject ID | % Identity | Alignment length | Mismatches | Gap opens | Query start | Query end | Sequence start | Sequence end | E-value | Bit score | Query coverage |
|---|---|---|---|---|---|---|---|---|---|---|---|---|
| DR_1238/NP_294962.1\|homocitrate | WP_025567708.1 | 96.39 | 388 | 14 | 0 | 6 | 393 | 1 | 388 | 0 | 772 | 99 |
| DR_1610/NP_295333.1\|3-isopropylmalate | WP_026138683.1 | 91.06 | 436 | 37 | 1 | 1 | 434 | 1 | 436 | 0 | 759 | 100 |
| DR_1614/NP_295337.1\|3-isopropylmalate | WP_017869523.1 | 94.48 | 163 | 9 | 0 | 1 | 163 | 1 | 163 | 1.00E-112 | 318 | 92 |
| DR_1674/NP_295397.1\|isocitrate | WP_017869576.1 | 94.59 | 333 | 18 | 0 | 1 | 333 | 1 | 333 | 0 | 649 | 100 |
| DR_1588/NP_295311.1\|class | WP_017869364.1 | 33.96 | 374 | 218 | 9 | 51 | 412 | 13 | 369 | 6.00E-47 | 164 | 86 |
| DR_2194/NP_295916.1\|ribosomal | WP_017869708.1 | 35.82 | 67 | 41 | 2 | 82 | 146 | 99 | 165 | 0.002 | 35.8 | 22 |
| DR_1420/NP_295143.2\|acetylglutamate/acetylaminoadipate | WP_017870427.1 | 94.76 | 267 | 14 | 0 | 1 | 267 | 1 | 267 | 9.00E-175 | 484 | 100 |
| DR_0963/NP_294687.2\|argC;N-acetyl-gamma-glutamyl-phosphate | WP_017868966.1 | 93.08 | 347 | 24 | 0 | 1 | 347 | 1 | 347 | 0 | 647 | 99 |
| DR_0794/NP_294518.1\|acetylornithine | WP_017870540.1 | 88.18 | 423 | 50 | 0 | 43 | 465 | 3 | 425 | 0 | 785 | 91 |
| DR_1413/NP_295136.1\|acetyl-lysine | WP_017870420.1 | 90.28 | 360 | 31 | 1 | 1 | 360 | 1 | 356 | 0 | 620 | 99 |
| DR_1252/NP_294976.1\|hypothetical | WP_017869895.1 | 94.57 | 405 | 22 | 0 | 1 | 405 | 1 | 405 | 0 | 805 | 100 |
| DR_1365/NP_295088.1\|aspartate | WP_040383940.1 | 94.27 | 471 | 27 | 0 | 1 | 471 | 1 | 471 | 0 | 836 | 99 |
| DR_2008/NP_295731.1\|aspartate-semialdehyde | WP_017871069.1 | 94.29 | 333 | 19 | 0 | 1 | 333 | 1 | 333 | 0 | 640 | 99 |
| DR_1278/NP_295002.1\|homoserine | WP_025568029.1 | 91.08 | 325 | 24 | 1 | 1 | 325 | 1 | 320 | 0 | 578 | 100 |
| DR_0671/NP_294394.1\|class | WP_017870104.1 | 32.34 | 337 | 199 | 8 | 54 | 378 | 66 | 385 | 5.00E-36 | 133 | 85 |
| DR_1758/NP_295481.1\|diaminopimelate | WP_025567190.1 | 85.45 | 378 | 55 | 0 | 1 | 378 | 1 | 378 | 0 | 654 | 99 |
| DR_0297/NP_294020.1\|UDP-N-acetylmuramoylalanyl-D-glutamate--2_6-diaminopimelate | WP_017870576.1 | 87.52 | 505 | 46 | 2 | 1 | 490 | 1 | 503 | 0 | 804 | 100 |
| DR_0768/NP_294492.1\|UDP-N-acetylmuramoylalanyl-D-glutamyl-2_6-diaminopimelate--D-alanyl-D-alanyl | WP_017870516.1 | 98.15 | 433 | 8 | 0 | 1 | 433 | 1 | 433 | 0 | 842 | 100 |

**REVERSE BLAST**

| | | | | | | | | | | | | |
|---|---|---|---|---|---|---|---|---|---|---|---|---|
| WP_017868966.1 | DR_0963/NP_294687.2 | 93.08 | 347 | 24 | 0 | 1 | 347 | 1 | 347 | 0 | 648 | 99 |
| WP_017869364.1 | NP_294754.1 | 84.49 | 374 | 56 | 1 | 1 | 372 | 1 | 374 | 0 | 620 | 99 |
| WP_017869523.1 | DR_1614/NP_295337.1 | 94.48 | 163 | 9 | 0 | 1 | 163 | 1 | 163 | 1.00E-112 | 318 | 97 |
| WP_017869576.1 | DR_1674/NP_295397.1 | 94.59 | 333 | 18 | 0 | 1 | 333 | 1 | 333 | 0 | 649 | 100 |
| WP_017869708.1 | NP_293843.1 | 97.08 | 445 | 13 | 0 | 1 | 445 | 1 | 445 | 0 | 897 | 100 |
| WP_017869895.1 | DR_1252/NP_294976.1 | 94.57 | 405 | 22 | 0 | 1 | 405 | 1 | 405 | 0 | 805 | 100 |
| WP_017870104.1 | NP_294346.1 | 88.14 | 388 | 46 | 0 | 1 | 388 | 1 | 388 | 0 | 693 | 100 |
| WP_017870420.1 | DR_1413/NP_295136.1 | 91.11 | 360 | 28 | 1 | 1 | 356 | 1 | 360 | 0 | 655 | 99 |
| WP_017870427.1 | DR_1420/NP_295143.2 | 94.76 | 267 | 14 | 0 | 1 | 267 | 1 | 267 | 3.00E-171 | 475 | 100 |
| WP_017870516.1 | DR_0768/NP_294492.1 | 98.15 | 433 | 8 | 0 | 1 | 433 | 1 | 433 | 0 | 842 | 100 |
| WP_017870540.1 | DR_0794/NP_294518.1 | 88.18 | 423 | 50 | 0 | 3 | 425 | 43 | 465 | 0 | 785 | 99 |
| WP_017870576.1 | DR_0297/NP_294020.1 | 87.52 | 505 | 46 | 2 | 1 | 503 | 1 | 490 | 0 | 804 | 99 |
| WP_017871069.1 | DR_2008/NP_295731.1 | 94.29 | 333 | 19 | 0 | 1 | 333 | 1 | 333 | 0 | 640 | 99 |
| WP_025567190.1 | DR_1758/NP_295481.1 | 85.45 | 378 | 55 | 0 | 1 | 378 | 1 | 378 | 0 | 654 | 99 |
| WP_025567708.1 | DR_1238/NP_294962.1 | 96.39 | 388 | 14 | 0 | 1 | 388 | 6 | 393 | 0 | 772 | 100 |
| WP_025568029.1 | DR_1278/NP_295002.1 | 91.08 | 325 | 24 | 1 | 1 | 320 | 1 | 325 | 0 | 578 | 100 |
| WP_026138683.1 | DR_1610/NP_295333.1 | 91.06 | 436 | 37 | 1 | 1 | 436 | 1 | 434 | 0 | 759 | 100 |
| WP_040383940.1 | DR_1365/NP_295088.1 | 94.27 | 471 | 27 | 0 | 1 | 471 | 1 | 471 | 0 | 815 | 99 |

#### ROBETTA ALIGNMENT DATA

| No. | NAME | LENGTH | TEMPLATES | ORGANISM | CONFIDENCE SCORE | ALIGNMENT |
| --- | --- | --- | --- | --- | --- | --- |
| 1. | (NP_294394.1) Class I Aminotransferase | 381 | 1bkg, 1o4s, 1u08, 1v2d, 1w7l, 1yiy, 2o0r, 3fvs, 3fvx, 4wp0, 5veh, 6f35, 6f77, 6l1l, 6l1n | *Deinococcus radiodurans* | 0.89 | MHPRARHAQTSIFTHMSLLAARHGAVNLGQGFPSNPPPAFLLDAARRAVGTVDQYTPPLGLPALREALGADLNVDPADVIVTSGATEGLLTLALSL<br>LVPTRDGQPAELVVFEPVYDVYVPQAELAGARAVPVPLHLDPEEGWSLDLAAVRAAITPRTQALLVNTPHNPTGLVFSWAELTELVALAREHDLW<br>LISDEVYDELYASERPTSLRELAPERTFTVGSAGKRLEVTGWRVGWILCPPGLGGGLSLAGGLANVRQVTSFCSPAPLQAAVAEALPLARSQGYYAA<br>LRADYAARRALLSSGLRSLGAQVHEPQGTYFLTAQHPTWTAEGLVESGAVAVIPGEAFYVTSPPPAGLLRFAFCKSRAELEQALERLARLSNF<br>>6f35A_201 weight: 0.3363 score: 860.28 eval: 1e-45 prob: 100 identity: 0.2677 startpos: 4<br>PASRISSIGVSEILKIGARAAakPVIILGAGEPDFDTPEHVKQAASDAig-ETKYTALDGTPELKKAIREKfaYELDEITVATGAKQILFNAMMASLDPG---<br>--DEVIIPTPYWTSYSDIVHICEGKPVLIACDAS—<br>SGFRLTAEKLEAAITPRTRWVLLNSPSNPSGAAYSAADYRPLLEVLLRhhVWLLVDDMYEHIVYDgrFVTPAQlIKNRTLTVNGVSKAYAMTGWRIG<br>YA<br>GGPRELI------KAMAVVQSQATSCPSSISQAASVVALNGP—<br>QDFLKERTESFQRRRDLVVNGLNAigLDCRVPEGAFYTFSGCAGvfCAYLLEDAHVAV<br>VPGSAFGL-----SPFFRISYATSEAELKEALERIAAACDR<br>>1v2dA_101 weight: 0.3026 score: 379.8 eval: 1.1e-11 prob: n/a identity: 0.5853 startpos: 11<br>----------IFPRMSGLAQRLGAVNLGQGFPSNPPPPFLLEAVRRALGRQDQYAPPAGLPALREALAEEFAVEPESVVVTSGATEALYVLLQSLVGP----<br>-GDEVVVLEPFFDVYLPDAFLAGAKARLVRLDLTP-<br>EGFRLDLSALEKALTPRTRALLLNTPMNPTGLVFGERELEAIARLARAHDLFLISDEVYDELYYGERPRRLREFAPERTFTVGSAGKRLEATGYRVGWI<br>VG<br>PK------<br>EFMPRLAGMRQWTSFSAPTPLQAGVAEALKLARREGFYEALREGYRRRRDLLAGGLRAMGLRVYVPEGTYFLMAELPGWDAFRLVEEARVA<br>LIPASAFYLED-PPKDLFRFAFCKTEEELHLALERLGRVV--<br>>1v2dA_301 weight: 0.2957 score: 18.78 eval: n/a prob: n/a identity: 0.5853 startpos: 1<br>-<br>MRLHPRTEAAIFPRMSGLAQRLGAVNLGQGFPSNPPPPFLLEAVRRALGRQDQYAPPAGLPALREALAEEFAVEPESVVVTSGATEALYVLLQSLV<br>GP<br>G-----DEVVVLEPFFDVYLPDAFLAGAKARLVRLDLTP-<br>EGFRLDLSALEKALTPRTRALLLNTPMNPTGLVFGERELEAIARLARAHDLFLISDEVYDELYYGERPRRLREFAPERTFTVGSAGKRLEATGYRVGWI<br>VG<br>PKEFM------<br>PRLAGMRQWTSFSAPTPLQAGVAEALKLARREGFYEALREGYRRRRDLLAGGLRAMGLRVYVPEGTYFLMAELPGWDAFRLVEEARVA<br>LIPASAFY-LEDPPKDLFRFAFCKTEEELHLALERLGRVV--<br>>6f35A_102 weight: 0.0087 score: 365.5 eval: 3.1e-11 prob: n/a identity: 0.2598 startpos: 4<br>PASRISSIGVSEILKIGARAAAmpVIILGAGEPDFDTPEHVKQAASDAIHreTKYTALDGTPELKKAIREKfaYELDEITVATGAKQILFNAMMASLDP---<br>--GDEVIIPTPYWTSYSDIVHICEGKPVLIACDAS—<br>SGFRLTAEKLEAAITPRTRWVLLNSPSNPSGAAYSAADYRPLLEVLLRhhVWLLVDDMYEHIVYDGffVTPAQLekNRTLTVNGVSKAYAMTGWRI<br>GYA<br>GGPR------ELIKAMAVVQSQATSCPSSISQAASVVALNGPQ—<br>DFLKERTESFQRRRDLVVNGLNAigLDCRVPEGAFYTFSGCAGvfCAYLLEDAHVAVV<br>PGSAFGL-----SPFFRISYATSEAELKEALERIAAACDR<br>>2o0rA_302 weight: 0.0074 score: 18.7 eval: n/a prob: n/a identity: 0.4016 startpos: 2<br>TVSRLRPYATTVFAEMSALATRIGAVNLGQGFPDEDGPPKMLQAAQDAIAgvNQYPPGPGSAPLRRAIAAQRryDptEVLVTVGATEAIAAAVLGL<br>VEP<br>G-----SEVLLIEPFYDSYSPVVAMAGAHRVTVPLV-<br>PDGRGFALDADALRRAVTPRTRALIINSPHNPTGAVLSATELAAIAEIAVAANLVVITDEVYEHLVFDHahLPLAGfmAERTITISSAAKMFNCTGWKI<br>GW<br>ACGPAELIAGVRAAKQYLS------YVGGAPFQPAVALALDTE--<br>DAWVAALRNSLRARRDRLAAGLTEIGFAVHDSYGTYFLCADPRPLgcAALPEKVGVAAIPMSAFCDPADVWNHLVRFTFCKRDDTLDEAIRRLSVL<br>AE- |

| | | | | | | |
|---|---|---|---|---|---|---|
|  |  |  |  |  |  | >5vehA_202 weight: 0.0063 score: 827.2 eval: 1e-45 prob: 100 identity: 0.2966 startpos: 4<br>LPKRYQGSTKSVWVEYIQLAAQYKPLNLGQGFPDYHAPKYALNALAAAANsaNQYTRGFGHPRLVQALSKLYsiNptEVLVTVGAYEALYATIQGHV<br>DE<br>G-----<br>DEVIIIEPFFDCYEPMVKAAGGIPRFIPLKPnsSADWVLDNNELEALFNEKTKMIIINTPHNPLGKVMDRAELEVVANLCKKWNVLCVSDEVYEHMV<br>FEpeHIRICTlmWERTITIGSAGKTFSLTGWKIGWAYGPEALL------<br>KNLQMVHQNCVYTCATPIQEAIAVGFEtkSPECYFNSISGELMAKRDYMASFLAEVGMNPTVPQGGYFMVADWSSLdtKWMTKSVGLQGIPPSA<br>FYSEPnlGEDFVRYCFFKKDENLQKAAEILRKWKG-<br>>6f77A_103 weight: 0.0054 score: 362.9 eval: 3.8e-11 prob: n/a identity: 0.2598 startpos: 3<br>LADALSRVKPSATIAVSQKARE-dVIGLGAGEPDFDTPDNIKKAAIDAIDreTKYTPVSGIPELREAIAKKfdYTAAQTIVGTGGKQILFNAFMATLNP---<br>--GDEVVIPAPYWVSYPEMVALCGGTPVFVPTRQEN--<br>NFKLKAEDLDRAITPKTKWFVFNSPSNPSGAAYSHEELKALTDVLMKhhVWVLTDDMYEHLTYGDffATPVEVeyERTLTMNGVSKAYAMTGWRI<br>GYAAGPLHLIKAMD------MIQGQQTSGAASIAQWAAVEALNG--<br>PQDFIGRNKEIFQGRRDLVVSMLNQagISCPTPEGAFYVYPSCAGlfVSELLETEGVAVVHGSAFGLG-----PNFRISYATSEALLEEACRRIQRFCAA<br>>1u08A_303 weight: 0.0041 score: 18 eval: n/a prob: n/a identity: 0.3071 startpos: 4<br>PQSKLPQLGTTIFTQMSALAQQHQAINLSQGFPDFDGPRYLQERLAHHVAqaNQYAPMTGVQALREAIAQKTepDAddITVTAGATEALYAAITA<br>LVRNG-----DEVICFDPSYDSYAPAIALSGGIVKRMALQPP---<br>HFRVDWQEFAALLSERTRLVILNTPHNPSATVWQQADFAALWQAIAGHEIFVISDEVYEHINFSqgHASVLAhlRERAVAVSSFGKTYHMTGWKV<br>GYCVAPAPISAEIRKVHQYLT------FSVNTPAQLALADMLRAEP--<br>EHYLALPDFYRQKRDILVNALNESRLEILPCEGTYFLLVDYSAVscQWLTQEHGVAAIPLSVFCADP-FPHKLIRLCFAKKESTLLAAAERLRQL---<br>>6l1nB_104 weight: 0.0037 score: 360 eval: 4.6e-11 prob: n/a identity: 0.2388 startpos: 5<br>PSDVIKTLPRQEFSLVFQKVKetHIINLGQGNPDLPTPPHIVEALREASLnfHGYGPFRGYPFLKEAIAAFytINptEVALFGGGKAGLYVLTQCLLNP----<br>-GDIALVPNPGYPEYLSGITMARAELYEMPLYEE--<br>NGYLPDFEKIDPAVLEKAKLMFLNYPNNPTGAVADAAFYAKAAAFAKEHNIHLIHDFAYGAFEFDQKPASFLEaaKTVGAELYSFSKTFNMAGWR<br>MAFAVGNEKIIQ------AVNEFQDHVFVGMFGGLQQAASAALSG--DPEHTESLKRIYKERIDFFTALCEKegWKMEKPKGTFYVWAEIP-<br>qFSDYLLEHAHVVVTPGEIFGSN---GKRHVRISMVSKQEDLREFVTRIQKLNLP<br>>6l1lB_105 weight: 0.0029 score: 355.7 eval: 6.3e-11 prob: n/a identity: 0.2388 startpos: 5<br>PSDVIKTLPRQEFSLVFQKVKeaHIINLGQGNPDLPTPPHIVEALREASLnfHGYGPFRGYPFLKEAIAAFytINptEVALFGGGKAGLYVLTQCLLNP----<br>-GDIALVPNPGYPEYLSGITMARAELYEMPLYEE--<br>NGYLPDFEKIDPAVLEKAKLMFLNYPNNPTGAVADAAFYAKAAAFAKEHNIHLIHDFAYGAFEFDQKPASFLEaaKTVGAELYSFSKTFNMAGWR<br>MAFAVGNEKIIQ------AVNEFQDHVFVGMFGGLQQAASAALSG--DPEHTESLKRIYKERIDFFTALCEKegWKMEKPKGTFYVWAEIP-<br>qFSDYLLEHAHVVVTPGEIFGSN---GKRHVRISMVSKQEDLREFVTRIQKLNLP<br>>4wp0B_304 weight: 0.0025 score: 17.71 eval: n/a prob: n/a identity: 0.3307 startpos: 3<br>QARRLDGIDYNPWVEFVKLASEHDVVNLGQGFPDFPPPDFAVEAFQHAVSglNQYTKTFGYPPLTKILASFFgiDprNVLVTVGGYGALFTAFQALV<br>DEG-----<br>DEVIIIEPFFDCYEPMTMMAGGRPVFVSLKPGGElnWQLDPMELAGKFTSRTKALVLNTPNNPLGKVFSREELELVASLCQQHDVVCITDEVYQW<br>MVYDGhhISIASlmWERTLTIGSAGKTFSATGWKVGWVLGPDHIMKHLRTVHQNS------<br>VFHCPTQSQAAVAESFEReqPSSYFVQFPQAMQRCRDHMIRSLQSVGLKPIIPQGSYFLITDIsrRFVKWMIKNKGLVAIPVSIFYSVPhhFDHYIRFC<br>FVKDEATLQAMDEKLRKWKVE<br>>6f77C_106 weight: 0.0025 score: 354.7 eval: 6.7e-11 prob: n/a identity: 0.2520 startpos: 14<br>=------------------------IGLGAGEPDFDTPDNIKKAAIDAIDreTKYTPVSGIPELREAIAKKfdYTAAQTIVGTGGKQILFNAFMATLNP-----<br>GDEVVIPAPYWVSYPEMVALCGGTPVFVPTRQE--<br>E4NNFKLKAEDLDRAITPKTKWFVFNSPSNPSGAAYSHEELKALTDVLMKhhVWVLTDDMYEHLTYGDffATPVEVeyERTLTMNGVSKAYAMTG<br>WRIGYAAGPLHLIKAMD------MIQGQQTSGAASIAQWAAVEALNG--<br>PQDFIGRNKEIFQGRRDLVVSMLNQagISCPTPEGAFYVYPSCAGlfVSELLETEGVAVVHGSAFGLG-----PNFRISYATSEALLEEACRRIQRFCAA<br>>2o0rA_203 weight: 0.0025 score: 1014 eval: 1e-45 prob: 100 identity: 0.4068 startpos: 2<br>TVSRLRPYATTVFAEMSALATRIGAVNLGQGFPDEDGPPKMLQAAQDAIagVNQYPPGPGSAPLRRAIAAQrdYDptEVLVTVGATEAIAAAVLGL<br>VEPG-----SEVLLIEPFYDSYSPVVAMAGAHRVTVPLVP-<br>DGRGFALDADALRRAVTPRTRALIINSPHNPTGAVLSATELAAIAEIAVAANLVVITDEVYEHLVFDhrHLPLAGfmAERTITISSAAKMFNCTGWKIG<br>WACGPAELI------AGVRAAKQYLSYVGGAPFQPAVALALDTE--<br>DAWVAALRNSLRARRDRLAAGLTEIGFAVHDSYGTYFLCADPRPLgcAALPEKVGVAAIPMSAFCDPADVWNHLVRFTFCKRDDTLDEAIRRLSVL<br>AE- |

| | | | | | | |
|---|---|---|---|---|---|---|
|  |  |  |  |  |  | >2o0rA_107 weight: 0.0022 score: 353.2 eval: 7.5e-11 prob: n/a identity: 0.4042 startpos: 11<br>---------TTVFAEMSALATRIGAVNLGQGFPDEDGPPKMLQAAQDAIagVNQYPPGPGSAPLRRAIAAQRryDptEVLVTVGATEAIAAAVLGLVEP--<br>---GSEVLLIEPFYDSYSPVVAMAGAHRVTVPLVPD-<br>GRGFALDADALRRAVTPRTRALIINSPHNPTGAVLSATELAAIAEIAVAANLVVITDEVYEHLVFDHahLPLAGfmAERTITISSAAKMFNCTGWKIG<br>WACGPA------ELIAGVRAAKQYLSYVGGAPFQPAVALALDT--<br>EDAWVAALRNSLRARRDRLAAGLTEIGFAVHDSYGTYFLCADPrtEFCAALPEKVGVAAIPMSAFCDPA-<br>vWNHLVRFTFCKRDDTLDEAIRRLSVLAE-<br>>1bkgA_110 weight: 0.0021 score: 347 eval: 1.2e-10 prob: n/a identity: 0.3333 startpos: 4<br>LSRRVQAMKPSATVAVNAKALE-<br>dLVALTAGEPDFDTPEHVKEAARRALagKTKYAPPAGIPELREALAEKFrvTPEETIVTVGGKQALFNLFQAILDP-----<br>GDEVIVLSPYWVSYPEMVRFAGGVVVEVETLPE--<br>EGFVPDPERVRRAITPRTKALVVNSPNNPTGAVYPKEVLEALARLAVEHDFYLVSDEIYEHLLYEGEHFSPGRVAPEHTLTVNGAAKAFAMTGWRI<br>GYACGPK------<br>EVIKAMASVSSQSTTSPDTIAQWATLEALTNqaSRAFVEMAREAYRRRRDLLLEGLTALGLKAVRPSGAFYVLMDTSPIaaAERLLEAGVAVVPGTD<br>FAA-----FGHVRLSYATSEENLRKALERFARVL--<br>>1o4sA_108 weight: 0.0021 score: 349.4 eval: 9.8e-11 prob: n/a identity: 0.2808 startpos: 1<br>VSRRISEIPISKTMELDAKAKAldVINLTAGEPDFPTPEPVVEEAVRFLqgEVKYTDPRGIYELREGIAKRIgiSPDQVVVTNGAKQALFNAFMALLDP---<br>--GDEVIVFSPVWVSYIPQIILAGGTVNVVETFMS--<br>KNFQPSLEEVEGLLVGKTKAVLINSPNNPTGVVYRREFLEGLVRLAKKRNFYIISDEVYDSLVYTDEFTSILDVsfDRIVYINGFSKSHSMTGWRVGYLIS<br>SE------KVATAVSKIQSHTTSCINTVAQYAALKALEV-----<br>DNSYMVQTFKERKNFVVERLKKMGVKFVEPEGAFYLFFKVRGdfCERLLEEKKVALVPGSAFLK-----PGFVRLSFATSIERLTEALDRIEDFLNS<br>>3fvxB_109 weight: 0.0021 score: 349.2 eval: 9.9e-11 prob: n/a identity: 0.3360 startpos: 3<br>QARRLDGIDYNPWVEFVKLASEHDVVNLGQGFPDFPPPDFAVEAFQHAVSglNQYTKTFGYPPLTKILASFFgiDprNVLVTVGGYGALFTAFQALV<br>D-----EGDEVIIIEPFFDCYEPMTMMAGGRPVFVSLKPGPI-<br>nWQLDPMELAGKFTSRTKALVLNTPNNPLGKVFSREELELVASLCQQHDVVCITDEVYQWMVYDGhhISIASLpwERTLTIGSAGKTFSATGWKV<br>GWVLGPDHIMK------HLRTVHQNSVFHCPTQSQAAVAESFEREQ-sSYFVQFPQAMQRCRDHMIRSLQSVGLKPIIPQGSYFLITDIS-<br>rFVKWMIKNKGLVAIPVSIFYSVPHqfDHYIRFCFVKDEATLQAMDEKLRKWKVE<br>>1w7lA_305 weight: 0.0018 score: 17.62 eval: n/a prob: n/a identity: 0.3333 startpos: 3<br>QARRLDGIDYNPWVEFVKLASEHDVVNLGQGFPDFPPPDFAVEAFQHAVSglNQYTKTFGYPPLTKILASFFgiDPlnVLVTVGGYGALFTAFQALV<br>DEG-----<br>DEVIIIEPFFDCYEPMTMMAGGRPVFVSLKPGPGElwQLDPMELAGKFTSRTKALVLNTPNNPLGKVFSREELELVASLCQQHDVVCITDEVYQW<br>MVYDGhhISIASlmWERTLTIGSAGKTFSATGWKVGWVLGPDHIMKHLRTVHQNS------<br>VFHCPTQSQAAVAESFEReqPSSYFVQFPQAMQRCRDHMIRSLQSVGLKPLIPQGSYFLITDIsrRFVKWMIKNKGLVAIPVSIFYSVpkHFDHYIRFC<br>FVKDEATLQAMDEKLRKWKVE<br>>1yiyA_306 weight: 0.0015 score: 17.52 eval: n/a prob: n/a identity: 0.2940 startpos: 4<br>LPKRYQGSTKSVWVEYIQLAAQYKPLNLGQGFPDYHAPKYALNALAAAANsaNQYTRGFGHPRLVQALSKLYsiNPmeVLVTVGAYEALYATIQGH<br>VDEG-----<br>DEVIIIEPFFDCYEPMVKAAGGIPRFIPLKPNKTGgwVLDNNELEALFNEKTKMIIINTPHNPLGKVMDRAELEVVANLCKKWNVLCVSDEVYEHMV<br>FEPfhIRICTlmWERTITIGSAGKTFSLTGWKIGWAYGPEALLKNLQMVHQNC------<br>VYTCATPIQEAIAVGFETesPECYFNSISGELMAKRDYMASFLAEVGMNPTVPQGGYFMVADWSSLdtKWMTKSVGLQGIPPSAFYSEPNKheDF<br>VRYCFFKKDENLQKAAEILRKWKGS<br>>4wp0D_307 weight: 0.0014 score: 17.5 eval: n/a prob: n/a identity: 0.3281 startpos: 3<br>QARRLDGIDYNPWVEFVKLASEHDVVNLGQGFPDFPPPDFAVEAFQHAVSglNQYTKTFGYPPLTKILASFFgiDPlnVLVTVGGYGALFTAFQALV<br>DEG-----<br>DEVIIIEPFFDCYEPMTMMAGGRPVFVSLKPGElsNWQLDPMELAGKFTSRTKALVLNTPNNPLGKVFSREELELVASLCQQHDVVCITDEVYQW<br>MVYDGhhISIASlmWERTLTIGSAGKTFSATGWKVGWVLGPDHIMKHLRTVHQNS------<br>VFHCPTQSQAAVAESFEReqPSSYFVQFPQAMQRCRDHMIRSLQSVGLKPIIPQGSYFLITDIsrRFVKWMIKNKGLVAIPVSIFYSVPhhFDHYIRFC<br>FVKDEATLQAMDEKLRKWKVE<br>>3fvxB_310 weight: 0.0013 score: 17.46 eval: n/a prob: n/a identity: 0.3307 startpos: 3<br>QARRLDGIDYNPWVEFVKLASEHDVVNLGQGFPDFPPPDFAVEAFQHAVSglNQYTKTFGYPPLTKILASFFgiDprNVLVTVGGYGALFTAFQALV<br>DEG-----<br>DEVIIIEPFFDCYEPMTMMAGGRPVFVSLKPGPIqnWQLDPMELAGKFTSRTKALVLNTPNNPLGKVFSREELELVASLCQQHDVVCITDEVYQW |

| | | | | | | |
|---|---|---|---|---|---|---|
|  |  |  |  |  |  | MVYDGhhISIASlmWERTLTIGSAGKTFSATGWKVGWVLGPDHIMKHLRTVHQNS------<br>VFHCPTQSQAAVAESFEReqPSSYFVQFPQAMQRCRDHMIRSLQSVGLKPIIPQGSYFLITDIsrRFVKWMIKNKGLVAIPVSIFYSVpkHFDHYIRFC<br>FVKDEATLQAMDEKLRKWKVE<br>>4wp0A_309 weight: 0.0013 score: 17.46 eval: n/a prob: n/a identity: 0.3281 startpos: 3<br>QARRLDGIDYNPWVEFVKLASEHDVVNLGQGFPDFPPPDFAVEAFQHAVSglNQYTKTFGYPPLTKILASFFgiDprNVLVTVGGYGALFTAFQALV<br>DEG-----<br>DEVIIIEPFFDCYEPMTMMAGGRPVFVSLKPGLgsNWQLDPMELAGKFTSRTKALVLNTPNNPLGKVFSREELELVASLCQQHDVVCITDEVYQW<br>MVYDGhhISIASlmWERTLTIGSAGKTFSATGWKVGWVLGPDHIMKHLRTVHQNS------<br>VFHCPTQSQAAVAESFEReqPSSYFVQFPQAMQRCRDHMIRSLQSVGLKPIIPQGSYFLITDIsrRFVKWMIKNKGLVAIPVSIFYSVpkHFDHYIRFC<br>FVKDEATLQAMDEKLRKWKVE<br>>3fvsA_308 weight: 0.0013 score: 17.49 eval: n/a prob: n/a identity: 0.3307 startpos: 3<br>QARRLDGIDYNPWVEFVKLASEHDVVNLGQGFPDFPPPDFAVEAFQHAVSglNQYTKTFGYPPLTKILASFFgiDPlnVLVTVGGYGALFTAFQALV<br>DEG-----<br>DEVIIIEPFFDCYEPMTMMAGGRPVFVSLKPGPIqnWQLDPMELAGKFTSRTKALVLNTPNNPLGKVFSREELELVASLCQQHDVVCITDEVYQW<br>MVYDGhhISIASlmWERTLTIGSAGKTFSATGWKVGWVLGPDHIMKHLRTVHQNS------<br>VFHCPTQSQAAVAESFEReqPSSYFVQFPQAMQRCRDHMIRSLQSVGLKPIIPQGSYFLITDIsrRFVKWMIKNKGLVAIPVSIFYSVpkHFDHYIRFC<br>FVKDEATLQAMDEKLRKWKVE<br>>3fvsA_204 weight: 0.0011 score: 964.58 eval: 1e-45 prob: 100 identity: 0.3360 startpos: 3<br>QARRLDGIDYNPWVEFVKLASEHDVVNLGQGFPDFPPPDFAVEAFQHAVsmLNQYTKTFGYPPLTKILASFfeIDprNVLVTVGGYGALFTAFQALV<br>DEG-----<br>DEVIIIEPFFDCYEPMTMMAGGRPVFVSLKPGPisNWQLDPMELAGKFTSRTKALVLNTPNNPLGKVFSREELELVASLCQQHDVVCITDEVYQW<br>MVYDgqHISIASlmWERTLTIGSAGKTFSATGWKVGWVLGPDHIM------<br>KHLRTVHQNSVFHCPTQSQAAVAESFEREqpSSYFVQFPQAMQRCRDHMIRSLQSVGLKPIIPQGSYFLITDISDFkvKWMIKNKGLVAIPVSIFYSV<br>PhhFDHYIRFCFVKDEATLQAMDEKLRKWKVE<br>>3fvxB_205 weight: 0.0005 score: 964.58 eval: 1e-45 prob: 100 identity: 0.3360 startpos: 3<br>QARRLDGIDYNPWVEFVKLASEHDVVNLGQGFPDFPPPDFAVEAFQHAVsmLNQYTKTFGYPPLTKILASFfeIDprNVLVTVGGYGALFTAFQALV<br>DEG-----<br>DEVIIIEPFFDCYEPMTMMAGGRPVFVSLKPGPisNWQLDPMELAGKFTSRTKALVLNTPNNPLGKVFSREELELVASLCQQHDVVCITDEVYQW<br>MVYDgqHISIASlmWERTLTIGSAGKTFSATGWKVGWVLGPDHIM------<br>KHLRTVHQNSVFHCPTQSQAAVAESFEREqpSSYFVQFPQAMQRCRDHMIRSLQSVGLKPIIPQGSYFLITDISDFkvKWMIKNKGLVAIPVSIFYSV<br>PhhFDHYIRFCFVKDEATLQAMDEKLRKWKVE<br>>1w7lA_206 weight: 0.0004 score: 971.9 eval: 1e-45 prob: 100 identity: 0.3386 startpos: 3<br>QARRLDGIDYNPWVEFVKLASEHDVVNLGQGFPDFPPPDFAVEAFQHAvgmLNQYTKTFGYPPLTKILASFfeIDprNVLVTVGGYGALFTAFQALV<br>DEG-----<br>DEVIIIEPFFDCYEPMTMMAGGRPVFVSLKPGpsSNWQLDPMELAGKFTSRTKALVLNTPNNPLGKVFSREELELVASLCQQHDVVCITDEVYQW<br>MVYDgqHISIASlmWERTLTIGSAGKTFSATGWKVGWVLGPDHIM------<br>KHLRTVHQNSVFHCPTQSQAAVAESFEREqpSSYFVQFPQAMQRCRDHMIRSLQSVGLKPLIPQGSYFLITDISDFkvKWMIKNKGLVAIPVSIFYSV<br>PhhFDHYIRFCFVKDEATLQAMDEKLRKWKVE<br>>5vehB_207 weight: 0.0003 score: 996.17 eval: 1e-45 prob: 100 identity: 0.2966 startpos: 4<br>LPKRYQGSTKSVWVEYIQLAAQYKPLNLGQGFPDYHAPKYALNALAAAaplANQYTRGFGHPRLVQALSKLytINPmeVLVTVGAYEALYATIQGH<br>VDEG-----<br>DEVIIIEPFFDCYEPMVKAAGGIPRFIPLKPnsSADWVLDNNELEALFNEKTKMIIINTPHNPLGKVMDRAELEVVANLCKKWNVLCVSDEVYEHMV<br>FEpeHIRICTlmWERTITIGSAGKTFSLTGWKIGWAYGPEALL------<br>KNLQMVHQNCVYTCATPIQEAIAVGFEtkSPECYFNSISGELMAKRDYMASFLAEVGMNPTVPQGGYFMVADWSSLatKWMTKSVGLQGIPPSA<br>FYSEPnlGEDFVRYCFFKKDENLQKAAEILRKWK-- |
| 2. | (WP_012693528.1) Pyridoxal phosphate dependent amino transferase | 388 | 6f35, 1bkg, 6f77, 1b5p, 1b5o, 5wml, 1gc3, 6l1n, 1j32, 6l1l, 5wmk, 5wmh, 5wmi, 2gb3, | *Deinococcus deserti* | 0.97 | MSGPFTLSQRALSLKPSATVAVTSRALELRRAGVDVISMSVGEPDFDTPEHIKAAAIHAIRSGQTKYTPVSGIPELREAISDKFRRENGLDYAPDAVT<br>VTSGGKQALFNAFFALLNPGDEVLIPAPYWVSYPEIVALTGAVPVPVQTSPDSGFLLDPEALEACVTSRTRMIVLNSPGNPTGAVFPPEVLQAVARI<br>AQRHNLIIVSDEMYEHLVYDAEQVSIGRYAPEHTLTVNGASKAYAMTGWRIGYAGGPKTVIAAMNALQSQSTSNASSVSQAAALAALVEHELTRK<br>FVDGALLAYRQRRDVIVAGLIDLGLQTPTPQGAFYVMANTTSIHADELEAARIILDQARVAVVPGTDFSAPGQVRLSYATSLENIHEVLRRLRTLTRP<br>>6f77A_201 weight: 0.3513 score: 983.71 eval: 1e-45 prob: 100 identity: 0.4742 startpos: 1<br>----<br>AFLADALSRVKPSATIAVSQKARELKAKGRDVIGLGAGEPDFDTPDNIKKAAIDAIDRGETKYTPVSGIPELREAIAKKFKRENNLDYTAAQTIVGTGG |

| | | | | | | |
|---|---|---|---|---|---|---|
|  |  |  |  |  |  | KQILFNAFMATLNPGDEVVIPAPYWVSYPEMVALCGGTPVFVPTRQENNFKLKAEDLDRAITPKTKWFVFNSPSNPSGAAYSHEELKALTDVLMK<br>hhVWVLTDDMYEHLTYGdrFATPVEvlYERTLTMNGVSKAYAMTGWRIGYAAGPLHLIKAMDMIQGQQTSGAASIAQWAAVEALNGPQ---<br>DFIGRNKEIFQGRRDLVVSMLNQagISCPTPEGAFYVYPSCAGLieTDEDFVSELLETEGVAVVHGSAFGLGPNFRISYATSEALLEEACRRIQRFCAA<br>>6f77A_101 weight: 0.3006 score: 437.6 eval: 6.5e-12 prob: n/a identity: 0.4742 startpos: 2<br>-----<br>FLADALSRVKPSATIAVSQKARELKAKGRDVIGLGAGEPDFDTPDNIKKAAIDAIDRGETKYTPVSGIPELREAIAKKFKRENNLDYTAAQTIVGTGGK<br>QILFNAFMATLNPGDEVVIPAPYWVSYPEMVALCGGTPVFVPTRQENNFKLKAEDLDRAITPKTKWFVFNSPSNPSGAAYSHEELKALTDVLMKh<br>hVWVLTDDMYEHLTYGDffATPVEVeyERTLTMNGVSKAYAMTGWRIGYAAGPLHLIKAMDMIQGQQTSGAASIAQWAAVEALNGP---<br>QDFIGRNKEIFQGRRDLVVSMLNQagISCPTPEGAFYVYPSCAGLieTDEDFVSELLETEGVAVVHGSAFGLGPNFRISYATSEALLEEACRRIQRFCA<br>A<br>>6f35A_301 weight: 0.2859 score: 24.2 eval: n/a prob: n/a identity: 0.4510 startpos: 1<br>---<br>GFQPASRISSIGVSEILKIGARAAAMKREGKPVIILGAGEPDFDTPEHVKQAASDAIHRGETKYTALDGTPELKKAIREKFQRENGLAYELDEITVATG<br>AKQILFNAMMASLDPGDEVIIPTPYWTSYSDIVHICEGKPVLIACDASSGFRLTAEKLEAAITPRTRWVLLNSPSNPSGAAYSAADYRPLLEVLLRHpv<br>WLLVDDMYEHIVYDgrFVTPAQLEpnRTLTVNGVSKAYAMTGWRIGYAGGPRELIKAMAVVQSQATSCPSSISQAASVVALNGP---<br>QDFLKERTESFQRRRDLVVNGLNAigLDCRVPEGAFYTFSGCAGVLgtDTDFCAYLLEDAHVAVVPGSAFGLSPFFRISYATSEAELKEALERIAAACD<br>R<br>>6f35A_102 weight: 0.0083 score: 437.4 eval: 6.6e-12 prob: n/a identity: 0.4459 startpos: 3<br>-----<br>QPASRISSIGVSEILKIGARAAAMKREGKPVIILGAGEPDFDTPEHVKQAASDAIHRGETKYTALDGTPELKKAIREKFQRENGLAYELDEITVATGAK<br>QILFNAMMASLDPGDEVIIPTPYWTSYSDIVHICEGKPVLIACDASSGFRLTAEKLEAAITPRTRWVLLNSPSNPSGAAYSAADYRPLLEVLLRhhVW<br>LLVDDMYEHIVYDGffVTPAQLekNRTLTVNGVSKAYAMTGWRIGYAGGPRELIKAMAVVQSQATSCPSSISQAASVVALNGPQ---<br>DFLKERTESFQRRRDLVVNGLNAigLDCRVPEGAFYTFSGCAGVlkTDTDFCAYLLEDAHVAVVPGSAFGLSPFFRISYATSEAELKEALERIAAACDR<br>>1bkgA_202 weight: 0.0069 score: 846.87 eval: 1e-45 prob: 100 identity: 0.5722 startpos: 1<br>MR---<br>GLSRRVQAMKPSATVAVNAKALELRRQGVDLVALTAGEPDFDTPEHVKEAARRALAQGKTKYAPPAGIPELREALAEKFRRENGLSVTPEETIVTV<br>GGKQALFNLFQAILDPGDEVIVLSPYWVSYPEMVRFAGGVVVEVETLPEEGFVPDPERVRRAITPRTKALVVNSPNNPTGAVYPKEVLEALARLAV<br>EHDFYLVSDEIYEHLLYEGEHFSPGRVAPEHTLTVNGAAKAFAMTGWRIGYACGPKEVIKAMASVSSQSTTSPDTIAQWATLEALTNQEASRAFVE<br>MAREAYRRRRDLLLEGLTALGLKAVRPSGAFYVLMDTSPIAPDEVRAAERLLE-AGVAVVPGTDFAAFGHVRLSYATSEENLRKALERFARVL--<br>>6f77A_302 weight: 0.0067 score: 24.16 eval: n/a prob: n/a identity: 0.4768 startpos: 1<br>----<br>AFLADALSRVKPSATIAVSQKARELKAKGRDVIGLGAGEPDFDTPDNIKKAAIDAIDRGETKYTPVSGIPELREAIAKKFKRENNLDYTAAQTIVGTGG<br>KQILFNAFMATLNPGDEVVIPAPYWVSYPEMVALCGGTPVFVPTRQENNFKLKAEDLDRAITPKTKWFVFNSPSNPSGAAYSHEELKALTDVLMK<br>HpvWVLTDDMYEHLTYGdrFATPVEVEpeRTLTMNGVSKAYAMTGWRIGYAAGPLHLIKAMDMIQGQQTSGAASIAQWAAVEALNGP---<br>QDFIGRNKEIFQGRRDLVVSMLNQagISCPTPEGAFYVYPSCAGliETDEDFVSELLETEGVAVVHGSAFGLGPNFRISYATSEALLEEACRRIQRFCAA<br>>1bkgA_103 weight: 0.0050 score: 422.7 eval: 1.7e-11 prob: n/a identity: 0.5696 startpos: 3<br>-----<br>GLSRRVQAMKPSATVAVNAKALELRRQGVDLVALTAGEPDFDTPEHVKEAARRALAQGKTKYAPPAGIPELREALAEKFRRENGLSVTPEETIVTV<br>GGKQALFNLFQAILDPGDEVIVLSPYWVSYPEMVRFAGGVVVEVETLPEEGFVPDPERVRRAITPRTKALVVNSPNNPTGAVYPKEVLEALARLAV<br>EHDFYLVSDEIYEHLLYEGEHFSPGRVAPEHTLTVNGAAKAFAMTGWRIGYACGPKEVIKAMASVSSQSTTSPDTIAQWATLEALTNQEASRAFVE<br>MAREAYRRRRDLLLEGLTALGLKAVRPSGAFYVLMDTSPIAPDEVRAAERLLE-AGVAVVPGTDFAAFGHVRLSYATSEENLRKALERFARVL--<br>>6f77C_303 weight: 0.0035 score: 22.62 eval: n/a prob: n/a identity: 0.4510 startpos: 1<br>=---------------------AFLADALSRVKP-<br>VIGLGAGEPDFDTPDNIKKAAIDAIDRGETKYTPVSGIPELREAIAKKFKRENNLDYTAAQTIVGTGGKQILFNAFMATLNPGDEVVIPAPYWVSYPE<br>MVALCGGTPVFVPTRQENNFKLKAEDLDRAITPKTKWFVFNSPSNPSGAAYSHEELKALTDVLMKHpvWVLTDDMYEHLTYGdrFATPVEVEpeR<br>TLTMNGVSKAYAMTGWRIGYAAGPLHLIKAMDMIQGQQTSGAASIAQWAAVEALNGP---<br>QDFIGRNKEIFQGRRDLVVSMLNQagISCPTPEGAFYVYPSCAGliETDEDFVSELLETEGVAVVHGSAFGLGPNFRISYATSEALLEEACRRIQRFCAA<br>>1b5pA_104 weight: 0.0034 score: 420.1 eval: 1.9e-11 prob: n/a identity: 0.5644 startpos: 3<br>-----<br>GLSRRVQAMKPSATVAVNAKALELRRQGVDLVALTAGEPDFDTPEHVKEAARRALAQGKTKYAPPAGIPELREALAEKFRRENGLSVTPEETIVTV<br>GGSQALFNLFQAILDPGDEVIVLSPYWVSYPEMVRFAGGVVVEVETLPEEGFVPDPERVRRAITPRTKALVVNSPNNPTGAVYPKEVLEALARLAVE |

| | | | | | | |
|---|---|---|---|---|---|---|
|  |  |  |  |  |  | HDFYLVSDEIYEHLLYEGEHFSPGRVAPEHTLTVNGAAKAFAMTGWRIGYACGPKEVIKAMASVSRQSTTSPDTIAQWATLEALTNQEASRAFVE<br>MAREAYRRRRDLLLEGLTALGLKAVRPSGAFYVLMDTSPIAPDEVRAAERLLE-AGVAVVPGTDFAAFGHVRLSYATSEENLRKALERFARVL--<br>>1b5oA_203 weight: 0.0028 score: 910.11 eval: 1e-45 prob: 100 identity: 0.5670 startpos: 1<br>---<br>MRGLSRRVQAMKPSATVAVNAKALELRRQGVDLVALTAGEPDFDTPEHVKEAARRALAQGKTKYAPPAGIPELREALAEKFRRENGLSVTPEETIV<br>TVGGSQALFNLFQAILDPGDEVIVLSPYWVSYPEMVRFAGGVVVEVETLPEEGFVPDPERVRRAITPRTKALVVNSPNNPTGAVYPKEVLEALARLA<br>VEHDFYLVSDEIYEHLLYEGEHFSPGRVAPEHTLTVNGAAKAFAMTGWRIGYACGPKEVIKAMASVSSQSTTSPDTIAQWATLEALTNQEASRAFV<br>EMAREAYRRRRDLLLEGLTALGLKAVRPSGAFYVLMDTSPIAPDEVRAAERLL-EAGVAVVPGTDFAAFGHVRLSYATSEENLRKALERFARVL--<br>>1b5oA_105 weight: 0.0027 score: 417.2 eval: 2.3e-11 prob: n/a identity: 0.5670 startpos: 3<br>-----<br>GLSRRVQAMKPSATVAVNAKALELRRQGVDLVALTAGEPDFDTPEHVKEAARRALAQGKTKYAPPAGIPELREALAEKFRRENGLSVTPEETIVTV<br>GGSQALFNLFQAILDPGDEVIVLSPYWVSYPEMVRFAGGVVVEVETLPEEGFVPDPERVRRAITPRTKALVVNSPNNPTGAVYPKEVLEALARLAVE<br>HDFYLVSDEIYEHLLYEGEHFSPGRVAPEHTLTVNGAAKAFAMTGWRIGYACGPKEVIKAMASVSSQSTTSPDTIAQWATLEALTNQEASRAFVE<br>MAREAYRRRRDLLLEGLTALGLKAVRPSGAFYVLMDTSPIAPDEVRAAERLLE-AGVAVVPGTDFAAFGHVRLSYATSEENLRKALERFARVL--<br>>6f77C_106 weight: 0.0023 score: 414.4 eval: 2.8e-11 prob: n/a identity: 0.4407 startpos: 1<br>=----------------------AFLADALSRVK-<br>vIGLGAGEPDFDTPDNIKKAAIDAIDRGETKYTPVSGIPELREAIAKKFKRENNLDYTAAQTIVGTGGKQILFNAFMATLNPGDEVVIPAPYWVSYPE<br>MVALCGGTPVFVPTRQENNFKLKAEDLDRAITPKTKWFVFNSPSNPSGAAYSHEELKALTDVLMKhhVWVLTDDMYEHLTYGDffATPVEVeyER<br>TLTMNGVSKAYAMTGWRIGYAAGPLHLIKAMDMIQGQQTSGAASIAQWAAVEALNGP---<br>QDFIGRNKEIFQGRRDLVVSMLNQagISCPTPEGAFYVYPSCAGliETDEDFVSELLETEGVAVVHGSAFGLGPNFRISYATSEALLEEACRRIQRFCAA<br>>5wmlB_304 weight: 0.0022 score: 22.03 eval: n/a prob: n/a identity: 0.4613 startpos: 1<br>-----<br>SLSPRVQSLKPSKTMVITDLAATLVQSGVPVIRLAAGEPDFDTPKVVAEAGINAIREGFTRYTLNAGITELREAICRKLKEENGLSYAPDQILVSNGAK<br>QSLLQAVLAVCSPGDEVIIPAPYWVSYTEQARLADATPVVIPTKISNNFLLDPKDLESKLTEKSRLLILCSPSNPTGSVYPKSLLEEIARIIAKhrLLVLSDEI<br>YEHIIYAptHTSFASlmYERTLTVNGFSAAFAMTGWRLGYLAGPKHIVAACSKLQGQVSSGASSIAQKAGVAALGLGKAGGETVAEMVKAYRERRD<br>FLVKSLGDigVKISEPQGAFYLFIDFSAYynDSSSLALYFLDKFQVAMVPGDAFGDDSCIRISYATSLDVLQAAVEKIRKALEP<br>>1gc3A_107 weight: 0.0021 score: 405.8 eval: 4.7e-11 prob: n/a identity: 0.5593 startpos: 3<br>-----<br>GLSRRVQAMKPDAVVAVNAKALELRRQGVDLVALTAGEPDFDTPEHVKEAARRALAQGKTKYAPPAGIPELREALAEKFRRENGLSVTPEETIVTV<br>GGSQALFNLFQAILDPGDEVIVLSPYWVSYPEMVRFAGGVVVEVETLPEEGFVPDPERVRRAITPRTKALVVNSPNNPTGAVYPKEVLEALARLAVE<br>HDFYLVSDEIYEHLLYEGEHFSPGRVAPEHTLTVNGAAKAFAMTGWRIGYACGPKEVIKAMASVSRQSTTSPDTIAQWATLEALTNQEASRAFVE<br>MAREAYRRRRDLLLEGLTALGLKAVRPSGAFYVLMDTSPIAPDEVRAAERLLE-AGVAVVPGTDFAAFGHVRLSYATSEENLRKALERFARVL--<br>>6l1nB_108 weight: 0.0020 score: 403.5 eval: 5.4e-11 prob: n/a identity: 0.2835 startpos: 4<br>=-----TPSDVIKTLPRQEFSLVFQKVKEME--<br>KTHIINLGQGNPDLPTPPHIVEALREASLNpfHGYGPFRGYPFLKEAIAAFYKREYGVTINptEVALFGGGKAGLYVLTQCLLNPGDIALVPNPGYPEYL<br>SGITMARAELYEMPLYEENGYLPDFEKIDPAVLEKAKLMFLNYPNNPTGAVADAAFYAKAAAFAKEHNIHLIHDFAYGAFEFDQKPASFLEaaKTVG<br>AELYSFSKTFNMAGWRMAFAVGNEKIIQAVNEFQDHVFVGMFGGLQQAASAALSG---<br>DPEHTESLKRIYKERIDFFTALCEKegWKMEKPKGTFYVWAEIPNTFETSHQFSDYLLEHAHVVVTPGEIFGskRHVRISMVSKQEDLREFVTRIQKLN<br>LP<br>>1j32A_109 weight: 0.0020 score: 400.9 eval: 6.4e-11 prob: n/a identity: 0.4072 startpos: 2<br>-----<br>KLAARVESVSPSMTLIIDAKAKAMKAEGIDVCSFSAGEPDFNTPKHIVEAAKAALEQGKTRYGPAAGEPRLREAIAQKLQRDNGLCYGADNILVTN<br>GGKQSIFNLMLAMIEPGDEVIIPAPFWVSYPEMVKLAEGTPVILPTTVETQFKVSPEQIRQAITPKTKLLVFNTPSNPTGMVYTPDEVRAIAQVAVE<br>AGLWVLSDEIYEKILYDDahLSIGAASpeRSVVCSGFAKTYAMTGWRVGFLAGPVPLVKAATKIQGHSTSNVCTFAQYGAIAAYEN---<br>SQDCVQEMLAAFAERRRYMLDALNAmgLECPKPDGAFYMFPSIAKTGRSSLDFCSELLDQHQVATVPGAAFGADDCIRLSYATDLDTIKRGMERL<br>EKFLHG<br>>6l1lB_110 weight: 0.0019 score: 400.6 eval: 6.5e-11 prob: n/a identity: 0.2861 startpos: 4<br>-----<br>TPSDVIKTLPRQEFSLVFQKVKEMEKTGAHIINLGQGNPDLPTPPHIVEALREASLnsFHGYGPFRGYPFLKEAIAAFYKREYGVTINptEVALFGGGK<br>AGLYVLTQCLLNPGDIALVPNPGYPEYLSGITMARAELYEMPLYEENGYLPDFEKIDPAVLEKAKLMFLNYPNNPTGAVADAAFYAKAAAFAKEHNI<br>HLIHDFAYGAFEFDQKPASFLEaaKTVGAELYSFSKTFNMAGWRMAFAVGNEKIIQAVNEFQDHVFVGMFGGLQQAASAALSGD--- |

| | | | | | | |
|---|---|---|---|---|---|---|
|  |  |  |  |  |  | PEHTESLKRIYKERIDFFTALCEKegWKMEKPKGTFYVWAEIPNTFETSHQFSDYLLEHAHVVVTPGEIFGskRHVRISMVSKQEDLREFVTRIQKLNL<br>P<br>>5wmkA_305 weight: 0.0016 score: 21.89 eval: n/a prob: n/a identity: 0.4613 startpos: 1<br>-----<br>SLSPRVQSLKPSKVMVITDLAATLVQSGVPVIRLAAGEPDFDTPKVVAEAGINAIREGFTRYTLNAGITELREAICRKLKEENGLSYAPDQILVSNGAV<br>QSLLQAVLAVCSPGDEVIIPAPYWVSYTEQARLADATPVVIPTKISNNFLLDPKDLESKLTEKSRLLILCSPSNPTGSVYPKSLLEEIARIIAKhrLLVLSDEI<br>YEHIIYaaTHTSFASLpyERTLTVNGFSKAFAMTGWRLGYLAGPKHIVAACSKLQGQVSSGASSIAQKAGVAALGLGKAGGETVAEMVKAYRERRD<br>FLVKSLGDigVKISEPQGAFYLFIDFSAYynDSSSLALYFLDKFQVAMVPGDAFGDDSCIRISYATSLDVLQAAVEKIRKALEP<br>>5wmhC_306 weight: 0.0014 score: 21.83 eval: n/a prob: n/a identity: 0.4665 startpos: 1<br>---<br>DMSLSPRVQSLKPSKTMVITDLAATLVQSGVPVIRLAAGEPDFDTPKVVAEAGINAIREGFTRYTLNAGITELREAICRKLKEENGLSYAPDQILVSNG<br>AKQSLLQAVLAVCSPGDEVIIPAPYWVSYTEQARLADATPVVIPTKISNNFLLDPKDLESKLTEKSRLLILCSPSNPTGSVYPKSLLEEIARIIAKHplLVLS<br>DEIYEHIIYaaTHTSFASlmYERTLTVNGFSKAFAMTGWRLGYLAGPKHIVAACSKLQGQVSSGASSIAQKAGVAALGLGKAGGETVAEMVKAYRER<br>RDFLVKSLGDigVKISEPQGAFYLFIDFSAyiNDSSSLALYFLDKFQVAMVPGDAFGDDSCIRISYATSLDVLQAAVEKIRKALEP<br>>5wmlA_307 weight: 0.0013 score: 21.8 eval: n/a prob: n/a identity: 0.4613 startpos: 1<br>----<br>MSLSPRVQSLKPSKTMVITDLAATLVQSGVPVIRLAAGEPDFDTPKVVAEAGINAIREGFTRYTLNAGITELREAICRKLKEENGLSYAPDQILVSNGA<br>KQSLLQAVLAVCSPGDEVIIPAPYWVSYTEQARLADATPVVIPTKISNNFLLDPKDLESKLTEKSRLLILCSPSNPTGSVYPKSLLEEIARIIAKhrLLVLSD<br>EIYEHIIYAptHTSFASLpyERTLTVNGFSAAFAMTGWRLGYLAGPKHIVAACSKLQGQVSSGASSIAQKAGVAALGLGKAGGETVAEMVKAYRERR<br>DFLVKSLGDigVKISEPQGAFYLFIDFSAyiNDSSSLALYFLDKFQVAMVPGDAFGDDSCIRISYATSLDVLQAAVEKIRKALEP<br>>5wmiA_309 weight: 0.0012 score: 21.53 eval: n/a prob: n/a identity: 0.4613 startpos: 1<br>----MSLSPRVQSLKPSKVMVITDLAATLVS-<br>GVPVIRLAAGEPDFDTPKVVAEAGINAIREGFTRYTLNAGITELREAICRKLKEENGLSYAPDQILVSNGAKQSLLQAVLAVCSPGDEVIIPAPYWVSY<br>TEQARLADATPVVIPTKISNNFLLDPKDLESKLTEKSRLLILCSPSNPTGSVYPKSLLEEIARIIAKHplLVLSDEIYEHIIYAptHTSFASLpyERTLTVNGFSK<br>AFAMTGWRLGYLAGPKHIVAACSKLQGQVSSGASSIAQKAGVAALGLGKAGGETVAEMVKAYRERRDFLVKSLGDigVKISEPQGAFYLFIDFSAY<br>ynDSSSLALYFLDKFQVAMVPGDAFGDDSCIRISYATSLDVLQAAVEKIRKALEP<br>>5wmhA_308 weight: 0.0012 score: 21.78 eval: n/a prob: n/a identity: 0.4639 startpos: 1<br>-----<br>SLSPRVQSLKPSKTMVITDLAATLVQSGVPVIRLAAGEPDFDTPKVVAEAGINAIREGFTRYTLNAGITELREAICRKLKEENGLSYAPDQILVSNGAK<br>QSLLQAVLAVCSPGDEVIIPAPYWVSYTEQARLADATPVVIPTKISNNFLLDPKDLESKLTEKSRLLILCSPSNPTGSVYPKSLLEEIARIIAKHplLVLSDEI<br>YEHIIYAptHTSFASlmYERTLTVNGFSKAFAMTGWRLGYLAGPKHIVAACSKLQGQVSSGASSIAQKAGVAALGLGKAGGETVAEMVKAYRERRD<br>FLVKSLGDigVKISEPQGAFYLFIDFSAYynDSSSLALYFLDKFQVAMVPGDAFGDDSCIRISYATSLDVLQAAVEKIRKALEP<br>>1j32A_310 weight: 0.0012 score: 20.6 eval: n/a prob: n/a identity: 0.4098 startpos: 1<br>----<br>MKLAARVESVSPSMTLIIDAKAKAMKAEGIDVCSFSAGEPDFNTPKHIVEAAKAALEQGKTRYGPAAGEPRLREAIAQKLQRDNGLCYGADNILVT<br>NGGKQSIFNLMLAMIEPGDEVIIPAPFWVSYPEMVKLAEGTPVILPTTVETQFKVSPEQIRQAITPKTKLLVFNTPSNPTGMVYTPDEVRAIAQVAV<br>EAGLWVLSDEIYEKILYdaQHLSIGAASpeRSVVCSGFAKTYAMTGWRVGFLAGPVPLVKAATKIQGHSTSNVCTFAQYGAIAAYENS---<br>QDCVQEMLAAFAERRRYMLDALNAmgLECPKPDGAFYMFPSIAKTGRSSLDFCSELLDQHQVATVPGAAFGADDCIRLSYATDLDTIKRGMERLE<br>KFLHG<br>>1j32A_204 weight: 0.0012 score: 1035.58 eval: 1e-45 prob: 100 identity: 0.4046 startpos: 1<br>----<br>MKLAARVESVSPSMTLIIDAKAKAMKAEGIDVCSFSAGEPDFNTPKHIVEAAKAALEQGKTRYGPAAGEPRLREAIAQKLQRDNGLCYGADNILVT<br>NGGKQSIFNLMLAMIEPGDEVIIPAPFWVSYPEMVKLAEGTPVILPTTVETQFKVSPEQIRQAITPKTKLLVFNTPSNPTGMVYTPDEVRAIAQVAV<br>EAGLWVLSDEIYEKILYDDahLSIGAAsyERSVVCSGFAKTYAMTGWRVGFLAGPVPLVKAATKIQGHSTSNVCTFAQYGAIAAYENSQ---<br>DCVQEMLAAFAERRRYMLDALNAmgLECPKPDGAFYMFPSIAKTGRSSLDFCSELLDQHQVATVPGAAFGADDCIRLSYATDLDTIKRGMERLEK<br>FLHG<br>>6f35A_205 weight: 0.0006 score: 902 eval: 1e-45 prob: 100 identity: 0.4536 startpos: 1<br>=---<br>GFQPASRISSIGVSEILKIGARAAAMKREGKPVIILGAGEPDFDTPEHVKQAASDAIHRGETKYTALDGTPELKKAIREKFQRENGLAYELDEITVATG<br>AKQILFNAMMASLDPGDEVIIPTPYWTSYSDIVHICEGKPVLIACDASSGFRLTAEKLEAAITPRTRWVLLNSPSNPSGAAYSAADYRPLLEVLLRhhV<br>WLLVDDMYEHIVYDgaQLEPGLKN--RTLTVNGVSKAYAMTGWRIGYAGGPRELIKAMAVVQSQATSCPSSISQAASVVALNGPQD---<br>FLKERTESFQRRRDLVVNGLNAigLDCRVPEGAFYTFSggKRIK-TDTDFCAYLLEDAHVAVVPGSAFGLSPFFRISYATSEAELKEALERIAAACDR |

| | | | | | | |
|---|---|---|---|---|---|---|
| | | | | | | >6l1lB_206 weight: 0.0004 score: 472.03 eval: 1e-45 prob: 100 identity: 0.2861 startpos: 1<br>--<br>MEITPSDVIKTLPRQEFSLVFQKVKEMEKTGAHIINLGQGNPDLPTPPHIVEALREASLNpfHGYGPFRGYPFLKEAIAAFYKREYGVTINPeeVALFG<br>GGKAGLYVLTQCLLNPGDIALVPNPGYPEYLSGITMARAELYEMPLYEENGYLPDFEKIDPAVLEKAKLMFLNYPNNPTGAVADAAFYAKAAAFAK<br>EHNIHLIHDFAYGAFEFDQKPASFLEAekTVGAELYSFSKTFNMAGWRMAFAVGNEKIIQAVNEFQDHVFVGMFGGLQQAASAALSGD---<br>PEHTESLKRIYKERIDFFTALCeeLGWKMEKPKGTFYVWAEIPNTFETSHQFSDYLLEHAHVVVTPGEIFGskRHVRISMVSKQEDLREFVTRIQKLNL<br>P<br>>2gb3B_207 weight: 0.0003 score: 798.52 eval: 1e-45 prob: 100 identity: 0.2577 startpos: 1<br>=-HHMDVFSDRVLLTEESPIRKLVPFAEMAKKRGVRIHHLNIGQPDLKTPEVFFERIYE--<br>NKPEvyYSHSAGIWELREAFASYYKRRQRVDVKPENVLVTNGGSEAILFSFAVIANPGDEILVLEPFYANYNAFAKIAGVKLIPVTRRMEEGFAI-<br>PQNLESFINERTKGIVLSNPCNPTGVVYGKDEMRYLVEIAERHGLFLIVDEVYSEIVFRGEFASALSIESDKVVVIDSVSKKFSACGARVGCLITreELISH<br>AMK-LAQGRLAPPLLEQIGSVGLLNLDD---SFFDFVRETYRERVETVLKKLEEhlKRFTKPSGAFYITAELPVE--<br>DAEEFARWMLTdeTTMVAPLRGFytkKEIRIACVLEKDLLSRAIDVLMEGLKM |
| 3. | (NP_295136.1) Acetyl-lysine deacetylase | 361 | 3pfo, 5xoy, 4q7a, 5vo3, 3isz, 4o23, 3gb0, 1vgy, 3ic1, 3ct9, 5k8p, 2rb7, 5k8n, 5k8m | *Deinococcus radiodurans* | 0.86 | MTPTPDAQAARELIRQAVSIPSLSGEEQAIAAFLRDWMARRGFDAQVDEAGNAVGVRGSGPLTVALLGHMDTVPGDIPVRVDEAGVLHGRGSV<br>DAKGSLCTFIAAVSALPPEALSAARFVCIGATEEEAPSSKGARYAMRQHRPDFVLIGEPSGWAGLTLGYKGRLVAKVRVEKDNFHTAGDGTSAADD<br>LTLGWQRVREWAAGFAPADSGGGGIFDRVQVTLQDLGSSGDGLTQRAWATIGLRLPPALAPYQAEEAIEQAFAGLGADLTFTGHESAVRHPKD<br>NALTRALRVAIREQGGTPTFKVKTGTSDMNVVAELWPVPTLAYGPGDSALDHTPEERLDLAEYDRAVAVLTSALTRLVGG<br>>3pfoA_201 weight: 0.3364 score: 484.37 eval: 1e-45 prob: 100 identity: 0.2182 startpos: 13<br>=-AVDRNFNDQVAFLQRMVQFRSVRGEEAPQQEWLAQQFADRGYKVDtaGSMQVVATADskGRSLILQGHIDVVPEgyEAKVR-<br>DGWMIGRGAQDMKGGVSAMIFALDAIRTayAPDARVHVQTVTEEE-STGNGALSTLMryRADACLIPEPTGH-<br>TLTRAQVGAVWFRLRVRGTPVHVAYSEtsAILSAMHLIRAFEEYTKELNAQ-<br>AVRDPWfnPIKFNVGIIKgwASSTAAWCELDCRLGLLTGDTPQEAMRGIEKCLAdnPAELVWsfQADPAVCEPGGVAEDVLTAAHKAanAPLDARL<br>STAVNDTRYYSVDYGIPALCYGPYGQG-PHAFDERIDLESLRKTTLSIALFVAEWCGL<br>>5xoyB_101 weight: 0.3027 score: 385.5 eval: 3.1e-13 prob: n/a identity: 0.4641 startpos: 1<br>=-------LDPVEFLKGALEIPSPSGKERLVAEYLAEGMQKLGLKGFVDEADNARGQVGEGPVQVVLLGHIDTVPGQIPVRLE-<br>GGRLFGRGAVDAKGPFVAMIFAAAGLSEEARKRLTVHLVGATEEEAPSSKGARFVAPRLKPHYAVIGEPSGWEGITLGYKGRLLVKARREKDHFHS<br>AHHEPNAAEELISYFVAIKAWAEAMN----VGQRPFDQVQYTLRDFRVHPAELRQVAEMFFDLRLPPRLPPEEAIRHLTAYA-<br>PPTIELEFFGREVPYQGPKDTPLTRAFRQAIRKAGGRPVFKLKTGTSDMNVLAPHWPVPMVAYGPGDSTLDHTPYEHVEVAEFLKGIEVLRGALEA<br>LAQT<br>>5xoyB_301 weight: 0.2958 score: 23.76 eval: n/a prob: n/a identity: 0.4669 startpos: 1<br>=-------LDPVEFLKGALEIPSPSGKERLVAEYLAEGMQKLGLKGFVDEADNARGQVGEGPVQVVLLGHIDTVPGQIPVR-<br>LEGGRLFGRGAVDAKGPFVAMIFAAAGLSEEARKRLTVHLVGATEEEAPSSKGARFVAPRLKPHYAVIGEPSGWEGITLGYKGRLLVKARREKDHF<br>HSAHHEPNAAEELISYFVAIKAWAEAMNV----GQRPFDQVQYTLRDFRVHPAELRQVAEMFFDLRLPPRLPPEEAIRHLTAYAPPT-<br>IELEFFGREVPYQGPKDTPLTRAFRQAIRKAGGRPVFKLKTGTSDMNVLAPHWPVPMVAYGPGDSTLDHTPYEHVEVAEFLKGIEVLRGALEALAQ<br>T<br>>4q7aA_102 weight: 0.0087 score: 377.4 eval: 5.8e-13 prob: n/a identity: 0.4448 startpos: 9<br>=------SMTTADLLRGLVSIPSPSGAEAPAVEWLCQQMAALGYQAEPDGAGNAVGTRGEGPREIMLLGHIDTVPGEVPVQVV-<br>DGVLYGRGAVDAKGPLATFVVAGARAK--<br>LPPGVRLTVVGAVEEEVMSSRGARHLIAteAPDAVVIGEPSGWDGVVLGYRGSVALEYRVTVPMSHSAGPEATAAELAADFWYRLRTWCAEWS-<br>--VGIDHAFHRVEPKLNALNSSSDGLYGEAVARIGLRLPPALSPEEAIAVATSLAS--<br>EGEVTATVNAPAFQTDKRQPIVAAFLAAVRAHGGTPRLKLKTGTSDMNLVGPAWGCPIVAYGPGDSRLDHTPEEHVPLADLERATAILTTAIERVA<br>AQ<br>>4q7aA_302 weight: 0.0074 score: 23.39 eval: n/a prob: n/a identity: 0.4475 startpos: 1<br>MLESISPMstADLLRGLVSIPSPSGAEAPAVEWLCQQMAALGYQAEPDGAGNAVGTRGEGPREIMLLGHIDTVPGEVPVQVV-<br>DGVLYGRGAVDAKGPLATFVVAGARA--<br>KLPPGVRLTVVGAVEEEVMSSRGARHLIATraPDAVVIGEPSGWDGVVLGYRGSVALEYRVTVPMSHSAGPEATAAELAADFWYRLRTWCAEWS<br>V---GIDHAFHRVEPKLNALNSSSDGLYGEAVARIGLRLPPALSPEEAIAVATSLAS--<br>EGEVTATVNAPAFQTDKRQPIVAAFLAAVRAHGGTPRLKLKTGTSDMNLVGPAWGCPIVAYGPGDSRLDHTPEEHVPLADLERATAILTTAIERVA<br>AQ<br>>5xoyA_202 weight: 0.0063 score: 930.38 eval: 1e-45 prob: 100 identity: 0.4503 startpos: 1<br>=-------LDPVEFLKGALEIPSPSGKERLVAEYLAEGMQKLGLKGFVDEADNARGQVGEGPVQVVLLGHIDTVPGQIPVRLE-<br>GGRLFGRGAVDAKGPFVAMIFAAAGLSEEARKRLTVHLVGATEEEAPSSKGARFVAPRLKPHYAVIGEPSGWEGITLGYKGRLLVKARREKDH------ |

| | | | | | | |
|---|---|---|---|---|---|---|
|  |  |  |  |  |  | EPNAAEELISYFVAIKAWAEAMNVG----QRPFDQVQYTLRDFRVHP---RQVAEMFFDLRLPPRLPPEEAIRHLTAYA-<br>PPTIELEFFGREVPYQGPKDTPLTRAFRQAIRKAGGRPVFKLKTGTSDMNVLAPHWPVPMVAYGPGDSTLDHTPYEHVEVAEFLKGIEVLRGALEA<br>LAQT<br>>5xoyA_103 weight: 0.0054 score: 375.1 eval: 7e-13 prob: n/a identity: 0.4530 startpos: 1<br>=-------LDPVEFLKGALEIPSPSGKERLVAEYLAEGMQKLGLKGFVDEADNARGQVGEGPVQVVLLGHIDTVPGQIPVRLE-<br>GGRLFGRGAVDAKGPFVAMIFAAAGLSEEARKRLTVHLVGATEEEAPSSKGARFVAPRLKPHYAVIGEPSGWEGITLGYKGRLLVKARREKD-------<br>ePNAAEELISYFVAIKAWAEAMN----VGQRPFDQVQYTLRDFRVHP---RQVAEMFFDLRLPPRLPPEEAIRHLTAYA-<br>PPTIELEFFGREVPYQGPKDTPLTRAFRQAIRKAGGRPVFKLKTGTSDMNVLAPHWPVPMVAYGPGDSTLDHTPYEHVEVAEFLKGIEVLRGALEA<br>LAQT<br>>4q7aD_303 weight: 0.0041 score: 22.55 eval: n/a prob: n/a identity: 0.4420 startpos: 1<br>------SMTTADLLRGLVSIPSPSGAEAPAVEWLCQQMAALGYQAEPDGAGNAVGTRGEGPREIMLLGHIDTVPGEVPVQVV-<br>DGVLYGRGAVDAKGPLATFVVAGARA--<br>KLPPGVRLTVVGAVEEEVMSSRGARHLIATraPDAVVIGEPSGWDGVVLGYRGSVALEYRVTVPMSHSAT----AAELAADFWYRLRTWCAEWSV-<br>--GIDHAFHRVEPKLNALNSSSDGLYGEAVARIGLRLPPALSPEEAIAVATSLAS--<br>EGEVTATVNAPAFQTDKRQPIVAAFLAAVRAHGGTPRLKLKTGTSDMNLVGPAWGCPIVAYGPGDSRLDHTPEEHVPLADLERATAILTTAIER----<br>>4q7aD_104 weight: 0.0037 score: 366.7 eval: 1.4e-12 prob: n/a identity: 0.4392 startpos: 1<br>------SMTTADLLRGLVSIPSPSGAEAPAVEWLCQQMAALGYQAEPDGAGNAVGTRGEGPREIMLLGHIDTVPGEVPVQVV-<br>DGVLYGRGAVDAKGPLATFVVAGARAK--<br>LPPGVRLTVVGAVEEEVMSSRGARHLIAteAPDAVVIGEPSGWDGVVLGYRGSVALEYRVTVPMSHS----ATAAELAADFWYRLRTWCAEWS---<br>VGIDHAFHRVEPKLNALNSSSDGLYGEAVARIGLRLPPALSPEEAIAVATSLA--<br>SEGEVTATVNAPAFQTDKRQPIVAAFLAAVRAHGGTPRLKLKTGTSDMNLVGPAWGCPIVAYGPGDSRLDHTPEEHVPLADLERATAILTTAIER--<br>--<br>>5vo3A_105 weight: 0.0029 score: 343 eval: 8.9e-12 prob: n/a identity: 0.1768 startpos: 6<br>------KEKVVSLAQDLIRRPSISPNDEGCQQIIAERLEKLGFQIEWMptLNLWAKHGTSEPVIAFAGHTDVVPtpFSAEII-<br>DGMLYGRGAADMKGSLAAMIVAAEEYVKANphKGTIALLITSDEEATAKDGTIHVVETlkITYCMVGEPSSadVVKNGRRGSITGNLYIQGIQGHVA<br>YPheNPIHKAALFLQELTTYQ------WDKGNEFFPPTSLQIANIHasNNVIPAELYIQFNLRYCTEVTDEIIKQKVAEMLEklKYRIEWNLSGKPFLTK-<br>PGKLLDSITSAIEETiiTPKAETGGGTSDGRFIALM-GAEVVEFGPLNS-TIHKVNECVSVEDLGKCGEIYHKMLVNLLD-<br>>4q7aA_203 weight: 0.0025 score: 528.35 eval: 1e-45 prob: 100 identity: 0.4530 startpos: 4<br>=-SISPMSMTTADLLRGLVSIPSPSGAEAPAVEWLCQQMAALGYQAEPDGAGNAVGTRGEGPREIMLLGHIDTVPGEVPVQV-<br>VDGVLYGRGAVDAKGPLATFVVAgaKLPP----<br>GVRLTVVGAVEEEVMSSRGARHLitREAPDAVVIGEPSGWDGVVLGYRGSVALEYRVTVPMSHSAGPEATAAELAADFWYRLRTWCAEWSVG--<br>-IDHAFHRVEPKLNALNSSSDGLYGEAVARIGLRLPPALSPEEAIAVATSLASEG--<br>EVTATVNAPAFQTDKRQPIVAAFLAAVRAHGGTPRLKLKTGTSDMNLVGPAWGCPIVAYGPGDSRLDHTPEEHVPLADLERATAILTTAIERVAA<br>Q<br>>3iszA_106 weight: 0.0025 score: 341.9 eval: 9.7e-12 prob: n/a identity: 0.1713 startpos: 2<br>------KEKVVSLAQDLIRRPSISPNDEGCQQIIAERLEKLGFQIEWMptLNLWAKHGTSEPVIAFAGHTDVVPtpFSAEII-<br>DGMLYGRGAADMKGSLAAMIVAAEEYVKANphKGTIALLITSDEEATAKDGTIHVVETlkITYCMVGEPSSadVVKNGRRGSITGNLYIQGI-----<br>HLAENPIHKAALFLQELTTYQ------WDKGNEFFPPTSLQIANIHasNNVIPAELYIQFNLRYCTEVTDEIIKQKVAEMLEklKYRIEWNLSGKPFLTK-<br>PGKLLDSITSAIEETiiTPKAETGGGTSDGRFIALM-GAEVVEFGPLNS-TIHKVNECVSVEDLGKCGEIYHKMLVNLLD-<br>>5xoyA_304 weight: 0.0025 score: 22.49 eval: n/a prob: n/a identity: 0.4558 startpos: 1<br>=-------LDPVEFLKGALEIPSPSGKERLVAEYLAEGMQKLGLKGFVDEADNARGQVGEGPVQVVLLGHIDTVPGQIPVR-<br>LEGGRLFGRGAVDAKGPFVAMIFAAAGLSEEARKRLTVHLVGATEEEAPSSKGARFVAPRLKPHYAVIGEPSGWEGITLGYKGRLLVKARREK--<br>DHE----PNAAEELISYFVAIKAWAEAMNV----GQRPFDQVQYTLRDFRVHP---RQVAEMFFDLRLPPRLPPEEAIRHLTAYA-<br>PPTIELEFFGREVPYQGPKDTPLTRAFRQAIRKAGGRPVFKLKTGTSDMNVLAPHWPVPMVAYGPGDSTLDHTPYEHVEVAEFLKGIEVLRGALEA<br>LAQT<br>>4o23A_107 weight: 0.0022 score: 341 eval: 1e-11 prob: n/a identity: 0.1823 startpos: 2<br>------ETQSLELAKELISRPSVTPDDRDCQKLLAERLHKIGFAAEELhtKNIWLRRGTKAPVVCFAGHTDVVPtpFEPAER-<br>DGRLYGRGAADMKTSIACFVTACERFVAKHphQGSIALLITSDEEGDALDGTTKVVDVllIDYCIVGEPTAvdMIKNGRRGSLSGNLTVKGKQGHIAY<br>PhiNPVHTFAPALLELTQEV------WDEGNEYFPPTSFQISNING-tNVIPGELNVKFNFRFSTESTEAGLKQRVHAILDkvQYDLQWSCSGQPFLTQ-<br>AGKLTDVARAAIAETciEAELSTTGGTSDGRFIKA-IAQELIELGPSNA-TIHQINENVRLNDIPKLSAVYEGILARLLA-<br>>3gb0A_108 weight: 0.0021 score: 339.4 eval: 1.2e-11 prob: n/a identity: 0.1685 startpos: 5 |

| | | | | | | |
|---|---|---|---|---|---|---|
|  |  |  |  |  |  | ------QERLVNEFMELVQVDSETKFEAEICKVLTKKFTDLGVEVFED-aGNLICTLPagVDTIYFTSHMDTVVPgiKPSIK-<br>DGYIVSDggADDKAGLASMFEAIRVLKEKNIPHGTIEFIITVGEES-<br>GLVGAKALDrrITAKYGYALDSDgvGEIVVAAPTQAKVNAIIRGKTAHAGVagVSAITIAAKAIAKMP------------LGRIDSETTANIGRFEG-<br>tNIVCDHVQIFAEARSLINEKMEAQVAKMKEAFegGHADVEVNVMYPGFKFADGDHVVEVAKRAAEKIGRTPSLHQSGGGSDANVIAGH-<br>GIPTVNLAVGYE-EIHTTNEKIPVEELAKTAELVVAIIEEVAK-<br>>1vgyA_110 weight: 0.0021 score: 337.1 eval: 1.4e-11 prob: n/a identity: 0.1796 startpos: 2<br>------ETQSLELAKELISRPSVTPDDRDCQKLMAERLHKIGFAAEEMhtKNIWLRRGTKAPVVCFAGHTDVVPtpFEPAER-<br>DGRLYGRGAADMKTSIACFVTACERFVAKHphQGSIALLITSDEEGDALDGTTKVVDVllIDYCIVGEPTAvdMIKNGRRGSLSGNLTVKGKQGHIAY<br>PhiNPVHTFAPALLELTQEV------WDEGNEYFPPTSFQISNING-tNVIPGELNVKFNFRFSTESTEAGLKQRVHAILDkvQYDLQWSCSGQPFLTQ-<br>AGKLTDVARAAIAETciEAELSTTGGTSDGRFIKA-MAQELIELGPSNA-TIHQINENVRLNDIPKLSAVYEGILVRLL--<br>>3ic1A_109 weight: 0.0021 score: 338 eval: 1.3e-11 prob: n/a identity: 0.1713 startpos: 2<br>------KEKVVSLAQDLIRRPSISPNDEGCQQIIAERLEKLGFQIEWMptLNLWAKHGTSEPVIAFAGHTDVVPtpFSAEII-<br>DGMLYGRGAADMKGSLAAMIVAAEEYVKANphKGTIALLITSDEEATAKDGTIHVVETlkITYCMVGEPSSadVVKNGRRGSITGNLYIQGIQ----<br>hLAENPIHKAALFLQELTTYQ------WDKGNEFFPPTSLQIANIHatGSVIPAELYIQFNLRYCTEVTDEIIKQKVAEMLEklKYRIEWNLSGKPFLTK-<br>PGKLLDSITSAIEETiiTPKAETGGGTSDGRFIALM-GAEVVEFGPLNS-TIHKVNECVSVEDLGKCGEIYHKMLVNLLD-<br>>3ct9B_305 weight: 0.0018 score: 20.64 eval: n/a prob: n/a identity: 0.1823 startpos: 1<br>YDIPTMTAEAVSLLKSLISIPSISREETQAADFLQNYIEAEGMQTGRKG-NNVWCLSPmdKPTILLNSHIDTVKPvfTPREENG-<br>KLYGLGSNDAGASVVSLLQVFLQLC-RTSQNYNLIYLASCEEEVSGKEGIESVLplPPVSFAIVGEPTEM-<br>QPAIAEKGLMVLDVTATGKAGHAARDedNAIYKVLNDIAWFRDY------<br>RFEKESPLLGPVKMSVTVIntQHNVVPDKCTFVVDIRSNELYSNEDLFAEIRK---HIACDAKasFRLNSSRIDEKHPFVQKAVKM------<br>GRIPFGSPTLSDQALM----SFASVKIGPGRSSRSHTAEEYIMLKEIEEAIGIYLDLLDGLKL-<br>>3ct9A_306 weight: 0.0015 score: 20.53 eval: n/a prob: n/a identity: 0.1878 startpos: 1<br>YDIPTMTAEAVSLLKSLISIPSISREETQAADFLQNYIEAEGMQTGRKG-NNVWCLSplKKPTILLNSHIDTVKPVFTPREE-<br>NGKLYGLGSNDAGASVVSLLQVFLQLC-RTSQNYNLIYLASCEEEVSGKEGIESVLPgpPVSFAIVGEPTEM-<br>QPAIAEKGLMVLDVTATGKAGHAARDedNAIYKVLNDIAWFRDY------<br>RFEKESPLLGPVKMSVTVIntQHNVVPDKCTFVVDIRSNELYSNEDLFAEIRKHIA---CDAKasFRLNSSRIDEKHPFVQKAVKM------<br>GRIPFGSPTLSDQALM----SFASVKIGPGRSSRSHTAEEYIMLKEIEEAIGIYLDLLDGLKL-<br>>3pfoA_307 weight: 0.0014 score: 19.78 eval: n/a prob: n/a identity: 0.2238 startpos: 12<br>AAVDRNFNDQVAFLQRMVQFRSVRGEEAPQQEWLAQQFADRGYKVDTfgSMQVVATADskGRSLILQGHIDVVPEgweAKV-<br>RDGWMIGRGAQDMKGGVSAMIFALDAIRtgYAPDARVHVQTVTEEE-STGNGALSTLmgYRADACLIPEPTGH-<br>TLTRAQVGAVWFRLRVRGTPVHVAYSetSAILSAMHLIRAFEEYTKELNAqdPWFGQVKNPIKFNVGIIkdWASSTAAWCELDCRLGLLTGDTPQE<br>AMRGIEKCLAdnPAELVWSgqADPAVCEPGGVAEDVLTAAHKAAfaPLDARLSTAVNDTRYYSVDYGIPALCYGPYG-<br>QGPHAFDERIDLESLRKTTLSIALFVAEWCGL<br>>5k8pA_310 weight: 0.0013 score: 19.67 eval: n/a prob: n/a identity: 0.2127 startpos: 8<br>KTLDGMREGLIQTAVELGSIEAPTGREGAAGDYVYEWMARNGFGpvFDDRFNVVGRLrgGGASLSFNSHLDTIMArhEAWHE-<br>EGRIYGYSVVNCKGPMACWLIAAKALKegAALKGDVVLTAVCGEIDCEpiGARYAIsgAISDYALVAEATNF-<br>KPAWVEAGKVFLKVTVfgPSRYtvAAlpNAIVRMAKLVEALEEWADNYEKRYTREYGGgvVPKVAIGAIryKIYAFPELCSIYMDIRLNPDTNPLVVQR<br>EVEAVVSKLGleVKPFLFRRGYEAQGIEPLQNALEVAHREVveRPGSPECSMWRDTNPYNEL-GIPSLTYGCGG--<br>GAGGGNTYFLVDDMLKAAKVYAMTAMDLCNR<br>>2rb7A_308 weight: 0.0013 score: 19.7 eval: n/a prob: n/a identity: 0.1713 startpos: 1<br>--GMSSMQHIVELTSDLIRFPSMHSriSRCAGFIMDWCAQNGIHAERmgIPSVMVLPEKGRAGLLLMAHIDVVDAefVPR-<br>VENDRLYGRGANDDKYAVALGLVMFRDRLNasQKDMALGLLITGDEEIGGMNGAAKALPLIRADYVVALDGGNPQQVITKEKGIIDIKLTCTGKA<br>AHGARPWmnAVDLLMEDYTRLKTLF-------AEENEDHWHRTVNLGRIreSTNKVPDVAEGWFNIRVTEHDDPGALIDKIRKTVSG---<br>TVSIVRTVPVF-LAADSPYTERLLALSG-----ATAGKAHGASDARYL-GENGLTGVVWGAEGFNTLHSRDECLHIPSLQSIYDPLMQLAREMEE-<br>>5k8nA_309 weight: 0.0013 score: 19.69 eval: n/a prob: n/a identity: 0.2127 startpos: 7<br>KTLDGMREGLIQTAVELGSIEAPTGREGAAGDYVYEWMARNGFGpvFDDRFNVVGRLrgGGASLSFNSHLDTIMArhEAWHE-<br>EGRIYGYSVVNCKGPMACWLIAAKALKegAALKGDVVLTAVCGEilAEDIGARYAIsgAISDYALVAEATNF-<br>KPAWVEAGKVFLKVTVfgPSRYtvAAlpNAIVRMAKLVEALEEWADNYEKRYTREYGGgvVPKVAIGAIrvPYKIypELCSIYMDIRLNPDTNPLVVQR<br>EVEAVVSKLGleVKPFLFRRGYEAQGIEPLQNALEVAHREvtERPGSPECSMWRDTNPYNEL-GIPSLTYGCGG--<br>GAGGGNTYFLVDDMLKAAKVYAMTAMDLCNR<br>>4q7aD_204 weight: 0.0011 score: 843.4 eval: 1e-45 prob: 100 identity: 0.4392 startpos: 1 |

| | | | | | | |
|---|---|---|---|---|---|---|
|  |  |  |  |  |  | ------SMTTADLLRGLVSIPSPSGAEAPAVEWLCQQMAALGYQAEPDGAGNAVGTRGEGPREIMLLGHIDTVPGEVPVQVV-<br>DGVLYGRGAVDAKGPLATFVVAGARAK--<br>LPPGVRLTVVGAVEEEVMSSRGARHLIAteAPDAVVIGEPSGWDGVVLGYRGSVALEYRVTVPMSHSAT----AAELAADFWYRLRTWCAEWSVG-<br>--IDHAFHRVEPKLNALNSSSDGLYGEAVARIGLRLPPALSPEEAIAVATSLA--<br>SEGEVTATVNAPAFQTDKRQPIVAAFLAAVRAHGGTPRLKLKTGTSDMNLVGPAWGCPIVAYGPGDSRLDHTPEEHVPLADLERATAILTTAIER--<br>--<br>>5xoyB_205 weight: 0.0005 score: 642.86 eval: 1e-45 prob: 100 identity: 0.4641 startpos: 1<br>=-------LDPVEFLKGALEIPSPSGKERLVAEYLAEGMQKLGLKGFVDEADNARGQVGEGPVQVVLLGHIDTVPGQIPVRLE-<br>GGRLFGRGAVDAKGPFVAMIFAAAGLSEEARKRLTVHLVGATEEEAPSSKGARFVAPRLKPHYAVIGEPSGWEGITLGYKGRLLVKARREKDHFHS<br>AHHEPNAAEELISYFVAIKAWAEAMNV----GQRPFDQVQYTLRDFRVHPAELRQVAEMFFDLRLPPRLPPEEAIRHLTAYAP-<br>PTIELEFFGREVPYQGPKDTPLTRAFRQAIRKAGGRPVFKLKTGTSDMNVLAPHWPVPMVAYGPGDSTLDHTPYEHVEVAEFLKGIEVLRGALEAL<br>AQT<br>>5k8mA_206 weight: 0.0004 score: 509.66 eval: 1e-45 prob: 100 identity: 0.2099 startpos: 7<br>KTLDGMREGLIQTAVELGSIEAPTGREGAAGDYVYEWMARNGFGPErdDRFNVVGRLRGTgaSLSFNSHLDTIEA-W---HE-<br>EGRIYGYSVVNCKGPMACWLIAAKALKegAALKGDVVLTAVCGEifQGHdgARYAIsaI-SDYALVAEATNF-<br>KPAWVEAGKVFLKVTVfgPSRYtvadSPNAIVRMAKLVEALEEWADNYEK-<br>RYTREYGgvVPKVAIGAIRGGvkIypELCSIYMDIRLNPDTNPLVVQREVEAVVSKlkAEVKPFLFRRGYEAQGIEPLQNALEVAHREVVGrpGSPECS<br>MWRDTNPYNE-LGIPSLTYGCGGN--T-----YFLVDDMLKAAKVYAMTAMDLCNR |
| 4. | (WP_034341940.1) LysW-lysine Hydrolase | 352 | 5xoy, 4q7a, 5vo3, 3isz, 4o23, 3ic1, 3gb0, 1vgy, 3pfo, 3ct9, 5k8o, 5k8p, 5k8m | *Deinococcus misasensis* | 0.98 | MSIELLRQAVAIPSVSYHEADVAKYLAGYMGKHGFENAHVDVSGSAIGERGNGPLTVVLLGHIDTVPGDIQVRIEDGKLYGRGSVDAKGSFCTFVS<br>AVQQLPESALQKARFVCIGATEEEAPTSRGARHAVTKYTPNYIFIGEPSSWDAITLGYKGRLIVKAEMVKDNFHTAGEGTSASDDVVEFCTRVKKW<br>AQEHAEKHNATGAFNTIQVMVQSIHGDADGMQQVAKASVGLRLPPGMHPSAAQKEIFQLKLPGMHLQFVSHEVPYRGEKDTPISRALRLAIREV<br>GGTPKFKVKTGTSDMNVVAPHWDAPMVAYGPGDSALDHTPFEHLELAEYEKAIEVLKRALIKLVGA<br>>5xoyB_201 weight: 0.3516 score: 505.5 eval: 1e-45 prob: 100 identity: 0.4801 startpos: 2<br>DPVEFLKGALEIPSPSGKERLVAEYLAEGMQKLGLK-<br>GFVDEADNARGQVGEGPVQVVLLGHIDTVPGQIPVRLEGGRLFGRGAVDAKGPFVAMIFAAAGLSEEARKRLTVHLVGATEEE-<br>asSKGARFVAPRLKPHYAVIGEPSGWEGITLGYKGRLLVKARREKDHFHSAHHEPNAAEELISYFVAIKAWAEAMNV---<br>GQRPFDQVQYTLRDFRVHPAELRQVAEMFFDLRLPPRLPPEEAIRHLTAYAPPTIELEFFGREVPYQGPKDTPLTRAFRQAIRKAGGRPVFKLKTGTS<br>DMNVLAPHWPVPMVAYGPGDSTLDHTPYEHVEVAEFLKGIEVLRGALEALAQT<br>>5xoyB_101 weight: 0.3008 score: 384.4 eval: 2.3e-13 prob: n/a identity: 0.4773 startpos: 2<br>DPVEFLKGALEIPSPSGKERLVAEYLAEGMQKLGLKGFVDE-<br>ADNARGQVGEGPVQVVLLGHIDTVPGQIPVRLEGGRLFGRGAVDAKGPFVAMIFAAAGLSEEARKRLTVHLVGATEEEAPSSKGARFVAPRLKPH<br>YAVIGEPSGWEGITLGYKGRLLVKARREKDHFHSAHHEPNAAEELISYFVAIKAWAEAMNVGQRPF---<br>DQVQYTLRDFRVHPAELRQVAEMFFDLRLPPRLPPEEAIRHLTAYAPPTIELEFFGREVPYQGPKDTPLTRAFRQAIRKAGGRPVFKLKTGTSDMNV<br>LAPHWPVPMVAYGPGDSTLDHTPYEHVEVAEFLKGIEVLRGALEALAQT<br>>5xoyB_301 weight: 0.2861 score: 24.46 eval: n/a prob: n/a identity: 0.4858 startpos: 2<br>DPVEFLKGALEIPSPSGKERLVAEYLAEGMQKLGLK-<br>GFVDEADNARGQVGEGPVQVVLLGHIDTVPGQIPVRLEGGRLFGRGAVDAKGPFVAMIFAAAGLSEEARKRLTVHLVGATEEEAPSSKGARFVAP<br>RLKPHYAVIGEPSGWEGITLGYKGRLLVKARREKDHFHSAHHEPNAAEELISYFVAIKAWAEAMNVGQRP---<br>FDQVQYTLRDFRVHPAELRQVAEMFFDLRLPPRLPPEEAIRHLTAYAPPTIELEFFGREVPYQGPKDTPLTRAFRQAIRKAGGRPVFKLKTGTSDMN<br>VLAPHWPVPMVAYGPGDSTLDHTPYEHVEVAEFLKGIEVLRGALEALAQT<br>>5xoyA_102 weight: 0.0083 score: 374.9 eval: 4.9e-13 prob: n/a identity: 0.4716 startpos: 2<br>DPVEFLKGALEIPSPSGKERLVAEYLAEGMQKLGLKGFVDE-<br>ADNARGQVGEGPVQVVLLGHIDTVPGQIPVRLEGGRLFGRGAVDAKGPFVAMIFAAAGLSEEARKRLTVHLVGATEEEAPSSKGARFVAPRLKPH<br>YAVIGEPSGWEGITLGYKGRLLVKARREKD-------ePNAAEELISYFVAIKAWAEAMNVGQ---RPFDQVQYTLRDFRVHP---<br>RQVAEMFFDLRLPPRLPPEEAIRHLTAYAPPTIELEFFGREVPYQGPKDTPLTRAFRQAIRKAGGRPVFKLKTGTSDMNVLAPHWPVPMVAYGPG<br>DSTLDHTPYEHVEVAEFLKGIEVLRGALEALAQT<br>>5xoyA_202 weight: 0.0069 score: 911.12 eval: 1e-45 prob: 100 identity: 0.4773 startpos: 2<br>DPVEFLKGALEIPSPSGKERLVAEYLAEGMQKLGLK-<br>GFVDEADNARGQVGEGPVQVVLLGHIDTVPGQIPVRLEGGRLFGRGAVDAKGPFVAMIFAAAGLSEEARKRLTVHLVGATEEEAPSSKGARFVAP<br>RLKPHYAVIGEPSGWEGITLGYKGRLLVKARREKDH------EPNAAEELISYFVAIKAWAEAMNVG---QRPFDQVQYTLRDFRVHP---<br>RQVAEMFFDLRLPPRLPPEEAIRHLTAYAPPTIELEFFGREVPYQGPKDTPLTRAFRQAIRKAGGRPVFKLKTGTSDMNVLAPHWPVPMVAYGPG<br>DSTLDHTPYEHVEVAEFLKGIEVLRGALEALAQT |

| | | | | | | |
|---|---|---|---|---|---|---|
|  |  |  |  |  |  | >5xoyA_302 weight: 0.0067 score: 23.01 eval: n/a prob: n/a identity: 0.4773 startpos: 2<br>DPVEFLKGALEIPSPSGKERLVAEYLAEGMQKLGLK-<br>GFVDEADNARGQVGEGPVQVVLLGHIDTVPGQIPVRLEGGRLFGRGAVDAKGPFVAMIFAAAGLSEEARKRLTVHLVGATEEEAPSSKGARFVAP<br>RLKPHYAVIGEPSGWEGITLGYKGRLLVKARREK--DHE----PNAAEELISYFVAIKAWAEAMNVGQRP---FDQVQYTLRDFRVHP---<br>RQVAEMFFDLRLPPRLPPEEAIRHLTAYAPPTIELEFFGREVPYQGPKDTPLTRAFRQAIRKAGGRPVFKLKTGTSDMNVLAPHWPVPMVAYGPG<br>DSTLDHTPYEHVEVAEFLKGIEVLRGALEALAQT<br>>4q7aA_103 weight: 0.0050 score: 371.6 eval: 6.4e-13 prob: n/a identity: 0.3892 startpos: 11<br>TTADLLRGLVSIPSPSGAEAPAVEWLCQQMAALGYQAEPDG-<br>AGNAVGTRGEGPREIMLLGHIDTVPGEVPVQVVDGVLYGRGAVDAKGPLATFVVAGARAKLP--<br>PGVRLTVVGAVEEEVMSSRGARHLIAteAPDAVVIGEPSGWDGVVLGYRGSVALEYRVTVPMSHSAGPEATAAELAADFWYRLRTWCAEWSVGI<br>DHA--FHRVEPKLNALNSSSDGLYGEAVARIGLRLPPALSPEEAIAVATSLA-<br>SEGEVTATVNAPAFQTDKRQPIVAAFLAAVRAHGGTPRLKLKTGTSDMNLVGPAWGCPIVAYGPGDSRLDHTPEEHVPLADLERATAILTTAIERV<br>AAQ<br>>4q7aA_303 weight: 0.0035 score: 22.58 eval: n/a prob: n/a identity: 0.3977 startpos: 11<br>TTADLLRGLVSIPSPSGAEAPAVEWLCQQMAALGY-<br>QAEPDGAGNAVGTRGEGPREIMLLGHIDTVPGEVPVQVVDGVLYGRGAVDAKGPLATFVVAGARA--<br>KLPPGVRLTVVGAVEEEVMSSRGARHLIATraPDAVVIGEPSGWDGVVLGYRGSVALEYRVTVPMSHSAGPEATAAELAADFWYRLRTWCAEWS<br>VGID--HAFHRVEPKLNALNSSSDGLYGEAVARIGLRLPPALSPEEAIAVATSLA-<br>SEGEVTATVNAPAFQTDKRQPIVAAFLAAVRAHGGTPRLKLKTGTSDMNLVGPAWGCPIVAYGPGDSRLDHTPEEHVPLADLERATAILTTAIERV<br>AAQ<br>>4q7aD_104 weight: 0.0034 score: 361.1 eval: 1.5e-12 prob: n/a identity: 0.3892 startpos: 3<br>TTADLLRGLVSIPSPSGAEAPAVEWLCQQMAALGYQAEPDG-<br>AGNAVGTRGEGPREIMLLGHIDTVPGEVPVQVVDGVLYGRGAVDAKGPLATFVVAGARAKLP--<br>PGVRLTVVGAVEEEVMSSRGARHLIATraPDAVVIGEPSGWDGVVLGYRGSVALEYRVTVPMSHSA----TAAELAADFWYRLRTWCAEWSVGI---<br>hAFHRVEPKLNALNSSSDGLYGEAVARIGLRLPPALSPEEAIAVATSLA-<br>SEGEVTATVNAPAFQTDKRQPIVAAFLAAVRAHGGTPRLKLKTGTSDMNLVGPAWGCPIVAYGPGDSRLDHTPEEHVPLADLERATAILTTAIER--<br>--<br>>4q7aD_203 weight: 0.0028 score: 858.06 eval: 1e-45 prob: 100 identity: 0.3920 startpos: 3<br>TTADLLRGLVSIPSPSGAEAPAVEWLCQQMAALGYQA-<br>EPDGAGNAVGTRGEGPREIMLLGHIDTVPGEVPVQVVDGVLYGRGAVDAKGPLATFVVAGARAK--<br>LPPGVRLTVVGAVEEEVMSSRGARHLIAteAPDAVVIGEPSGWDGVVLGYRGSVALEYRVTVPMSHSAT----AAELAADFWYRLRTWCAEWSVG-<br>-IDHAFHRVEPKLNALNSSSDGLYGEAVARIGLRLPPALSPEEAIAVATSL-<br>ASEGEVTATVNAPAFQTDKRQPIVAAFLAAVRAHGGTPRLKLKTGTSDMNLVGPAWGCPIVAYGPGDSRLDHTPEEHVPLADLERATAILTTAIER<br>----<br>>5vo3A_105 weight: 0.0027 score: 340.3 eval: 7.8e-12 prob: n/a identity: 0.2102 startpos: 8<br>KVVSLAQDLIRRPSISPNDEGCQQIIAERLEKLGFQIEWMpdTLNLWAKHGTSEPVIAFAGHTDVVPtpFSAEIIDGMLYGRGAADMKGSLAAMIV<br>AAEEYVKAnnHKGTIALLITSDEEATAKDGTIHVVETlkITYCMVGEPSSadVVKNGRRGSITGNLYIQGIQGHVAYpaENPIHKAALFLQELTTYQ-----<br>WDKGNEFFPPTSLQIANIHA-nNVIPAELYIQFNLRYCTEVTDEIIKQKVAEMlnLKYRIEWNLSGKPFLTK-<br>PGKLLDSITSAIEETiiTPKAETGGGTSDGRFIA-LMGAEVVEFGPLNS-TIHKVNECVSVEDLGKCGEIYHKMLVNLLD-<br>>3iszA_106 weight: 0.0023 score: 337.6 eval: 9.7e-12 prob: n/a identity: 0.2045 startpos: 4<br>KVVSLAQDLIRRPSISPNDEGCQQIIAERLEKLGFQIEWMpdTLNLWAKHGTSEPVIAFAGHTDVVPtpFSAEIIDGMLYGRGAADMKGSLAAMIV<br>AAEEYVKAnnHKGTIALLITSDEEATAKDGTIHVVETlkITYCMVGEPSSadVVKNGRRGSITGNLYIQGI-----HLAENPIHKAALFLQELTTYQ-----<br>WDKGNEFFPPTSLQIANIHA-nNVIPAELYIQFNLRYCTEVTDEIIKQKVAEMlnLKYRIEWNLSGKPFLTK-<br>PGKLLDSITSAIEETiiTPKAETGGGTSDGRFIALM-GAEVVEFGPLNS-TIHKVNECVSVEDLGKCGEIYHKMLVNLLD-<br>>4q7aD_304 weight: 0.0022 score: 22.01 eval: n/a prob: n/a identity: 0.3949 startpos: 3<br>TTADLLRGLVSIPSPSGAEAPAVEWLCQQMAALGYQ-<br>AEPDGAGNAVGTRGEGPREIMLLGHIDTVPGEVPVQVVDGVLYGRGAVDAKGPLATFVVAGARAKLPPG--<br>VRLTVVGAVEEEVMSSRGARHLIATraPDAVVIGEPSGWDGVVLGYRGSVALEYRVTVPMSHSAT----AAELAADFWYRLRTWCAEWSVGID--<br>HAFHRVEPKLNALNSSSDGLYGEAVARIGLRLPPALSPEEAIAVATSL-<br>ASEGEVTATVNAPAFQTDKRQPIVAAFLAAVRAHGGTPRLKLKTGTSDMNLVGPAWGCPIVAYGPGDSRLDHTPEEHVPLADLERATAILTTAIER<br>----<br>>4o23A_107 weight: 0.0021 score: 337.2 eval: 1e-11 prob: n/a identity: 0.2074 startpos: 4 |

| | | | | | | |
|---|---|---|---|---|---|---|
|  |  |  |  |  |  | QSLELAKELISRPSVTPDDRDCQKLLAERLHKIGFAAEELHftKNIWLRRGTKAPVVCFAGHTDVVPtpFEPAERDGRLYGRGAADMKTSIACFVTAC<br>ERFVAKHphQGSIALLITSDEEGDALDGTTKVVDVllIDYCIVGEPTAvdMIKNGRRGSLSGNLTVKGKQGHIAYPhiNPVHTFAPALLELTQEVWD---<br>--EGNEYFPPTSFQISNING-tNVIPGELNVKFNFRFSTESTEAGLKQRVHAIlgVQYDLQWSCSGQPFLTQ-<br>AGKLTDVARAAIAETciEAELSTTGGTSDGRFIK--iAQELIELGPSNA-TIHQINENVRLNDIPKLSAVYEGILARLLA-<br>>3ic1A_108 weight: 0.0020 score: 335.6 eval: 1.1e-11 prob: n/a identity: 0.2045 startpos: 4<br>KVVSLAQDLIRRPSISPNDEGCQQIIAERLEKLGFQIEWMpdTLNLWAKHGTSEPVIAFAGHTDVVPtpFSAEIIDGMLYGRGAADMKGSLAAMIV<br>AAEEYVKAnnHKGTIALLITSDEEATAKDGTIHVVETlkITYCMVGEPSSadVVKNGRRGSITGNLYIQGIQ----hLAENPIHKAALFLQELTTYQ-----<br>WDKGNEFFPPTSLQIANIHAggSVIPAELYIQFNLRYCTEVTDEIIKQKVAEMlnLKYRIEWNLSGKPFLTK-<br>PGKLLDSITSAIEETiiTPKAETGGGTSDGRFIA-LMGAEVVEFGPLNS-TIHKVNECVSVEDLGKCGEIYHKMLVNLLD-<br>>3gb0A_110 weight: 0.0020 score: 330.6 eval: 1.7e-11 prob: n/a identity: 0.1818 startpos: 7<br>RLVNEFMELVQVDSETKFEAEICKVLTKKFTDLGVEVFEDDtaGNLICTLPagVDTIYFTSHMDTVVPgiKPSIKDGYIVSDggADDKAGLASMFEAIRV<br>LKEKNIPHGTIEFIITVGEES-GLVGAKALDReiTAKYGYALDSDgvGEIVVAAPTQAKVNAIIRGKTAHAGVagVSAITIAAKAIAKMPLG----------<br>RIDSETTANIGRFEG-tNIVCDHVQIFAEARSLINEKMEAQVAKMKEAF-<br>gHADVEVNVMYPGFKFADGDHVVEVAKRAAEKIGRTPSLHQSGGGSDANVIAGH-GIPTVNLAVGYE-<br>EIHTTNEKIPVEELAKTAELVVAIIEEVAK-<br>>1vgyA_109 weight: 0.0020 score: 335.4 eval: 1.2e-11 prob: n/a identity: 0.2045 startpos: 4<br>QSLELAKELISRPSVTPDDRDCQKLMAERLHKIGFAAEEMhnTKNIWLRRGTKAPVVCFAGHTDVVPtpFEPAERDGRLYGRGAADMKTSIACFVT<br>ACERFVAKhnHQGSIALLITSDEEGDALDGTTKVVDVllIDYCIVGEPTAvdMIKNGRRGSLSGNLTVKGKQGHIAYPhiNPVHTFAPALLELTQEV----<br>-WDEGNEYFPPTSFQISNING-tNVIPGELNVKFNFRFSTESTEAGLKQRVHAIlgVQYDLQWSCSGQPFLTQ-<br>AGKLTDVARAAIAETciEAELSTTGGTSDGRFIK--mAQELIELGPSNA-TIHQINENVRLNDIPKLSAVYEGILVRLL--<br>>3pfoA_305 weight: 0.0016 score: 19.87 eval: n/a prob: n/a identity: 0.2159 startpos: 20<br>DQVAFLQRMVQFRSVRGEEAPQQEWLAQQFADRGYKVDTFsgSMQVVATADskGRSLILQGHIDVVPEgyEAKVRDGWMIGRGAQDMKGGV<br>SAMIFALDAIRTAGypDARVHVQTVTEEE-STGNGALSTLMryRADACLIPEPTGH-<br>TLTRAQVGAVWFRLRVRGTPVHVAYSEtsAILSAMHLIRAFEEYTKELNAQAVRDPWfnPIKFNVGIIKgwASSTAAWCELDCRLGLLTGDTPQEAM<br>RGIEKCLanPAELVWSGfaDPAVCEPGGVAEDVLTAAHKAAfaPLDARLSTAVNDTRYYSVDYGIPALCYGPYGQG-<br>PHAFDERIDLESLRKTTLSIALFVAEWCGL<br>>3ct9A_306 weight: 0.0014 score: 19.67 eval: n/a prob: n/a identity: 0.2017 startpos: 9<br>EAVSLLKSLISIPSISREETQAADFLQNYIEAEGMQT--<br>GRKGNNVWCLsdLKKPTILLNSHIDTVKPVFTPREENGKLYGLGSNDAGASVVSLLQVFLQLCRT-<br>SQNYNLIYLASCEEEVSGKEGIESVLPgpPVSFAIVGEPTEM-QPAIAEKGLMVLDVTATGKAGHAARDedNAIYKVLNDIAWFRDY------<br>RFEksPLLGPVKMSVTVINaqHNVVPDKCTFVVDIRSNELYSNEDLFAEIRKHIACDAKAR-SFRLNSSRIDEKHPFVQKAVKM------<br>GRIPFGSPTLSDQALM----SFASVKIGPGRSSRSHTAEEYIMLKEIEEAIGIYLDLLDGLKL-<br>>3ct9B_307 weight: 0.0013 score: 19.66 eval: n/a prob: n/a identity: 0.2017 startpos: 9<br>EAVSLLKSLISIPSISREETQAADFLQNYIEAEGMQT--<br>GRKGNNVWCLSPmdKPTILLNSHIDTVKPvfTPREENGKLYGLGSNDAGASVVSLLQVFLQLCRT-<br>SQNYNLIYLASCEEEVSGKEGIESVLPGLpvSFAIVGEPTEM-QPAIAEKGLMVLDVTATGKAGHAARDedNAIYKVLNDIAWFRDY------<br>RFeeSPLLGPVKMSVTVINaqHNVVPDKCTFVVDIRSNELYSNEDLFAEIRK--HIACDAKasFRLNSSRIDEKHPFVQKAVKM------<br>GRIPFGSPTLSDQALM----SFASVKIGPGRSSRSHTAEEYIMLKEIEEAIGIYLDLLDGLKL-<br>>4q7aA_204 weight: 0.0012 score: 748.77 eval: 1e-45 prob: 100 identity: 0.3920 startpos: 11<br>TTADLLRGLVSIPSPSGAEAPAVEWLCQQMAALGYQ-AEPDGAGNAVGTRGepR-<br>EIMLLGHIDTVPGEVPVQVVDGVLYGRGAVDAKGPLATFVVAGARAKL--<br>PPGVRLTVVGAVEEEVMSSRGARHLIAteAPDAVVIGEPSGWDGVVLGYRGSVALEYRVTVPMSHSAGPEATAAELAADFWYRLRTWCAEWSV<br>GI-DH-AFHRVEPKLNALNSSSDGLYGEAVARIGLRLPPALSPEEAIAVATSL-<br>ASEGEVTATVNAPAFQTDKRQPIVAAFLAAVRAHGGTPRLKLKTGTSDMNLVGPAWGCPIVAYGPGDSRLDHTPEEHVPLADLERATAILTTAIER<br>VAAQ<br>>5k8oA_308 weight: 0.0012 score: 19.25 eval: n/a prob: n/a identity: 0.1875 startpos: 17<br>GLIQTAVELGSIEAPTGREGAAGDYVYEWMARNGFGPERVgdRFNVVGRLrgGGASLSFNSHLDTIMArhEAWHEEGRIYGYSVVNCKGPMACW<br>LIAAKALKEAgaLKGDVVLTAVCGEIDCEpiGARYAIShaISDYALVAEATNF-<br>KPAWVEAGKVFLKVTVfgPSRYtvAAlpNAIVRMAKLVEALEEWADNYEKRYTREYGgvVPKVAIGAIRGGvyRFPELCSIYMDIRLNPDTNPLVVQR<br>EVEAVvgLKAEVKPFLFRRGYEAQGIEPLQNALEVAHREvtERPGSPECSMWRDTNPYNEL-GIPSLTYGCGGG--<br>AGGGNTYFLVDDMLKAAKVYAMTAMDLCNR<br>>5k8pA_310 weight: 0.0012 score: 19.18 eval: n/a prob: n/a identity: 0.1875 startpos: 16 |

| | | | | | | |
|---|---|---|---|---|---|---|
|  |  |  |  |  |  | GLIQTAVELGSIEAPTGREGAAGDYVYEWMARNGFgrVGVfdRFNVVGRLrgGGASLSFNSHLDTIMArhEAWHEEGRIYGYSVVNCKGPMACWL<br>IAAKALKEAgaLKGDVVLTAVCGEIDCEpiGARYAIShaISDYALVAEATNF-<br>KPAWVEAGKVFLKVTVfgPSRYtvAAlpNAIVRMAKLVEALEEWADNYEKRYTREYGgvVPKVAIGAIRGGvyAFPELCSIYMDIRLNPDTNPLVVQR<br>EVEAVvgLKAEVKPFLFRRGYEAQGIEPLQNALEVAHREVveRPGSPECSMWRDTNPYNEL-GIPSLTYGCGG--<br>GAGGGNTYFLVDDMLKAAKVYAMTAMDLCNR<br>>5k8pC_309 weight: 0.0012 score: 19.23 eval: n/a prob: n/a identity: 0.1903 startpos: 17<br>GLIQTAVELGSIEAPTGREGAAGDYVYEWMARNGFeRVGVfdRFNVVGRLrgGGASLSFNSHLDTIMArhEAWHEEGRIYGYSVVNCKGPMACW<br>LIAAKALKEAgaLKGDVVLTAVCGEIDCEpiGARYAIShaISDYALVAEATNF-<br>KPAWVEAGKVFLKVTVfgPSRYtvAAlpNAIVRMAKLVEALEEWADNYEKRYTREYGgvVPKVAIGAIRGGvyAFPELCSIYMDIRLNPDTNPLVVQR<br>EVEAVvgLKAEVKPFLFRRGYEAQGIEPLQNALEVAHREVveRPGSPECSMWRDTNPYNEL-GIPSLTYGCGG--<br>GAGGGNTYFLVDDMLKAAKVYAMTAMDLCNR<br>>5k8mA_205 weight: 0.0006 score: 517.9 eval: 1e-45 prob: 100 identity: 0.1903 startpos: 15<br>GLIQTAVELGSIEAPTGREGAAGDYVYEWMARNGFG-PErdDRFNVVGRLRGTgaSLSFNSHLDTIEA-W--HEE-<br>GRIYGYSVVNCKGPMACWLIAAKALKegAALKGDVVLTAVCGEifQGHdgARYAISHgiSDYALVAEATNF-<br>KPAWVEAGKVFLKVTVfgPSRYtvadSPNAIVRMAKLVEALEEWADNYEKRYTREYGgvVPKVAIGAIRGGvkIypELCSIYMDIRLNPDTNPLVVQRE<br>VEAVVslKAEVKPFLFRRGYEAQGIEPLQNALEVAHREVVGrpGSPECSMWRDTNPYNE-LGIPSLTYGCGGN--T-----<br>YFLVDDMLKAAKVYAMTAMDLCNR |
| 5. | (NP_295143.2) Acetyl glutamate or acetyl aminoadipate kinase | 267 | 3wwm, 3wwn, 4usj, 3u6u, 2buf, 2ap9, 2v5h, 2jj4, 2bty, 3l86, 2rd5 | *Deinococcus radiodurans* | 1 | MIVVKVGGSDGIDYDAVCADLAERWQAGEKLILVHGGSGETNRVAEALGHPPKFVTSPSGYTSRFTDRQTLEIFEMVYCGKMNKGLVERLQRLGV<br>NAVGLSGLDGRIFEGKHKDSVRSVENGKVKVLRGDHTGTVEKVNTGLIELLLGAGYLPVLTPPAASYEGVAINVDGDRAAAQLAAALRAEALLLLSN<br>VPGLLRDYPDEASLIREIPANDVESYLEFAQDRMKKKVLGAAEAVAGGVGRVVFGDARAGKPISAALAGEGTVVS<br>>3wwmA_201 weight: 0.3513 score: 409.36 eval: 1e-45 prob: 100 identity: 0.6854 startpos: 1<br>MIVVKVGGAEGINYEAVAKDAASLWKEGVKLLLVHGGSAETNKVAEALGHPPRFLTHPGGQVSRLTDRKTLEIFEMVYCGLVNKRLVELLQKEGA<br>NAIGLSGLDGRLFVGRRKTAVKYVENGKVKVHRGDYTGTVEEVNKALLDLLLQAGYLPVLTPPALSYENEAINTDGDQIAALLATLYGAEALVYLSN<br>VPGLLARYPDEASLVREIPVERIeeYLALAQGRMKRKVMGAVEAVRGGVKRVVFADARVENPIRRALSGEGTVVR<br>>3wwmA_101 weight: 0.3006 score: 295.5 eval: 4.3e-11 prob: n/a identity: 0.6854 startpos: 1<br>MIVVKVGGAEGINYEAVAKDAASLWKEGVKLLLVHGGSAETNKVAEALGHPPRFLTHPGGQVSRLTDRKTLEIFEMVYCGLVNKRLVELLQKEGA<br>NAIGLSGLDGRLFVGRRKTAVKYVENGKVKVHRGDYTGTVEEVNKALLDLLLQAGYLPVLTPPALSYENEAINTDGDQIAALLATLYGAEALVYLSN<br>VPGLLARYPDEASLVREIPVERIEdyLALAQGRMKRKVMGAVEAVRGGVKRVVFADARVENPIRRALSGEGTVVR<br>>3wwmA_301 weight: 0.2859 score: 31.54 eval: n/a prob: n/a identity: 0.6816 startpos: 1<br>MIVVKVGGAEGINYEAVAKDAASLWKEGVKLLLVHGGSAETNKVAEALGHPPRFLTHPGGQVSRLTDRKTLEIFEMVYCGLVNKRLVELLQKEGA<br>NAIGLSGLDGRLFVGRRKTAVKYVENGKVKVHRGDYTGTVEEVNKALLDLLLQAGYLPVLTPPALSYENEAINTDGDQIAALLATLYGAEALVYLSN<br>VPGLLARYPDEASLVREIPVERipEYLALAQGRMKRKVMGAVEAVRGGVKRVVFADARVENPIRRALSGEGTVVR<br>>3wwnA_102 weight: 0.0083 score: 291.2 eval: 6.3e-11 prob: n/a identity: 0.6704 startpos: 1<br>MIVVKVGGAEGINYEAVAKDAASLWKEGVKLLLVHGGSAETNKVAEALGHPPRFLTHPGGQVSRLTDRKTLEIFEMVYCGLVNKRLVELLQKEGA<br>NAIGLSGLDGRLFVGRRKTAVKYVENGKVKVHRGDYTGTVEEVNKALLDLLLQAGYLPVLTPPALSYENEAINTDGDQIAALLATLYGAEALVYLSN<br>VPGLLARYPDEASLVREIPVERIEDPE----GRMKRKVMGAVEAVRGGVKRVVFADARVENPIRRALSGEGTVVR<br>>3wwnA_202 weight: 0.0069 score: 460.35 eval: 1e-45 prob: 100 identity: 0.6517 startpos: 1<br>MIVVKVGGAEGINYEAVAKDAASLWKEGVKLLLVHGGSAETNKVAEALGHPPRFLTHPGGQVSRLTDRKTLEIFEMVYCGLVNKRLVELLQKEGA<br>NAIGLSGLDGRLFVGRRKTAVKYV--<br>ENgkVHRGDYTGTVEEVNKALLDLLLQAGYLPVLTPPALSYENEAINTDGDQIAALLATLYGAEALVYLSNVPGLLARYPDEASLVREIPVERIED----<br>PEGRMKRKVMGAVEAVRGGVKRVVFADARVENPIRRALSGEGTVVR<br>>3wwnA_302 weight: 0.0067 score: 30.71 eval: n/a prob: n/a identity: 0.6704 startpos: 1<br>MIVVKVGGAEGINYEAVAKDAASLWKEGVKLLLVHGGSAETNKVAEALGHPPRFLTHPGGQVSRLTDRKTLEIFEMVYCGLVNKRLVELLQKEGA<br>NAIGLSGLDGRLFVGRRKTAVKYVENGKVKVHRGDYTGTVEEVNKALLDLLLQAGYLPVLTPPALSYENEAINTDGDQIAALLATLYGAEALVYLSN<br>VPGLLARYPDEASLVREIPVERIED----PEGRMKRKVMGAVEAVRGGVKRVVFADARVENPIRRALSGEGTVVR<br>>3u6uA_103 weight: 0.0050 score: 277.8 eval: 2e-10 prob: n/a identity: 0.6330 startpos: 1<br>MIVVKVGGAEGINYEAVAKDAASLWKEGVKLLLVHGGSAETNKVAEALGHPPRFLTHP---<br>VSRLTDRKTLEIFEMVYCGLVNKRLVELLQKEGANAIGLSGLDGRLFVGRRK---------------<br>DYTGTVEEVNKALLDLLLQAGYLPVLTPPALSYENEAINTDGDQIAALLATLYGAEALVYLSNVPGLLA--<br>PDEASLVREIPVERIEdyLALAQGRMKRKVMGAVEAVKGGVKRVVFADGRVENPIRRALSGEGTVVR<br>>3u6uA_303 weight: 0.0035 score: 30.08 eval: n/a prob: n/a identity: 0.6255 startpos: 1 |

| | | | | | | |
|---|---|---|---|---|---|---|
|  |  |  |  |  |  | MIVVKVGGAEGINYEAVAKDAASLWKEGVKLLLVHGGSAETNKVAEALGHPPRFLT---<br>HPVSRLTDRKTLEIFEMVYCGLVNKRLVELLQKEGANAIGLSGLDGRLFVGRRK---------------<br>DYTGTVEEVNKALLDLLLQAGYLPVLTPPALSYENEAINTDGDQIAALLATLYGAEALVYLSNVPGLLAP--<br>DEASLVREIPVERIEdyLALAQGRMKRKVMGAVEAVKGGVKRVVFADGRVENPIRRALSGEGTVVR<br>>4usjA_104 weight: 0.0034 score: 267.9 eval: 4.7e-10 prob: n/a identity: 0.3408 startpos: 21<br>TIVVKYGGAAMtlKSSVVSDLVLLACVGLRPILVHGGGPDINRYLKQLNIPAEFRD-----<br>GLRVTDATTMEIVSMVLVGKVNKNLVSLINAAGATAVGLSGHDGRLLTARPVPNSA----------<br>QLGFVGEVARVDPSVLRPLVDYGYIPVIASVAADDSGQAYNINADTVAGELAAALGAEKLILLTDVAGILENKEDPSSLIKEIDIKGVKKMigKVAGG<br>MIPKVKCCIRSLAQGVKTASIIDGRRQHSLLHEImgAGTMIT<br>>3u6uA_203 weight: 0.0028 score: 543.65 eval: 1e-45 prob: 100 identity: 0.6292 startpos: 1<br>MIVVKVGGAEGINYEAVAKDAASLWKEGVKLLLVHGGSAETNKVAEALGHPPRFLTH-PV--<br>SRLTDRKTLEIFEMVYCGLVNKRLVELLQKEGANAIGLSGLDGRLFVGRRK---------------<br>DYTGTVEEVNKALLDLLLQAGYLPVLTPPALSYENEAINTDGDQIAALLATLYGAEALVYLSNVPGLLA--<br>PDEASLVREIPVERIeeYLALAQGRMKRKVMGAVEAVKGGVKRVVFADGRVENPIRRALSGEGTVVR<br>>2bufA_105 weight: 0.0027 score: 266.3 eval: 5.4e-10 prob: n/a identity: 0.2772 startpos: 28<br>TLVIKYGGNAMelKAGFARDVVLMKAVGINPVVVHGGGPQIGDLLKRLSIESHFID-----<br>GMRVTDAATMDVVEMVLGGQVNKDIVNLINRHGGSAIGLTGKDAELIRAKKLTVTRQ-------<br>IIDIGHVGEVTGVNVGLLNMLVKGDFIPVIAPIGVGSNGESYNINADLVAGKVAEALKAEKLMLLTNIAGLMDKQ---<br>GQVLTGLSTEQVNELigTIYGGMLPKIRCALEAVQGGVTSAHIIDGRVPNAVLLEisGVGTLIS<br>>2ap9A_106 weight: 0.0023 score: 265.8 eval: 5.7e-10 prob: n/a identity: 0.3109 startpos: 27<br>VVVVKYGGNAMtlRRAFAADMAFLRNCGIHPVVVHGGGPQITAMLRRLGIEGDFKG-----<br>GFRVTTPEVLDVARMVLFGQVGRELVNLINAHGPYAVGITGEDAQLFTAVRRSVTVDGV-----<br>ATDIGLVGDVDQVNTAAMLDLVAAGRIPVVSTLAPDADGVVHNINADTAAAAVAEALGAEKLLMLTDIDGLYTRWPDRDSLVSEIDTGTLAQLLP<br>TLELGMVPKVEACLRAVIGGVPSAHIIDGRVTHCVLVElaGTGTKVV<br>>2rd5A_304 weight: 0.0022 score: 28.76 eval: n/a prob: n/a identity: 0.3446 startpos: 21<br>TIVVKYGGAAMTskSSVVSDLVLLACVGLRPILVHGGGPDINRYLKQLNIPAEFRDG-----<br>LRVTDATTMEIVSMVLVGKVNKNLVSLINAAGATAVGLSGHDGRLLTARPVPNS----------<br>AQLGFVGEVARVDPSVLRPLVDYGYIPVIASVAADDSGQAYNINADTVAGELAAALGAEKLILLTDVAGILENKEDPSSLIKEIDIKGVKKMIekVAGG<br>MIPKVKCCIRSLAQGVKTASIIDGRRQHSLLHEImgAGTMIT<br>>2rd5A_107 weight: 0.0021 score: 264.9 eval: 6.1e-10 prob: n/a identity: 0.3408 startpos: 21<br>TIVVKYGGAAMTskSSVVSDLVLLACVGLRPILVHGGGPDINRYLKQLNIPAEFRD-----<br>GLRVTDATTMEIVSMVLVGKVNKNLVSLINAAGATAVGLSGHDGRLLTARPVPNSA----------<br>QLGFVGEVARVDPSVLRPLVDYGYIPVIASVAADDSGQAYNINADTVAGELAAALGAEKLILLTDVAGILENKEDPSSLIKEIDIKGVKKMigKVAGG<br>MIPKVKCCIRSLAQGVKTASIIDGRRQHSLLHEImgAGTMIT<br>>2jj4A_108 weight: 0.0020 score: 263.4 eval: 7e-10 prob: n/a identity: 0.3184 startpos: 24<br>TVVVKYGGAAMklKEAVMRDIVFLACVGMRPVVVHGGGPEINAWLGRVGIEPQFHN-----<br>GLRVTDADTMEVVEMVLVGRVNKDIVSRINTTGGRAVGFCGTDGRLVLARPHD-----------<br>QEGIGFVGEVNSVNSEVIEPLLERGYIPVISSVAADENGQSFNINADTVAGEIAAALNAEKLILLTDTRGILEDPKRPESLIPRLNIPQSRELigIVGGGMI<br>PKVDCCIRSLAQGVRAAHIIDGRIPHALLLEiaGIGTMIV<br>>2btyA_109 weight: 0.0020 score: 254.9 eval: 1.5e-09 prob: n/a identity: 0.3184 startpos: 23<br>TFVIKFGGSAMkaKKAFIQDIILLKYTGIKPIIVHGGGPAISQMMKDLGIEPVFK------<br>GHRVTDEKTMEIVEMVLVGKINKEIVMNLNLHGGRAVGICGKDSKLIVAEKET----------<br>KHGDIGYVGKVKKVNPEILHALIENDYIPVIAPVGIGEDGHSYNINADTAAAEIAKSLMAEKLILLTDVDGVLKD----<br>GKLISTLTPDEAEELigTVTGGMIPKVECAVSAVRGGVGAVHIINGGLEHAILLEIfgIGTMIK<br>>2v5hA_110 weight: 0.0019 score: 249.9 eval: 2.2e-09 prob: n/a identity: 0.3184 startpos: 24<br>TVVVKYGGAAMklKEAVMRDIVFLACVGMRPVVVHGGGPEINAWLGRVGIEPQFHN-----<br>GLRVTDADTMEVVEMVLVGRVNKDIVSRINTTGGRAVGFCGTDGRLVLARPHD-----------<br>QEGIGFVGEVNSVNSEVIEPLLERGYIPVISSVAADENGQSFNINADTVAGEIAAALNAEKLILLTDTRGILEDPKRPESLIPRLNIPQSRELigIVGGGMI<br>PKVDCCIRSLAQGVRAAHIIDGRIPHALLLEiaGIGTMIV<br>>4usjA_305 weight: 0.0016 score: 28.65 eval: n/a prob: n/a identity: 0.3446 startpos: 21<br>TIVVKYGGAAMTskSSVVSDLVLLACVGLRPILVHGGGPDINRYLKQLNIPAEFRDG-----<br>LRVTDATTMEIVSMVLVGKVNKNLVSLINAAGATAVGLSGHDGRLLTARPVPNS---------- |

| | | | | | | |
|---|---|---|---|---|---|---|
|  |  |  |  |  |  | AQLGFVGEVARVDPSVLRPLVDYGYIPVIASVAADDSGQAYNINADTVAGELAAALGAEKLILLTDVAGILENKEDPSSLIKEIDIKGVKKMIekVAGG<br>MIPKVKCCIRSLAQGVKTASIIDGRRQHSLLHEieGAGTMIT<br>>2btyA_306 weight: 0.0014 score: 28.05 eval: n/a prob: n/a identity: 0.3109 startpos: 23<br>TFVIKFGGSAMKqkKAFIQDIILLKYTGIKPIIVHGGGPAISQMMKDLGIEPVFKG------<br>HRVTDEKTMEIVEMVLVGKINKEIVMNLNLHGGRAVGICGKDSKLIVAEKE------TKHG----<br>DIGYVGKVKKVNPEILHALIENDYIPVIAPVGIGEDGHSYNINADTAAAEIAKSLMAEKLILLTDVDGVL----<br>KDGKLISTLTPDEAEELigTVTGGMIPKVECAVSAVRGGVGAVHIINGGLEHAILLEIfgIGTMIK<br>>2bufA_307 weight: 0.0013 score: 27.94 eval: n/a prob: n/a identity: 0.2734 startpos: 28<br>TLVIKYGGNAMEskAGFARDVVLMKAVGINPVVVHGGGPQIGDLLKRLSIESHFIDG-----<br>MRVTDAATMDVVEMVLGGQVNKDIVNLINRHGGSAIGLTGKDAELIRAKKLTV----TRQII---<br>DIGHVGEVTGVNVGLLNMLVKGDFIPVIAPIGVGSNGESYNINADLVAGKVAEALKAEKLMLLTNIAGLMDK---<br>QGQVLTGLSTEQVNELIatIYGGMLPKIRCALEAVQGGVTSAHIIDGRVPNAVLLEisGVGTLIS<br>>3l86A_310 weight: 0.0012 score: 27.41 eval: n/a prob: n/a identity: 0.2472 startpos: 4<br>IIVIKIGGVASQQlgDFLSQIKNWQDAGKQLVIVHGGGFAINKLMEENQVPVKKING-----<br>LRVTSKDDMVLVSHALLDLVGKNLQEKLRQAGVSCQQLKSDIKHVVAADYLDK-----------<br>DTYGYVGDVTHINKRVIEEFLENRQIPILASLGYSKEGDMLNINADYLATAVAVALAADKLILMTNVKGVL----<br>ENGAVLEKITSHQVQEKIdvITAGMIPKIESAAKTVAAGVGQVLIGDNLLT----------GTLIT<br>>3l86A_204 weight: 0.0012 score: 567 eval: 1e-45 prob: 100 identity: 0.2509 startpos: 4<br>IIVIKIGGVASQQlgDFLSQIKNWQDAGKQLVIVHGGGFAINKLMEENQVPVKKIN---G--<br>LRVTSKDDMVLVSHALLDLVGKNLQEKLRQAGVSCQQLKSDIKHVVAADYLD-----------<br>KDTYGYVGDVTHINKRVIEEFLENRQIPILASLGYSKEGDMLNINADYLATAVAVALAADKLILMTNVKGVLENG----<br>AVLEKITSHQVQEktAVITAGMIPKIESAAKTVAAGVGQVLIGDNLLT----------GTLIT<br>>2ap9A_309 weight: 0.0012 score: 27.68 eval: n/a prob: n/a identity: 0.3146 startpos: 27<br>VVVVKYGGNAMTdrRAFAADMAFLRNCGIHPVVVHGGGPQITAMLRRLGIEGDFKGG-----<br>FRVTTPEVLDVARMVLFGQVGRELVNLINAHGPYAVGITGEDAQLFTAVRRSV----TVDGVAT-<br>DIGLVGDVDQVNTAAMLDLVAAGRIPVVSTLAPDADGVVHNINADTAAAAVAEALGAEKLLMLTDIDGLYTRWPDRDSLVSEIDTGTLAQLLPTL<br>ELGMVPKVEACLRAVIGGVPSAHIIDGRVTHCVLVELfgTGTKVV<br>>2jj4A_308 weight: 0.0012 score: 27.92 eval: n/a prob: n/a identity: 0.3146 startpos: 24<br>TVVVKYGGAAMKqkEAVMRDIVFLACVGMRPVVVHGGGPEINAWLGRVGIEPQFHNG-----<br>LRVTDADTMEVVEMVLVGRVNKDIVSRINTTGGRAVGFCGTDGRLVLARPHDQ-----------<br>EGIGFVGEVNSVNSEVIEPLLERGYIPVISSVAADENGQSFNINADTVAGEIAAALNAEKLILLTDTRGILEDPKRPESLIPRLNIPQSRELIaiVGGGMIP<br>KVDCCIRSLAQGVRAAHIIDGRIPHALLLEIfgIGTMIV<br>>2v5hA_205 weight: 0.0006 score: 477.82 eval: 1e-45 prob: 100 identity: 0.3184 startpos: 24<br>TVVVKYGGAAMKQeeAVMRDIVFLACVGMRPVVVHGGGPEINAWLGRVGIEPQFHN---G--<br>LRVTDADTMEVVEMVLVGRVNKDIVSRINTTGGRAVGFCGTDGRLVLARPHD-----------<br>QEGIGFVGEVNSVNSEVIEPLLERGYIPVISSVAADENGQSFNINADTVAGEIAAALNAEKLILLTDTRGILEDPKRPESLIPRLNIPQSRElqGIVGGGM<br>IPKVDCCIRSLAQGVRAAHIIDGRIPHALLLEIfgIGTMIV<br>>2btyA_206 weight: 0.0004 score: 569.09 eval: 1e-45 prob: 100 identity: 0.3146 startpos: 23<br>TFVIKFGGSAMKQekAFIQDIILLKYTGIKPIIVHGGGPAISQMMKDLGIEPVFK-G-----<br>HRVTDEKTMEIVEMVLVGKINKEIVMNLNLHGGRAVGICGKDSKLIVAEKETK----------<br>HGDIGYVGKVKKVNPEILHALIENDYIPVIAPVGIGEDGHSYNINADTAAAEIAKSLMAEKLILLTDVDGVLKD----<br>GKLISTLTPDEAEEIdGTVTGGMIPKVECAVSAVRGGVGAVHIINGGLEHAILLEikGIGTMIK<br>>2rd5A_207 weight: 0.0003 score: 447.54 eval: 1e-45 prob: 100 identity: 0.3446 startpos: 21<br>TIVVKYGGAAMTSPesVVSDLVLLACVGLRPILVHGGGPDINRYLKQLNIPAEFRD---G--<br>LRVTDATTMEIVSMVLVGKVNKNLVSLINAAGATAVGLSGHDGRLLTARPVPN----------<br>SAQLGFVGEVARVDPSVLRPLVDYGYIPVIASVAADDSGQAYNINADTVAGELAAALGAEKLILLTDVAGILENKEDPSSLIKEIDIKGVKKMigKVAG<br>GMIPKVKCCIRSLAQGVKTASIIDGRRQHSLLHemsgAGTMIT |
| 6. | (WP_051363596.1)<br>LysW-<br>Aminoadipatekinase | 289 | 3wwm, 3wwn,<br>4usj, 3u6u, 2rd5,<br>2buf, 2ap9, 2v5h,<br>2jj4, 2bty, 3l86,<br>2r8v | *Deinococcus murrayi* | 0.9 | MTPFLATTSPLVIKIGGAQNIDPRALGHDLAAEWHRGRPLVALHGGSAETDALAAALGHPPRFLTSPSGHVSRHTDRRTLEIFAMATARVNRLLVE<br>TLQGLGVNALGLCGADGQLLRARRKAAQRHVEEGGRVRLVRDDWTGTVTGASGDLLRLLLEAGYLPVVAPLALGEGGELLNVDGDRAAAAVAG<br>ALGAGALLLLSNVPGLLRAYPDEASLVGHLPAERLDEAGRWAQGRMKRKVLAAGEALAAGVPRVVIGDARRERPVRDALAGRGTVLGQPLTPQP<br>SPEVPA<br>>3wwmA_201 weight: 0.3363 score: 407.34 eval: 1e-45 prob: 100 identity: 0.5156 startpos: 1 |

| | | | | | | |
|---|---|---|---|---|---|---|
|  |  |  |  |  |  | ---------<br>MIVVKVGGAEGINYEAVAKDAASLWKEGVKLLLVHGGSAETNKVAEALGHPPRFLTHPGGQVSRLTDRKTLEIFEMVYclVNKRLVELLQKEGANA<br>IGLSGLDGRLFVGRRKTAVKYV-<br>ENGKVKVHRGDYTGTVEEVNKALLDLLLQAGYLPVLTPPALSYENEAINTDGDQIAALLATLYGAEALVYLSNVPGLLARYPDEASLVREIPVERIeeY<br>LALAQGRMKRKVMGAVEAVRGGVKRVVFADARVENPIRRALSGEGTVVR-------------<br>>3wwmA_101 weight: 0.3026 score: 286.3 eval: 1.9e-09 prob: n/a identity: 0.5052 startpos: 1<br>---------<br>MIVVKVGGAEGINYEAVAKDAASLWKEGVKLLLVHGGSAETNKVAEALGHPPRFLTHPGGQVSRLTDRKTLEIFEMVYclVNKRLVELLQKEGANA<br>IGLSGLDGRLFVGRRK-<br>TAVKYVENGKVKVHRGDYTGTVEEVNKALLDLLLQAGYLPVLTPPALSYENEAINTDGDQIAALLATLYGAEALVYLSNVPGLLARYPDEASLVREIP<br>VERIEdyLALAQGRMKRKVMGAVEAVRGGVKRVVFADARVENPIRRALSGEGTVVR-------------<br>>3wwmA_301 weight: 0.2957 score: 27.4 eval: n/a prob: n/a identity: 0.5156 startpos: 1<br>---------<br>MIVVKVGGAEGINYEAVAKDAASLWKEGVKLLLVHGGSAETNKVAEALGHPPRFLTHPGGQVSRLTDRKTLEIFEMVYclVNKRLVELLQKEGANA<br>IGLSGLDGRLFVGRRKTAVKYVENG-<br>KVKVHRGDYTGTVEEVNKALLDLLLQAGYLPVLTPPALSYENEAINTDGDQIAALLATLYGAEALVYLSNVPGLLARYPDEASLVREIPVERipEYLALA<br>QGRMKRKVMGAVEAVRGGVKRVVFADARVENPIRRALSGEGTVVR-------------<br>>3wwnA_102 weight: 0.0087 score: 280.3 eval: 3e-09 prob: n/a identity: 0.4983 startpos: 1<br>---------<br>MIVVKVGGAEGINYEAVAKDAASLWKEGVKLLLVHGGSAETNKVAEALGHPPRFLTHPGGQVSRLTDRKTLEIFEMVYclVNKRLVELLQKEGANA<br>IGLSGLDGRLFVGRRKT-<br>AVKYVENGKVKVHRGDYTGTVEEVNKALLDLLLQAGYLPVLTPPALSYENEAINTDGDQIAALLATLYGAEALVYLSNVPGLLARYPDEASLVREIPV<br>ERIEDPE----GRMKRKVMGAVEAVRGGVKRVVFADARVENPIRRALSGEGTVVR-------------<br>>4usjA_302 weight: 0.0074 score: 26.89 eval: n/a prob: n/a identity: 0.2768 startpos: 12<br>PFIQKFRGKTIVVKYGGAAMTskSSVVSDLVLLACVGLRPILVHGGGPDINRYLKQLNIPAEFRDG-----<br>LRVTDATTMEIVSMVLVgvNKNLVSLINAAGATAVGLSGHDGRLLTARPVPNS-----------<br>AQLGFVGEVARVDPSVLRPLVDYGYIPVIASVAADDSGQAYNINADTVAGELAAALGAEKLILLTDVAGILENKEDPSSLIKEIDIKGVKKMigKVAGG<br>MIPKVKCCIRSLAQGVKTASIIDGRRQHSLLHEieGAGTMITG------------<br>>3wwnA_202 weight: 0.0063 score: 461.16 eval: 1e-45 prob: 100 identity: 0.5052 startpos: 1<br>---------<br>MIVVKVGGAEGINYEAVAKDAASLWKEGVKLLLVHGGSAETNKVAEALGHPPRFLTHPGGQVSRLTDRKTLEIFEMVYclVNKRLVELLQKEGANA<br>IGLSGLDGRLFVGRRKTAVKYV-<br>ENGKVKVHRGDYTGTVEEVNKALLDLLLQAGYLPVLTPPALSYENEAINTDGDQIAALLATLYGAEALVYLSNVPGLLARYPDEASLVREIPVERIED--<br>--PEGRMKRKVMGAVEAVRGGVKRVVFADARVENPIRRALSGEGTVVR-------------<br>>3u6uA_103 weight: 0.0054 score: 271.5 eval: 5.8e-09 prob: n/a identity: 0.4810 startpos: 1<br>---------MIVVKVGGAEGINYEAVAKDAASLWKEGVKLLLVHGGSAETNKVAEALGHPPRFLTHP---<br>VSRLTDRKTLEIFEMVYclVNKRLVELLQKEGANAIGLSGLDGRLFVGRRK-----------------<br>DYTGTVEEVNKALLDLLLQAGYLPVLTPPALSYENEAINTDGDQIAALLATLYGAEALVYLSNVPGLLA--<br>PDEASLVREIPVERIEdyLALAQGRMKRKVMGAVEAVKGGVKRVVFADGRVENPIRRALSGEGTVVR-------------<br>>2rd5A_303 weight: 0.0041 score: 26.88 eval: n/a prob: n/a identity: 0.2768 startpos: 12<br>PFIQKFRGKTIVVKYGGAAMTskSSVVSDLVLLACVGLRPILVHGGGPDINRYLKQLNIPAEFRDG-----<br>LRVTDATTMEIVSMVLVgvNKNLVSLINAAGATAVGLSGHDGRLLTARPVPNS-----------<br>AQLGFVGEVARVDPSVLRPLVDYGYIPVIASVAADDSGQAYNINADTVAGELAAALGAEKLILLTDVAGILENKEDPSSLIKEIDIKGVKKMigKVAGG<br>MIPKVKCCIRSLAQGVKTASIIDGRRQHSLLHEImgAGTMITG------------<br>>4usjA_104 weight: 0.0037 score: 267.8 eval: 7.7e-09 prob: n/a identity: 0.2768 startpos: 20<br>--------KTIVVKYGGAAMTlKSSVVSDLVLLACVGLRPILVHGGGPDINRYLKQLNIPAEFRD-----<br>GLRVTDATTMEIVSMVLvkVNKNLVSLINAAGATAVGLSGHDGRLLTARPVPNSA-----------<br>QLGFVGEVARVDPSVLRPLVDYGYIPVIASVAADDSGQAYNINADTVAGELAAALGAEKLILLTDVAGILENKEDPSSLIKEIDIKGVKKMigKVAGG<br>MIPKVKCCIRSLAQGVKTASIIDGRRQHSLLHEImgAGTMITG------------<br>>2bufA_105 weight: 0.0029 score: 267.7 eval: 7.7e-09 prob: n/a identity: 0.2734 startpos: 21<br>--IRRFVGKTLVIKYGGNAMelKAGFARDVVLMKAVGINPVVVHGGGPQIGDLLKRLSIESHFID-----<br>GMRVTDAATMDVVEMVLgqVNKDIVNLINRHGGSAIGLTGKDAELIRAKKLTVTRQ-------- |

| | | | | | | |
|---|---|---|---|---|---|---|
|  |  |  |  |  |  | IIDIGHVGEVTGVNVGLLNMLVKGDFIPVIAPIGVGSNGESYNINADLVAGKVAEALKAEKLMLLTNIAGLMDKQ---<br>GQVLTGLSTEQVNELigTIYGGMLPKIRCALEAVQGGVTSAHIIDGRVPNAVLLEisGVGTLISNRKRH--------<br>>2ap9A_203 weight: 0.0025 score: 595.68 eval: 1e-45 prob: 100 identity: 0.2976 startpos: 18<br>PWLKQLHGKVVVVKYGGNAMTDdrAFAADMAFLRNCGIHPVVVHGGGPQITAMLRRLGIEGDFKG---G--<br>FRVTTPEVLDVARMVLfqVGRELVNLINAHGPYAVGITGEDAQLFTAVRRSVT-----V-<br>DGVATDIGLVGDVDQVNTAAMLDLVAAGRIPVVSTLAPDADGVVHNINADTAAAAVAEALGAEKLLMLTDIDGLYTRWPDRDSLVSEIDTGTLA<br>QLLPTLELGMVPKVEACLRAVIGGVPSAHIIDGRVTHCVLVELfgTGTKVVRGEGHHHHH----<br>>3u6uA_304 weight: 0.0025 score: 26.78 eval: n/a prob: n/a identity: 0.4810 startpos: 1<br>---------MIVVKVGGAEGINYEAVAKDAASLWKEGVKLLLVHGGSAETNKVAEALGHPPRFLTHPV---<br>SRLTDRKTLEIFEMVYclVNKRLVELLQKEGANAIGLSGLDGRLFVGRRK-----------------<br>DYTGTVEEVNKALLDLLLQAGYLPVLTPPALSYENEAINTDGDQIAALLATLYGAEALVYLSNVPGLLA--<br>PDEASLVREIPVERIeeYLALAQGRMKRKVMGAVEAVKGGVKRVVFADGRVENPIRRALSGEGTVVR-------------<br>>2ap9A_106 weight: 0.0025 score: 267.4 eval: 7.9e-09 prob: n/a identity: 0.2941 startpos: 27<br>---------VVVVKYGGNAMtlRRAFAADMAFLRNCGIHPVVVHGGGPQITAMLRRLGIEGDFKG-----<br>GFRVTTPEVLDVARMVLfqVGRELVNLINAHGPYAVGITGEDAQLFTAVRRSVTVDGV------<br>ATDIGLVGDVDQVNTAAMLDLVAAGRIPVVSTLAPDADGVVHNINADTAAAAVAEALGAEKLLMLTDIDGLYTRWPDRDSLVSEIDTGTLAQLLP<br>TLELGMVPKVEACLRAVIGGVPSAHIIDGRVTHCVLVElaGTGTKVVRGEGHHH------<br>>2rd5A_107 weight: 0.0022 score: 266.9 eval: 8.2e-09 prob: n/a identity: 0.2768 startpos: 20<br>--------KTIVVKYGGAAMTskSSVVSDLVLLACVGLRPILVHGGGPDINRYLKQLNIPAEFRD-----<br>GLRVTDATTMEIVSMVLvkVNKNLVSLINAAGATAVGLSGHDGRLLTARPVPNSA-----------<br>QLGFVGEVARVDPSVLRPLVDYGYIPVIASVAADDSGQAYNINADTVAGELAAALGAEKLILLTDVAGILENKEDPSSLIKEIDIKGVKKMigKVAGG<br>MIPKVKCCIRSLAQGVKTASIIDGRRQHSLLHEImgAGTMITG------------<br>>2v5hA_110 weight: 0.0021 score: 254.5 eval: 2.1e-08 prob: n/a identity: 0.2734 startpos: 23<br>--------RTVVVKYGGAAMklKEAVMRDIVFLACVGMRPVVVHGGGPEINAWLGRVGIEPQFHN-----<br>GLRVTDADTMEVVEMVLvrVNKDIVSRINTTGGRAVGFCGTDGRLVLARPHDQEGI------------<br>GFVGEVNSVNSEVIEPLLERGYIPVISSVAADENGQSFNINADTVAGEIAAALNAEKLILLTDTRGILEDPKRPESLIPRLNIPQSRELigIVGGGMIPKV<br>DCCIRSLAQGVRAAHIIDGRIPHALLLEIfgIGTMIVGS-----------<br>>2jj4A_108 weight: 0.0021 score: 263.6 eval: 1.1e-08 prob: n/a identity: 0.2768 startpos: 23<br>--------RTVVVKYGGAAMklKEAVMRDIVFLACVGMRPVVVHGGGPEINAWLGRVGIEPQFHN-----<br>GLRVTDADTMEVVEMVLvrVNKDIVSRINTTGGRAVGFCGTDGRLVLARPHD------------<br>QEGIGFVGEVNSVNSEVIEPLLERGYIPVISSVAADENGQSFNINADTVAGEIAAALNAEKLILLTDTRGILEDPKRPESLIPRLNIPQSRELigIVGGGMI<br>PKVDCCIRSLAQGVRAAHIIDGRIPHALLLEiaGIGTMIVGS-----------<br>>2btyA_109 weight: 0.0021 score: 256.5 eval: 1.8e-08 prob: n/a identity: 0.2664 startpos: 22<br>--------KTFVIKFGGSAMkaKKAFIQDIILLKYTGIKPIIVHGGGPAISQMMKDLGIEPVFK------<br>GHRVTDEKTMEIVEMVLvkINKEIVMNLNLHGGRAVGICGKDSKLIVAEKET-----------<br>KHGDIGYVGKVKKVNPEILHALIENDYIPVIAPVGIGEDGHSYNINADTAAAEIAKSLMAEKLILLTDVDGVLKD----<br>GKLISTLTPDEAEELigTVTGGMIPKVECAVSAVRGGVGAVHIINGGLEHAILLEIfgIGTMIKELE----------<br>>2bufA_305 weight: 0.0018 score: 26.76 eval: n/a prob: n/a identity: 0.2734 startpos: 19<br>PYIRRFVGKTLVIKYGGNAMEskAGFARDVVLMKAVGINPVVVHGGGPQIGDLLKRLSIESHFIDG-----<br>MRVTDAATMDVVEMVLGgvNKDIVNLINRHGGSAIGLTGKDAELIRAKKL----TVTRQII----<br>DIGHVGEVTGVNVGLLNMLVKGDFIPVIAPIGVGSNGESYNINADLVAGKVAEALKAEKLMLLTNIAGLMD---<br>KQGQVLTGLSTEQVNELigTIYGGMLPKIRCALEAVQGGVTSAHIIDGRVPNAVLLEIfgVGTLISNRKRH--------<br>>3wwnA_306 weight: 0.0015 score: 26.64 eval: n/a prob: n/a identity: 0.5052 startpos: 1<br>---------<br>MIVVKVGGAEGINYEAVAKDAASLWKEGVKLLLVHGGSAETNKVAEALGHPPRFLTHPGGQVSRLTDRKTLEIFEMVYclVNKRLVELLQKEGANA<br>IGLSGLDGRLFVGRRKTAVKYVENG-<br>KVKVHRGDYTGTVEEVNKALLDLLLQAGYLPVLTPPALSYENEAINTDGDQIAALLATLYGAEALVYLSNVPGLLARYPDEASLVREIPVERIED----<br>PEGRMKRKVMGAVEAVRGGVKRVVFADARVENPIRRALSGEGTVVR-------------<br>>2ap9A_307 weight: 0.0014 score: 26.59 eval: n/a prob: n/a identity: 0.2976 startpos: 18<br>PWLKQLHGKVVVVKYGGNAMTdrRAFAADMAFLRNCGIHPVVVHGGGPQITAMLRRLGIEGDFKGG-----<br>FRVTTPEVLDVARMVLFgvGRELVNLINAHGPYAVGITGEDAQLFTAVRRSVT---VDGVAT--- |

| | | | | | | |
|---|---|---|---|---|---|---|
|  |  |  |  |  |  | DIGLVGDVDQVNTAAMLDLVAAGRIPVVSTLAPDADGVVHNINADTAAAAVAEALGAEKLLMLTDIDGLYTRWPDRDSLVSEIDTGTLAQLLPTL<br>ELGMVPKVEACLRAVIGGVPSAHIIDGRVTHCVLVELfgTGTKVVRGEGHHHHHH---<br>>3l86A_310 weight: 0.0013 score: 24.63 eval: n/a prob: n/a identity: 0.2215 startpos: 1<br>------MKDIIVIKIGGVASQQlgDFLSQIKNWQDAGKQLVIVHGGGFAINKLMEENQVPVKKING-----<br>LRVTSKDDMVLVSHALLdvGKNLQEKLRQAGVSCQQLKSDIKHVVAADYLDK-----------<br>DTYGYVGDVTHINKRVIEEFLENRQIPILASLGYSKEGDMLNINADYLATAVAVALAADKLILMTNVKGVL----<br>ENGAVLEKITSHQVQEKiaVITAGMIPKIESAAKTVAAGVGQVLIGDNLLT----------GTLITAD-----------<br>>2jj4A_309 weight: 0.0013 score: 26.15 eval: n/a prob: n/a identity: 0.2734 startpos: 15<br>PYLQQFAGRTVVVKYGGAAMKqkEAVMRDIVFLACVGMRPVVVHGGGPEINAWLGRVGIEPQFHNG-----<br>LRVTDADTMEVVEMVLvrVNKDIVSRINTTGGRAVGFCGTDGRLVLARPH--------DQE----<br>GIGFVGEVNSVNSEVIEPLLERGYIPVISSVAADENGQSFNINADTVAGEIAAALNAEKLILLTDTRGILEDPKRPESLIPRLNIPQSRELigIVGGGMIPK<br>VDCCIRSLAQGVRAAHIIDGRIPHALLLEIfgIGTMIVGS-----------<br>>2btyA_308 weight: 0.0013 score: 26.31 eval: n/a prob: n/a identity: 0.2699 startpos: 14<br>PYIKEFYGKTFVIKFGGSAMKqkKAFIQDIILLKYTGIKPIIVHGGGPAISQMMKDLGIEPVFKG------<br>HRVTDEKTMEIVEMVLVgiNKEIVMNLNLHGGRAVGICGKDSKLIVAEKE------TKHG-----<br>DIGYVGKVKKVNPEILHALIENDYIPVIAPVGIGEDGHSYNINADTAAAEIAKSLMAEKLILLTDVDGVL----<br>KDGKLISTLTPDEAEELigTVTGGMIPKVECAVSAVRGGVGAVHIINGGLEHAILLEIfgIGTMIKELEGR--------<br>>3u6uA_204 weight: 0.0011 score: 381.15 eval: 1e-45 prob: 100 identity: 0.4810 startpos: 1<br>---------MIVVKVGGAEGINYEAVAKDAASLWKEGVKLLLVHGGSAETNKVAEALGHPPRFLTHP---<br>VSRLTDRKTLEIFEMVYclVNKRLVELLQKEGANAIGLSGLDGRLFVGRRK----------------<br>DYTGTVEEVNKALLDLLLQAGYLPVLTPPALSYENEAINTDGDQIAALLATLYGAEALVYLSNVPGLL--<br>APDEASLVREIPVERIEDPeaLAQGRMKRKVMGAVEAVKGGVKRVVFADGRVENPIRRALSGEGTVVR------------<br>>2r8vA_205 weight: 0.0005 score: 508.18 eval: 1e-45 prob: 100 identity: 0.2422 startpos: 13<br>-YIRQMRGTTLVAGIDGRLLEglNKLAADIGLLSQLGIRLVLIHGAYHFLDRLAAAQGRTPHYCR---G--LRVTDETSLGQAQQFAGTVRSRFEAALCG-<br>----SSVPLVSGNFLTARPIGVI---------<br>DGTDMEYAGVIRKTDTAALRFQLDAGNIVWMPPLGHSYGGKTFNLDMVQAAASVAVSLQAEKLVYLTLSDGISRP---<br>DGTLAETLSAQEAQSLAEhaS-ETRRLISSAVAALEGGVHRVQILNGAADGSLLQElnGIGTSIAKEAFVSIRQAHS-<br>>2v5hA_206 weight: 0.0004 score: 558.43 eval: 1e-45 prob: 100 identity: 0.2768 startpos: 15<br>PYLQQFAGRTVVVKYGGAAMKQeeAVMRDIVFLACVGMRPVVVHGGGPEINAWLGRVGIEPQFHN---G--<br>LRVTDADTMEVVEMVlgRVNKDIVSRINTTGGRAVGFCGTDGRLVLARPHD-----------<br>QEGIGFVGEVNSVNSEVIEPLLERGYIPVISSVAADENGQSFNINADTVAGEIAAALNAEKLILLTDTRGILEDPKRPESLIPRLNIPQSRElqGIVGGGM<br>IPKVDCCIRSLAQGVRAAHIIDGRIPHALLLEIfgIGTMIVGS-----------<br>>3l86A_207 weight: 0.0003 score: 547.09 eval: 1e-45 prob: 100 identity: 0.2249 startpos: 1<br>------MKDIIVIKIGGVASQQLsdFLSQIKNWQDAGKQLVIVHGGGFAINKLMEENQVPVKKIN---G--<br>LRVTSKDDMVLVSHALllVGKNLQEKLRQAGVSCQQLKSDIKHVVAADYLD-----------<br>KDTYGYVGDVTHINKRVIEEFLENRQIPILASLGYSKEGDMLNINADYLATAVAVALAADKLILMTNVKGVLENG----<br>AVLEKITSHQVQEktAVITAGMIPKIESAAKTVAAGVGQVLIGDNLLT----------GTLITAD---------- |
